# Post-weaning social isolation impairs the orbitofrontal cortical circuit subserving contagious pain and prosocial behaviors

**DOI:** 10.64898/2025.12.16.694530

**Authors:** Yu Gong, Su-Shan Guo, Xue Chen, Zichen Zhang, Xing-Lei Song, Qin Jiang, Ying Li, Dehua Wu, Ting-Ting Zhang, Xuan Zhao, Fan Jiang, Hong Jiang, Ming-Gang Liu, Tian-Le Xu

**Affiliations:** Department of Anesthesiology, Songjiang Hospital and Songjiang Research Institute, Shanghai Key Laboratory of Emotions and Affective Disorders, Shanghai Jiao Tong University School of Medicine, Shanghai 201600, China; Department of Anatomy and Physiology, Shanghai Jiao Tong University School of Medicine, Shanghai 200025, China; Department of Anesthesiology, Shanghai Ninth People’s Hospital, Shanghai Jiao Tong University School of Medicine, Shanghai, 200011, China; Department of Anesthesiology, Shanghai Tenth People’s Hospital, School of Medicine, Tongji University, Shanghai 200072, China; Department of Developmental and Behavioral Pediatrics, Pediatric Translational Medicine Institute, National Children’s Medical Center, Shanghai Children’s Medical Center, Shanghai Jiao Tong University School of Medicine, Shanghai 200127, China; Institute of Mental Health and Drug Discovery, Oujiang Laboratory (Zhejiang Lab for Regenerative Medicine, Vision and Brain Health), Wenzhou, Zhejiang, China

**Keywords:** Contagious pain, allo-grooming/allo-licking, prosocial behaviors, social isolation, orbitofrontal cortex, *Grik3*

## Abstract

Empathic behaviors are sensitive to environmental factors like post-weaning social isolation (SI), yet the mechanisms by which SI affects empathy remain unclear. Here, we show that mice subjected to SI exhibit marked impairments in contagious pain and prosocial behaviors, including allo-grooming and allo-licking toward cagemates experiencing inflammatory pain. Mechanistically, we identify the glutamatergic projection from the ventromedial thalamic nucleus (VM) to the orbitofrontal cortex (OFC) as critical for these empathy-like responses. SI induces hypoexcitability of OFC glutamatergic neurons and attenuates excitatory synaptic transmission within the VM→OFC pathway. Remarkably, chemogenetic activation of OFC neurons or the VM→OFC projection restores empathic behaviors in SI mice. Furthermore, we uncover a molecular basis for SI-induced OFC hypoexcitability: the downregulation of *Grik3*, encoding a kainate-type glutamate receptor subunit. These findings reveal a previously uncharacterized thalamocortical mechanism through which early-life social deprivation disrupts empathic behaviors, offering insights into the underpinnings of social-affective dysfunction.

## Introduction

Empathy—the capacity to perceive, share, and respond to the emotional states of others—is essential for prosocial interaction, group cohesion, and cooperative behavior in complex environments^1–3^. Impairments in empathic processing are implicated in several neuropsychiatric conditions, including autism spectrum disorder, psychopathy, and schizophrenia^4,5^. Although once thought to be unique to humans, accumulating evidence demonstrates that rodents exhibit fundamental forms of empathy-like behaviors, such as observational fear^6–8^, emotional contagion of pain^9–12^, consolation^13–15^, and prosocial helping^16–18^. These models have enabled researchers to identify key neural substrates involved in empathy, such as anterior cingulate cortex and insular cortex, which are crucial for affective sharing and emotional contagion^7,11,12,19^. Other regions, such as the medial prefrontal cortex (mPFC), amygdala, and periaqueductal gray, contribute to social-affective learning, contextual processing, and behavioral output during empathic interactions^20–23^. Recent advances in optogenetics and *in vivo* imaging have begun to unravel the dynamic interactions among these regions during empathic responses. However, the precise circuit and molecular mechanisms that integrate social-affective cues and produce appropriate prosocial behaviors remain incompletely understood.

The orbitofrontal cortex (OFC), a ventral subdivision of the mPFC^24,25^, is recognized for its roles in value-based decision-making^26^, reversal learning^27,28^, reward valuation^29^, emotional regulation^30^, and sensory integration^31,32^. Given its anatomical connectivity and functional properties, the OFC is well-positioned to integrate sensory, emotional, and social information, suggesting a potential role in empathy-related processing. Human neuroimaging studies link OFC structural features to cognitive empathy^33^, and OFC dysfunction has been implicated in psychiatric disorders marked by social and emotional deficits^34–36^. Nevertheless, direct evidence for OFC involvement in rodent empathy-like behaviors is scarce. The specific circuits mediating this role remain unclear.

Social interactions during early life, especially in the juvenile period, are critical for the maturation of brain circuits involved in social-affective processing^37,38^. Juvenile experiences, such as play, communal nesting, and parental care, shape neural development and influence social recognition and emotional reactivity. The post-weaning period is particularly sensitive to environmental perturbation. Disruption during this developmental window, such as through social isolation (SI), has been shown to cause persistent changes in brain function and behavior ^39,40^, including heightened anxiety^41^, deficits in social interaction and recognition^42,43^, and altered reward-related processing^44^. However, the specific components of empathy most vulnerable to SI—and the underlying neural substrates—remain poorly characterized.

In this study, we identify a previously uncharacterized thalamocortical circuit and molecular mechanism through which early-life SI disrupts empathy-like behaviors. Using a post-weaning SI mouse model, we demonstrate that SI impairs both pain contagion and prosocial behaviors. We show that SI leads to hypoexcitability of OFC glutamatergic neurons and weakens excitatory synaptic input from the ventromedial thalamic nucleus (VM) to the OFC—a pathway not previously associated with empathic processing. Chemogenetic activation of either the OFC glutamatergic neurons or the VM→OFC projection rescues the behavioral deficits induced by SI. At the molecular level, we identify downregulation of *Grik3*, which encodes a kainate-type glutamate receptor subunit, as a key contributor to OFC hypoactivity and impaired empathic responses. These findings delineate a novel neural and molecular framework by which early-life social experiences shape empathy-related circuits and behavior.

## Results

### Post-weaning SI impairs contagious pain and prosocial behaviors

To investigate whether post-weaning SI disrupts contagious pain and prosocial behaviors in mice, we employed a single-housing protocol^45^ and established empathic social interaction paradigms^46^ (Fig. 1a). Male and female mice were socially isolated from weaning (3 weeks old) for 4 weeks, followed by 2 weeks of group housing (GH) to establish familiarity with cagemates. After a 3-day handling and habituation period, baseline mechanical pain thresholds were measured and found to be comparable between GH and SI groups (Extended Data Fig. 1a, c). On day four, bee venom (BV) was injected into the hind paw of a familiar demonstrator (painful cagemate), and observer mice (GH or SI) were allowed to interact freely with the injured demonstrator for 30 minutes. Following interaction, GH mice showed a significant reduction in mechanical pain thresholds, indicative of pain contagion (male: baseline *vs*. post: 0.70 ± 0.06 g *vs*. 0.16 ± 0.03 g; female: 0.77 ± 0.08 g *vs*. 0.37 ± 0.08 g), while SI mice exhibited no such changes (male: 0.81 ± 0.07 g *vs*. 0.75 ± 0.08 g; female: 0.57 ± 0.10 g *vs*. 0.56 ± 0.07 g; Fig. 1b, d). This deficit was further evident when comparing the percentage change from baseline (Fig. 1c, e) and the proportion of mice showing contagious pain (Extended Data Fig. 1b, d).

**Fig. 1.**
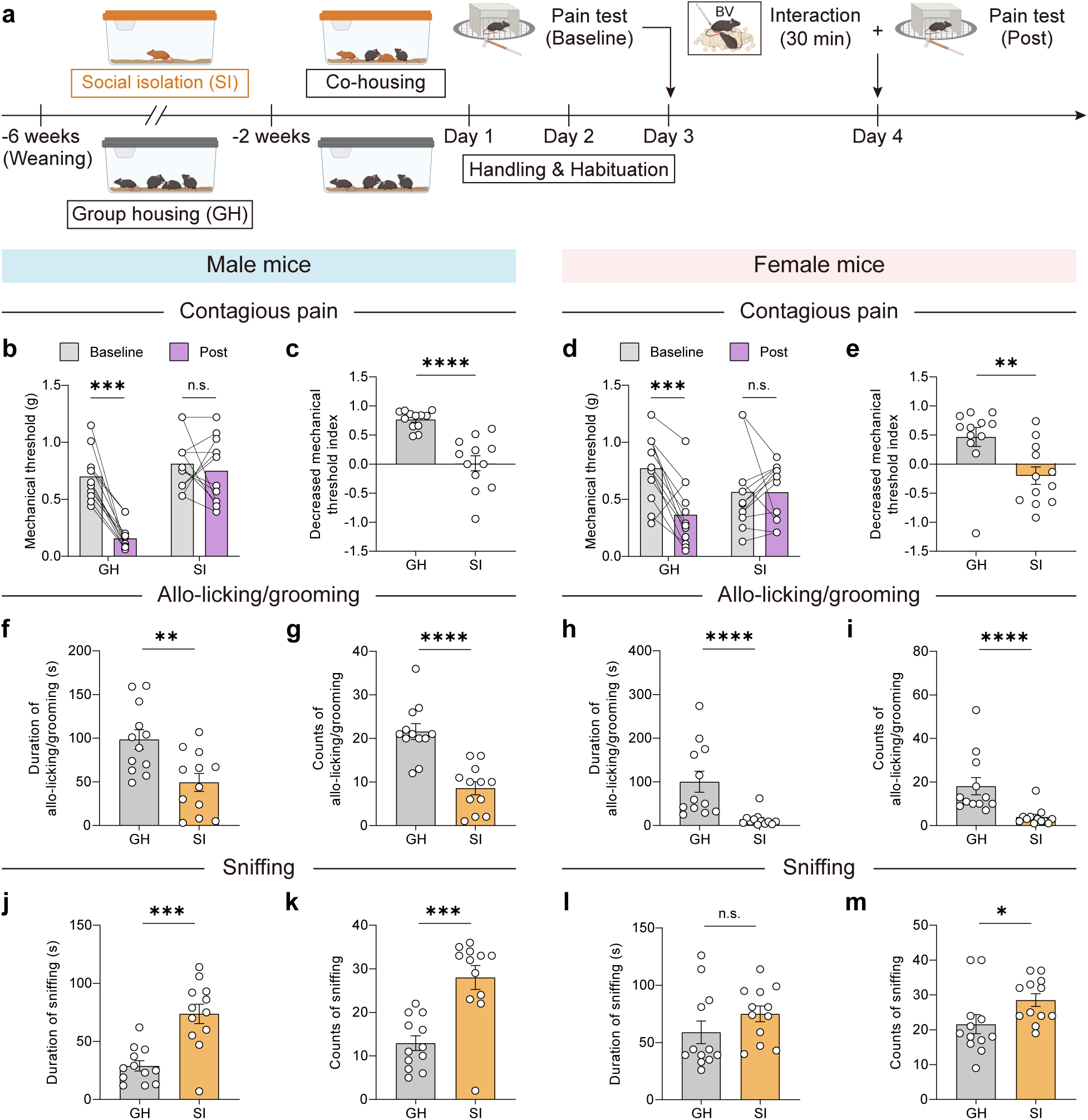
SI impairs contagious pain and prosocial behaviors in a sex-independent manner. **a,** Experimental timeline illustrating the effects of SI on empathic social interaction. GH mice served as controls. BV, bee venom. Created with BioRender.com. **b,** Changes in mechanical pain thresholds following a 30-minute interaction with a conspecific experiencing pain in male mice. *n* = 12 mice per group. GH: *p* = 0.0005, Wilcoxon matched-pairs signed rank test; SI: *p* = 0.5426, paired *t*-test. **c,** Percentage decrease in mechanical pain thresholds in male mice, calculated as (baseline − post)/baseline. *n* = 12 mice per group. *p* < 0.0001, unpaired *t*-test with Welch’s correction. **d,** Same as (**b**) but in female mice. *n* = 12 mice per group. GH: *p* = 0.0007, paired *t*-test; SI: *p* = 0.6772, Wilcoxon matched-pairs signed rank test. **e,** Same as (**c**) but in female mice. *n* = 12 mice per group. *p* = 0.0056, Mann-Whitney test. **f,** Total duration of allo-licking and allo-grooming behaviors during a 30-minute empathic social interaction in male mice. *n* = 12 mice per group. *p* = 0.0038, unpaired *t*-test. **g,** Number of bouts of allo-licking and allo-grooming behaviors in male mice. *n* = 12 mice per group. *p* < 0.0001, unpaired *t*-test. **h, i,** Same as (**f** and **g**), but in female mice. *n* = 12 mice per group. (**h**) *p* < 0.0001; (**i**) *p* < 0.0001; both Mann-Whitney tests. **j,** Total duration of sniffing behavior in male mice. *n* = 12 mice per group. *p* = 0.0002, unpaired *t*-test with Welch’s correction. **k,** Number of sniffing bouts in male mice. *n* = 12 mice per group. *p* = 0.0002, Mann-Whitney test. **l, m,** Same as (**j** and **k**), but in female mice. *n* = 12 mice per group. (**l**) *p* = 0.0908; (**m**) *p* = 0.0175; both Mann-Whitney tests. \**p* < 0.05, \*\**p* < 0.01, \*\*\**p* < 0.001, n.s., no significant difference. Data are expressed as mean ± SEM. See Supplementary Table 1 for full statistical details.

SI mice also displayed pronounced reductions in prosocial behaviors. Specifically, SI observers engaged in significantly less allo-licking and allo-grooming toward their BV-inflamed cagemates (male duration: GH *vs*. SI, 98.50 ± 11.20 s *vs*. 49.33 ± 10.24 s; counts: 21.58 ± 1.81 *vs*. 8.58 ± 1.51; Fig. 1f, g and Supplementary Videos 1 and 2), while simultaneously exhibiting increased sniffing behavior (duration: 28.92 ± 4.43 s *vs*. 73.67 ± 8.38 s; counts: 12.92 ± 1.70 *vs*. 28.00 ± 2.74; Fig. 1j, k). SI mice also displayed a delayed latency to initiate allo-licking/allo-grooming (Extended Data Fig. 1e) and an earlier latency to begin sniffing (Extended Data Fig. 1i). Bout analysis of these behaviors confirmed reduced duration per bout of allo-grooming/allo-licking and increased sniffing in SI mice (Extended Data Fig. 1f, j). These effects were similarly observed in female observers, though with some variation in behavioral parameters (Fig. 1h, i, l, m and Extended Data Fig. 1g, h, k, l). Self-grooming behavior, in contrast, remained largely unchanged in male mice (Extended Data Fig. 1m, n, q, r), while female SI mice showed a modest reduction (Extended Data Fig. 1o, p, s, t), suggesting that the SI effects were specific to other-directed social behaviors.

To better understand the temporal dynamics of these interactions, we analyzed behavioral counts and duration in 5-minute intervals over the 30-minute session in male observers. Prosocial and sniffing behaviors peaked during the initial 5 minutes (Extended Data Fig. 2a-f), indicating that the observers rapidly responded to the demonstrator’s distress. Across the session, SI mice consistently exhibited fewer prosocial behaviors and more sniffing compared to GH mice. Self-grooming was evenly distributed throughout the session and did not differ between groups (Extended Data Fig. 2g-i), further supporting the behavior-specific nature of the SI effects.

Given that adolescent social deprivation is associated with increased anxiety and impaired social function in adulthood^41,43,47^, we next tested whether the observed deficits stemmed from elevated anxiety or reduced sociability. Anxiety-like behavior was assessed using the open field test (OFT) and elevated plus maze (EPM), while social preference and discrimination were examined using the three-chamber test (Extended Data Fig. 3a, 4a). Neither male nor female SI mice displayed significant anxiety-like behavior or altered locomotion (Extended Data Fig. 3b-q). Both GH and SI mice showed robust preference for novel mice over empty cups and exhibited intact social discrimination between familiar and novel mice (Extended Data Fig. 4b-m), indicating normal sociability and social memory.

Acute social isolation (ASI) is associated with increased social need and loneliness states^48,49^. To determine whether ASI similarly disrupts empathic behaviors, we isolated adult male mice for 3 days following 2 weeks of co-housing (Extended Data Fig. 5a). After exposure to BV-inflamed cagemates, ASI-treated observers exhibited a clear decrease in mechanical pain thresholds, suggesting intact pain contagion (Extended Data Fig. 5b-e). Moreover, ASI had no effect on allo-licking/allo-grooming or sniffing behavior (Extended Data Fig. 5f-m).

Together, these findings demonstrate that post-weaning SI specifically impairs empathy-related behaviors in adult mice—manifested as deficits in contagious pain and prosocial interactions—without affecting anxiety levels, general sociability, or social memory.

### OFC glutamatergic neurons initiate prosocial behaviors

Having established that SI impairs empathy-related behaviors, we next investigated the underlying neural substrates. The OFC is classically known for mediating flexible decision-making by integrating risk, reward, and punishment signals^50,51^, but its role in social-emotional functions such as empathy remains poorly defined. To determine whether OFC neuronal activity contributes to empathic interactions and whether it is altered by SI, we performed c-Fos immunofluorescence staining in GH and SI male mice following observer-demonstrator interaction (Extended Data Fig. 6a). In GH mice, the OFC showed robust c-Fos activation, whereas this induction was significantly attenuated in SI mice (Extended Data Fig. 6b-d). The ventrolateral OFC (vlOFC) showed marked group differences (Extended Data Fig. 6b, c), while activation in the medial OFC (mOFC) remained low and unchanged (Extended Data Fig. 6b, d), identifying the vlOFC as the subregion most likely mediating SI-induced empathy deficits. All subsequent analyses, therefore, focused on the vlOFC.

To directly assess the role of OFC glutamatergic neurons in empathic behaviors, we conducted *in vivo* fiber photometry in GH male mice (Fig. 2a-c). We expressed GCaMP6m, a genetically encoded calcium indicator, in the OFC and implanted an optical fiber in the same region (Fig. 2b). Alignment of calcium signals with behavioral events revealed a consistent increase in OFC neuronal activity approximately 1 s prior to the initiation of both prosocial and sniffing behaviors (Fig. 2d-g and Supplementary Video 3). Notably, the calcium signal intensity during prosocial behaviors, quantified as either area under curve (AUC) or peak z-score, was significantly higher than that during sniffing events (Fig. 2f, h, I). These findings suggest that OFC activation not only precedes social behaviors but that the magnitude of activation may determine the behavioral outcome. Specifically, strong activation appears to drive affiliative behaviors (e.g., wound licking, grooming), whereas lower activation levels are associated with non-affiliative investigatory behaviors such as sniffing.

**Fig. 2.**
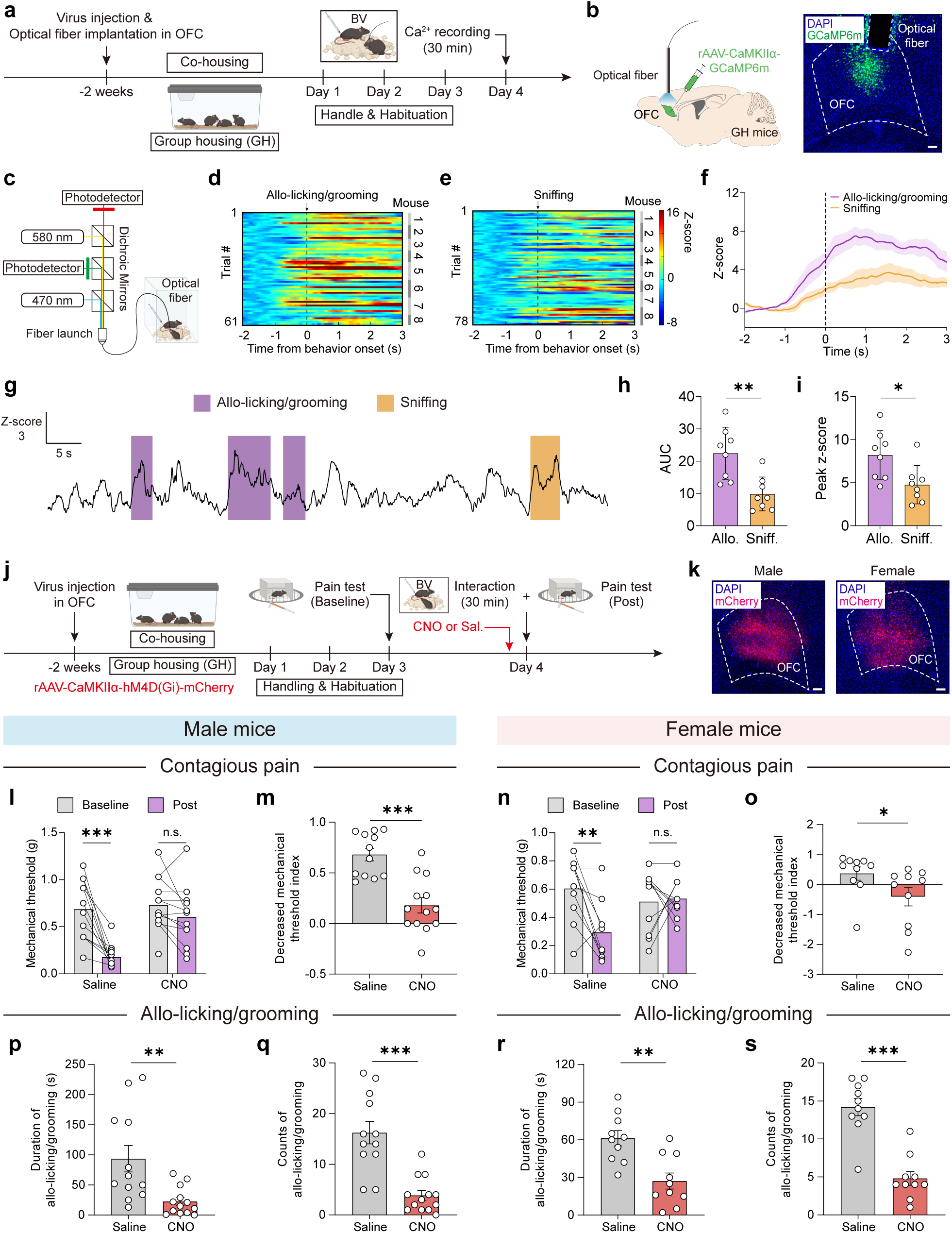
Glutamatergic neurons in the OFC mediate the initiation of prosocial behaviors. **a,** Timeline of fiber photometry recordings in OFC glutamatergic neurons in GH mice. **b,** Schematic (left) and representative image (right) showing virus injection and fiber placement in the OFC. Scale bar, 100 μm. **c,** Diagram of the two-color fiber photometry recording system. **d, e,** Heatmaps of Ca²⁺ signals aligned to the onset of allo-licking/grooming (**d**) and sniffing (**e**) behaviors. Each row shows one behavioral event. Color scale reflects Z-scored fluorescence. *n* = 8 mice. **f,** Per-event average traces of Z-scored Ca²⁺ signals during sniffing and prosocial behaviors. Solid lines represent the mean; shaded areas indicate SEM. *n* = 8 mice. **g,** Representative trace of Z-scored Ca²⁺ activity during sniffing and prosocial episodes. **h, i,** Quantification of Ca²⁺ activity changes as area under the curve (AUC; **h**) and peak Z-score (**i**). *n* = 8 mice per group. (**h**) *p* = 0.0024; (**i**) *p* = 0.0169; both unpaired *t*-tests. **j,** Timeline of chemogenetic inhibition of OFC glutamatergic neurons during empathic interaction in GH mice. **k,** Representative images of virus expression in the OFC of male (left) and female (right) mice. Scale bar, 100 μm. **l,** Mechanical pain threshold changes after 30-minute interaction with a painful demonstrator in male mice. Saline: *n* = 12; CNO: *n* = 13. Saline: *p* = 0.0005, Wilcoxon matched-pairs signed rank test; CNO: *p* = 0.0638, paired *t*-test. **m,** Percentage reduction in mechanical threshold. Saline: *n* = 12; CNO: *n* = 13. *p* = 0.0003, Mann-Whitney test. **n, o,** Same as (**l** and **m**), but in female mice. *n* = 10 mice per group. (**n**) Saline: *p* = 0.0054; CNO: *p* > 0.9999; two-way RM ANOVA with Bonferroni *post hoc* tests; (**o**) *p* = 0.0137, Mann-Whitney test. **p,** Total duration of prosocial behaviors in male mice. Saline: *n* = 12; CNO: *n* = 13. *p* = 0.0016, Mann-Whitney test. **q,** Number of prosocial behavior bouts. Saline: *n* = 12; CNO: *n* = 13. *p* < 0.0001, Mann-Whitney test. **r, s,** Same as (**p** and **q**), but in female mice. *n* = 10 mice per group. (**r**) *p* = 0.0011; (**s**) *p* < 0.0001; both unpaired *t*-tests. \**p* < 0.05, \*\**p* < 0.01, \*\*\**p* < 0.001, n.s., no significant difference. Data are shown as mean ± SEM. See Supplementary Table 1 for full statistical details.

To causally test this hypothesis, we chemogenetically suppressed OFC glutamatergic activity by bilaterally injecting AAV-CaMKIIα-hM4Di-mCherry into the OFC of GH mice and administering CNO 30 minutes prior to behavioral testing (Fig. 2j, k). Inhibition of OFC glutamatergic neurons significantly impaired contagious pain (male: Saline baseline *vs*. post, 0.69 ± 0.09 g *vs*. 0.18 ± 0.04 g; CNO: 0.73 ± 0.08 g *vs*. 0.60 ± 0.09 g; female: Saline, 0.61 ± 0.07 g *vs*. 0.29 ± 0.07 g; CNO: 0.51 ± 0.07 g *vs*. 0.60 ± 0.09 g; Fig. 2l-o and Extended Data Fig. 7a-d) and prosocial behaviors (male: Saline *vs*. CNO, duration: 93.50 ± 21.87 s *vs*. 22.46 ± 6.45 s; counts: 16.25 ± 2.21 *vs*. 3.85 ± 0.97; female: Saline *vs*. CNO, duration: 61.20 ± 6.05 s *vs*. 27.10 ± 6.36 s; counts: 14.20 ± 1.13 *vs*. 4.80 ± 0.90; Fig. 2p-s, Extended Data Fig. 7e-h and Supplementary Video 4). Meanwhile, sniffing behavior was significantly increased in both sexes (Extended Data Fig. 7i-p), with no changes in locomotor activity or anxiety-like behavior (Extended Data Fig. 8). These results confirm that OFC glutamatergic neuron activity is required for empathic behaviors.

### SI induces hypoexcitability of OFC glutamatergic neurons and impairs contagious pain and prosocial behaviors

To explore how SI alters OFC neuronal function, we recorded OFC glutamatergic activity in SI mice using fiber photometry (Extended Data Fig. 9a, b). Despite reduced prosocial and elevated sniffing behaviors in SI mice (Fig. 1f-m), the pre-behavior calcium rise was preserved, with similar AUC and peak Z-score signatures for each behavior type (Extended Data Fig. 9c-h). These findings suggest that while SI does not diminish evoked activity once initiated, it likely lowers the intrinsic excitability of OFC neurons, making them more difficult to reach the activation threshold required to trigger prosocial behaviors.

To examine possible alterations in the intrinsic excitability of OFC glutamatergic neurons, we performed *ex vivo* whole-cell patch-clamp recordings in OFC slices from GH and SI mice (Fig. 3a). SI mice exhibited significantly reduced action potential firing in response to depolarizing current steps (150 pA: GH *vs*. SI, 20.46 ± 1.65 Hz *vs*. 10.20 ± 1.84 Hz; Fig. 3b, c). Additionally, SI neurons showed hyperpolarized resting membrane potentials, elevated AP thresholds, increased rheobase, and prolonged rise tau, with no significant differences in other passive or active properties (Extended Data Fig. 10a-g). We further found a significant reduction in spontaneous excitatory postsynaptic currents (sEPSCs) frequency (GH *vs*. SI, 13.25 ± 2.83 Hz *vs*. 6.64 ± 1.22 Hz; Fig. 3d, f), while sEPSCs amplitude remained unchanged (Fig. 3d, e). Spontaneous inhibitory postsynaptic currents (sIPSCs) were unaffected in both frequency and amplitude (Fig. 3g-i), suggesting a selective loss of excitatory synaptic input and a disruption of excitation/inhibition balance (Extended Data Fig. 10h, i). Collectively, these data indicate that SI reduces both intrinsic excitability and excitatory synaptic drive onto OFC glutamatergic neurons.

**Fig. 3.**
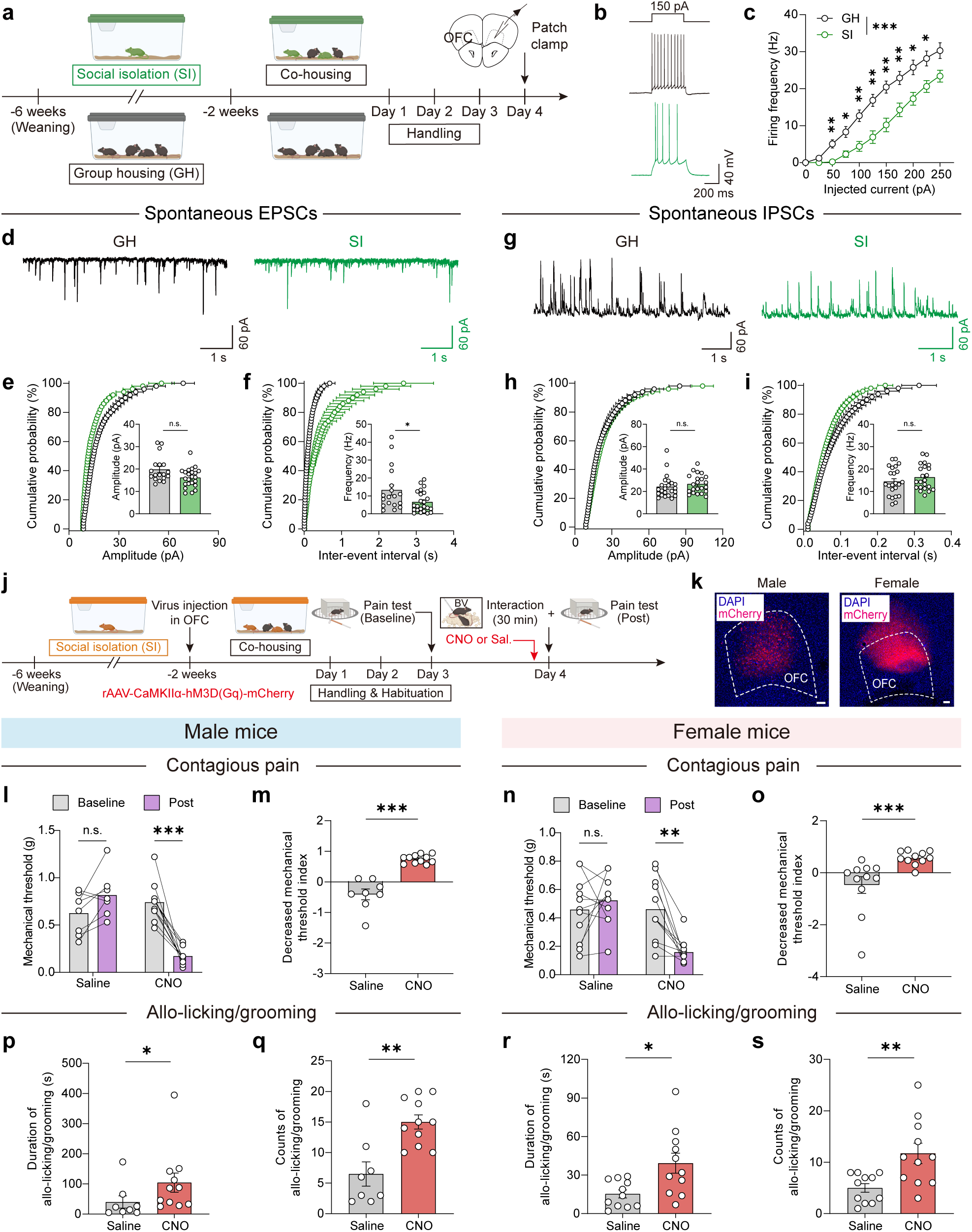
SI reduces excitability of OFC glutamatergic neurons, impairing contagious pain and prosocial behaviors. **a,** Experimental timeline for whole-cell patch-clamp recordings from OFC glutamatergic neurons in GH and SI mice. **b,** Representative traces of action potentials in response to 150 pA current injections from GH (black) and SI (green) mice. **c,** Firing frequency across current injection steps. GH: *n* = 26 neurons from 8 mice; SI: *n* = 20 neurons from 3 mice. GH vs. SI: *p* < 0.0001. *Post hoc p*-values: 0 pA: *p* > 0.9999; 25 pA: *p* = 0.0837; 50 pA: *p* = 0.0023; 75 pA: *p* = 0.0152; 100 pA: *p* = 0.0018; 125 pA: *p* = 0.0022; 150 pA: *p* = 0.0022; 175 pA: *p* = 0.0054; 200 pA: *p* = 0.0152; 225 pA: *p* = 0.0345; 250 pA: *p* = 0.0837. Two-way RM ANOVA with Mann-Whitney *post hoc* tests. **d,** Representative traces of spontaneous excitatory postsynaptic currents (sEPSCs) in OFC neurons from GH and SI mice. **e,** Cumulative probability and mean amplitude of sEPSCs. GH: *n* = 18 neurons from 6 mice; SI: *n* = 23 neurons from 4 mice. *p* = 0.0598, Mann-Whitney test. **f,** Cumulative probability and mean frequency of sEPSCs. GH: *n* = 18 neurons from 6 mice; SI: *n* = 23 neurons from 4 mice. *p* = 0.0436, Mann-Whitney test. **g-i,** Same as (**d–f**), but for spontaneous inhibitory postsynaptic currents (sIPSCs). GH: *n* = 23 neurons from 6 mice; SI: *n* = 22 neurons from 4 mice. (**h**) *p* = 0.1712, Mann-Whitney test; (**i**) *p* = 0.2872, unpaired *t*-test. **j,** Timeline of chemogenetic activation of OFC glutamatergic neurons during empathic interaction in SI mice. **k,** Representative images showing virus expression in OFC of male (left) and female (right) mice. Scale bar, 100 μm. **l,** Changes in mechanical pain thresholds in male mice. Saline: *n* = 8; CNO: *n* = 11. Saline: *p* = 0.0772; CNO: *p* < 0.0001; two-way RM ANOVA with Bonferroni *post hoc* tests. **m,** Percentage reduction in mechanical threshold. *p* = 0.0002, unpaired *t*-test with Welch’s correction. **n, o,** Same as (**l** and **m**), but in female mice. *n* = 11 mice per group. (**n**) Saline: *p* = 0.4246, paired *t*-test; CNO: *p* = 0.002, Wilcoxon matched-pairs signed rank test. (**o**) *p* = 0.0001, Mann-Whitney test. **p,** Duration of prosocial behaviors in male SI mice. Saline: *n* = 8; CNO: *n* = 11. *p* = 0.0198, Mann-Whitney test. **q,** Number of prosocial behavior bouts. Saline: *n* = 8; CNO: *n* = 11. *p* = 0.001, unpaired *t*-test. **r, s,** Same as (**p** and **q**), but in female SI mice. *n* = 11 mice per group. (**r**) *p* = 0.0136, unpaired *t*-test with Welch’s correction; (**s**) *p* = 0.0096, Mann-Whitney test. \**p* < 0.05, \*\**p* < 0.01, \*\*\**p* < 0.001, n.s., no significant difference. Data are shown as mean ± SEM. See Supplementary Table 1 for full statistical details.

Given the established necessity of OFC neuronal activity for empathic behaviors (Fig. 2j-s), we next tested whether reactivating OFC neurons could rescue the impaired empathic behaviors in SI mice. To this end, we bilaterally injected AAV-CaMKIIα-hM3Dq-mCherry into the OFC and administered CNO to chemogenetically enhance neuronal excitability (Fig. 3j, k). Activation of OFC glutamatergic neurons rescued contagious pain responses (male: Saline *vs*. CNO, baseline *vs*. post, 0.63 ± 0.08 g *vs*. 0.82 ± 0.08 g; 0.74 ± 0.06 g *vs*. 0.17 ± 0.03 g; female: 0.46 ± 0.07 g *vs*. 0.52 ± 0.05 g; 0.46 ± 0.07 g *vs*. 0.16 ± 0.03 g; Fig. 3l-o and Extended Data Fig. 11a-d) and significantly increased prosocial behaviors (male: Saline *vs*. CNO, duration: 40.00 ± 20.27 s *vs*. 104.20 ± 32.14 s; counts: 6.50 ± 1.97 *vs*. 15.00 ± 1.15; female: duration: 15.36 ± 3.12 s *vs*. 39.27 ± 7.79 s; counts: 5.00 ± 0.84 *vs*. 11.73 ± 1.98; Fig. 3p-s, Extended Data Fig. 11e-h and Supplementary Video 4). CNO treatment also significantly reduced sniffing behavior in male—but not female—SI mice (Extended Data Fig.11i-p).

In summary, these findings demonstrate that SI impairs empathy-related behaviors by reducing the excitability and excitatory synaptic input of OFC glutamatergic neurons. Restoring OFC excitability is sufficient to rescue both contagious pain and prosocial behaviors, highlighting the central role of OFC neuronal dysfunction in social-affective impairments induced by social deprivation.

### VM→OFC glutamatergic projection mediates contagious pain and prosocial behaviors

Given that OFC glutamatergic neurons receive reduced excitatory input following SI (Fig. 3d-f), we hypothesized that upstream afferents play a critical role in initiating empathic responses. To identify presynaptic sources, we employed the TRAP2 system^11,52^ to genetically tag OFC neurons activated during empathic social interaction, followed by monosynaptic retrograde tracing using modified rabies virus (Extended Data Fig. 12a). TRAP2 mice received injections of AAV–DIO–RVG and AAV–DIO–TVA–EGFP into the OFC, followed by 2 weeks of co-housing for activity-dependent labeling. After habituation, we administered 4-hydroxytamoxifen (4–OHT; 50 mg/kg, i.p.) 30 minutes before a 2-hour empathic interaction to drive Cre-mediated recombination in active OFC neurons. Three weeks later, EnvA-pseudotyped, glycoprotein-deleted rabies virus encoding DsRed (RV–EnvA–ΔG–DsRed) was injected into the same OFC site. One week post-injection, co-labeled starter cells (GFP^+^/DsRed^+^) confirmed selective targeting of socially activated OFC neurons (Extended Data Fig. 12b, c). Quantitative brain-wide mapping revealed that the ventromedial thalamus (VM) was a prominent source of monosynaptic input to social interaction-TRAPed OFC neurons (Extended Data Fig. 12d, e), implicating VM as a key upstream regulator of empathic processing.

To determine the target cell type within the OFC, we injected the anterograde tracer AAV2/1-EF1α-Flp into the VM and co-injected AAV-nEF1α-fDIO-mCherry and AAV-EF1α-DIO-H2B-EGFP into the OFC of CaMKIIα-Cre mice (Extended Data Fig. 12f). After 3–4 weeks, we observed extensive mCherry expression in OFC neurons, with substantial colocalization with EGFP (46.1% ± 3.5%, *n* = 4), indicating that VM projections primarily target OFC glutamatergic neurons (Extended Data Fig. 12g, h).

To assess real-time activity in this circuit during empathic behaviors, we performed fiber photometry in GH mice. AAV-CaMKIIα-GCaMP6m was injected into the VM, and an optical fiber was implanted over the OFC to record calcium signals from VM axon terminals during specific social behaviors (Fig.4a, b). VM→OFC terminal activity increased prior to the initiation of both allo-licking/allo-grooming and sniffing behaviors (Fig. 4c-f), with higher signal amplitude associated with prosocial compared to sniffing behaviors (Fig. 4g, h). These data suggest that enhanced excitatory input from VM facilitates prosocial engagement.

**Fig. 4.**
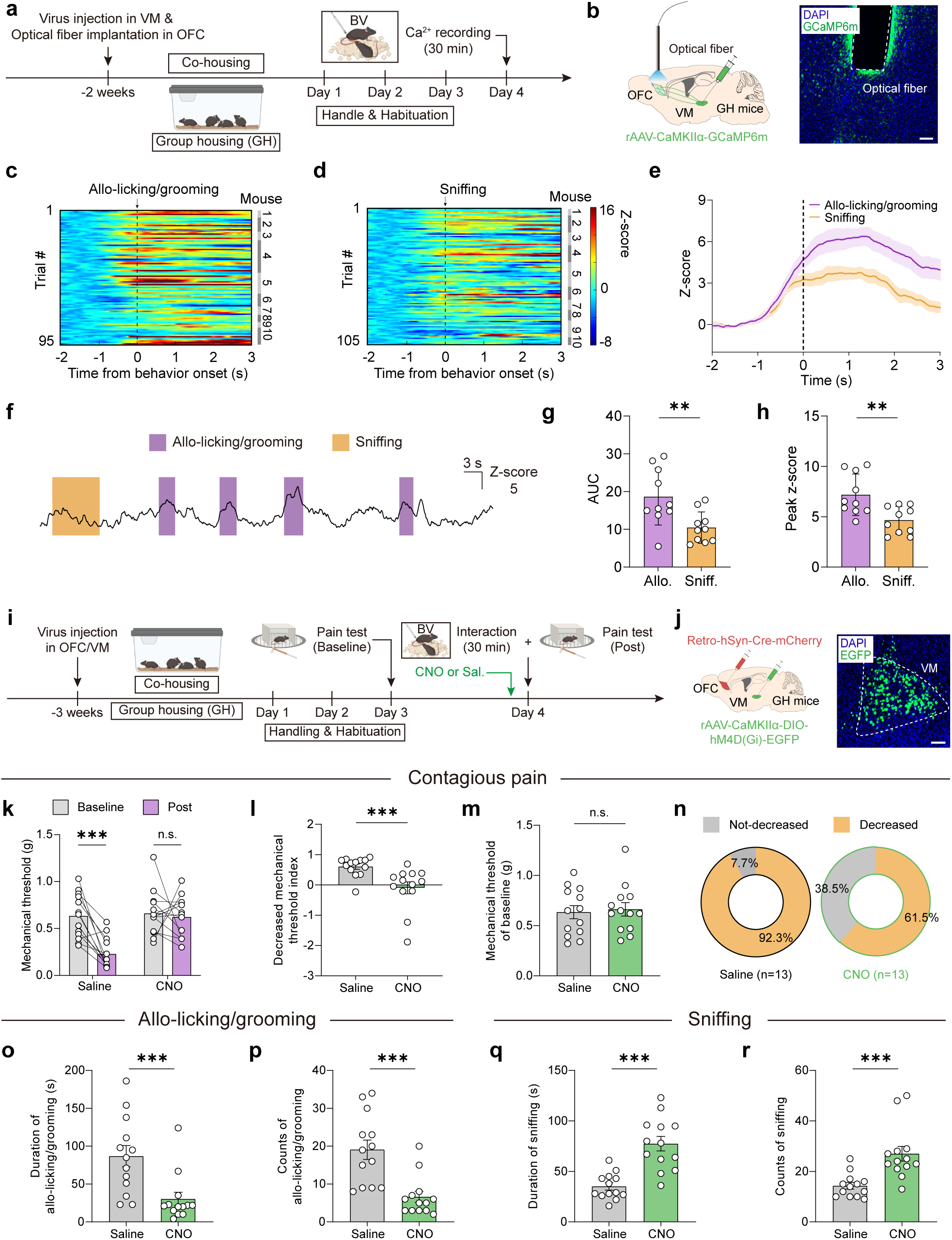
Glutamatergic projections from the ventromedial thalamus (VM) to the OFC mediate contagious pain and prosocial behaviors. **a,** Experimental timeline for fiber photometry recording of VM glutamatergic axon terminals within the OFC of GH mice. **b,** Schematic diagram (left) and representative image (right) for virus injection sites and optical fiber implantation in the VM and OFC. Scale bar, 100 μm. **c, d,** Heatmaps of GCaMP6m calcium signals aligned to the onset of allo-licking/grooming (**c**) or sniffing (**d**) behaviors. *n* = 10 mice. **e,** Event-aligned Z-scored calcium signals during episodes of sniffing and prosocial behaviors. *n* = 10 mice. **f,** Representative trace of Z-scored Ca²⁺ activity during sniffing and prosocial episodes. **g, h,** Quantification of GCaMP6m responses as AUC (**g**) or peak Z-score (**h**). *n* = 10 mice per group. (**g**) *p* = 0.0073; (**h**) *p* = 0.0046; both unpaired *t*-tests. (**i**) Experimental timeline for chemogenetic inhibition of VM→OFC projections during empathic social interaction in GH male mice. **j,** Schematic diagram (left) and representative image (right) for virus injection sites in the VM and OFC. Scale bar, 100 μm. **k,** Mechanical pain threshold changes. *n* = 13 mice per group. Saline: *p* = 0.0005, Wilcoxon matched-pairs signed rank test; CNO: *p* = 0.7006, paired *t*-test. **l,** Percentage decrease in mechanical pain thresholds. *n* = 13 mice per group. *p* = 0.0008, Mann-Whitney test. **m,** Baseline mechanical thresholds prior to social interaction. *n* = 13 mice per group. *p* = 0.7643, unpaired *t*-test. **n,** Proportion of mice exhibiting or not exhibiting contagious pain. **o,** Total duration of prosocial behaviors during a 30-minute social interaction. *n* = 13 mice per group. *p* = 0.0004, Mann-Whitney test. **p,** Number of prosocial behavior bouts. *n* = 13 mice per group. *p* < 0.0001, Mann-Whitney test. **q,** Total duration of sniffing behavior. *n* = 13 mice per group. *p* < 0.0001, unpaired *t*-test with Welch’s correction. **r,** Number of sniffing bouts. *n* = 13 mice per group. *p* = 0.0001, Mann-Whitney test. \*\**p* < 0.01, \*\*\**p* < 0.001, n.s., no significant difference. Data represent mean ± SEM. See Supplementary Table 1 for full statistical details.

Next, to test the necessity of this projection, we chemogenetically silenced the VM→OFC pathway by injecting AAV-Retro-hSyn-Cre into the OFC and Cre-dependent AAV-CaMKIIα-DIO-hM4Di-EGFP into the VM of GH mice. (Fig. 4i, j). Upon CNO administration, mice displayed impaired contagious pain (baseline vs. post: Saline: 0.63 ± 0.07 g *vs*. 0.23 ± 0.05 g; CNO: 0.66 ± 0.07 g *vs*. 0.62 ± 0.06 g; Fig. 4k-n), along with marked reductions in both counts and duration of prosocial behaviors (Saline *vs*. CNO, duration: 86.85 ± 14.08 s *vs*. 30.23 ± 8.91 s; counts: 19.08 ± 2.56 *vs*. 6.62 ± 1.46; Fig. 4o, p and Extended Data Fig. 13a, b), and a concomitant increase in sniffing behavior (Fig. 4q, r and Extended Data Fig. 13c, d). Locomotor activity and anxiety-like behavior remained unaffected (Extended Data Fig. 13e-k). Finally, given the substantial inputs to the OFC from the thalamic nucleus submedius (Sub) (Extended Data Fig. 12d, e) and previously-reported role of Sub in nociceptive modulation^53^, we also tested the possible involvement of this pathway. However, chemogenetic inhibition of the Sub→OFC circuit had no effect on contagious pain or social behaviors (Extended Data Fig. 14), confirming the specificity of VM→OFC projections in mediating empathic responses.

Collectively, these results establish the VM→OFC glutamatergic projection as a critical circuit mediating empathic behaviors. This pathway provides the excitatory drive necessary for OFC activation to reach the threshold required for initiating prosocial actions.

### Chemogenetic activation of the VM→OFC projection restores SI-induced deficits in contagious pain and prosocial behaviors

To investigate whether SI alters synaptic strength within the VM→OFC circuit, we expressed AAV-CaMKIIα-ChR2-mCherry in VM neurons of GH and SI mice, and prepared acute OFC slices after 3 weeks (Fig. 5a). Optical stimulation (473 nm, 5 ms pulses) under aCSF, TTX (0.5 µM), and TTX + 4-AP (100 µM) confirmed monosynaptic glutamatergic connectivity, with no direct inhibitory inputs (Fig. 5b, c and Extended Data Fig. 15a, b). Voltage-clamp recordings at –70 mV and 0 mV during paired-pulse stimulation (25–500 ms intervals) revealed increased excitatory paired-pulse ratios (ePPRs) in SI mice (GH *vs*. SI: 0.36 ± 0.04 *vs*. 0.55 ± 0.05 at 200 ms; Fig. 5d, e), indicating reduced presynaptic release probability. Inhibitory PPRs (iPPRs) were unaffected (Extended Data Fig. 15c, d). Furthermore, SI led to a significant reduction in AMPA/NMDA ratio (GH *vs*. SI: 5.40 ± 0.80 *vs*. 3.25 ± 0.46; Fig. 5f, g), suggesting weakened postsynaptic efficacy. Together, these findings demonstrate diminished excitatory synaptic transmission in the VM→OFC circuit following SI, thus providing a mechanistic link to impaired empathic behaviors.

**Fig. 5.**
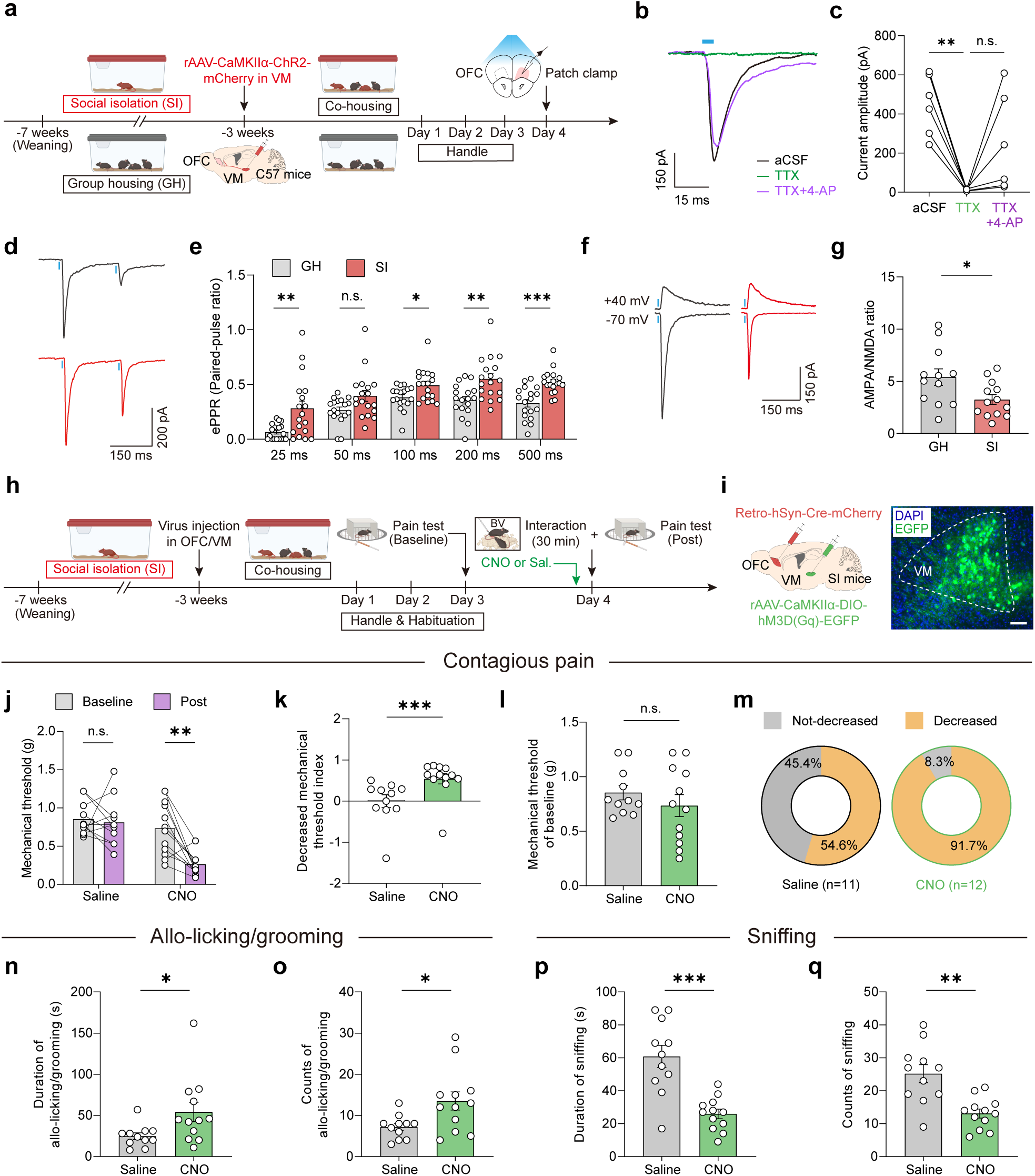
Chemogenetic activation of VM→OFC projections rescues SI-induced deficits in contagious pain and prosocial behaviors. **a,** Experimental timeline for whole-cell patch-clamp recordings of light-evoked EPSCs in OFC glutamatergic neurons. **b,** Representative traces of light-evoked EPSCs in artificial cerebrospinal fluid (aCSF), tetrodotoxin (TTX), or TTX + 4-aminopyridine (4-AP). **c,** Quantification of EPSC amplitudes. *n* = 6 neurons from 3 mice. aCSF vs. TTX: *p* = 0.0028; TTX vs. TTX+4-AP: *p* = 0.2124; one-way RM ANOVA with Bonferroni *post hoc* tests. **d, e,** Example traces (**d**) and summary (**e**) of excitatory paired-pulse ratio (ePPR) of light-evoked EPSCs. GH: *n* = 18 neurons from 5 mice; SI: *n* = 18 neurons from 4 mice. 25 ms: *p* = 0.0051, Mann-Whitney test; 50 ms: *p* = 0.0778, Mann-Whitney test; 100 ms: *p* = 0.0224, Mann-Whitney test; 200 ms: *p* = 0.0019, unpaired *t-*test; 500 ms: *p* < 0.0001, unpaired *t*-test. **f, g,** Representative traces (**f**) and quantification (**g**) of AMPA/NMDA ratio. GH: *n* = 12 neurons from 4 mice; SI: *n* = 13 neurons from 5 mice. *p* = 0.0262, unpaired *t*-test. **h,** Timeline for chemogenetic activation of VM→OFC projections during empathic social interaction in SI male mice. **i,** Schematic diagram (left) and representative image (right) for virus injection sites in the VM and OFC. Scale bar, 100 μm. **j,** Mechanical pain threshold changes. Saline: *n* = 11; CNO: *n* = 12. Saline: *p* = 0.7086, paired *t*-test; CNO: *p* = 0.0024, Wilcoxon matched-pairs signed rank test. **k,** Percentage reduction in pain threshold. Saline: *n* = 11; CNO: *n* = 12. *p* = 0.0003, Mann-Whitney test. **l,** Baseline mechanical thresholds. Saline: *n* = 11; CNO: *n* = 12. *p* = 0.3432, unpaired *t*-test. **m,** Proportion of mice with or without contagious pain. **n,** Total duration of prosocial behaviors. Saline: *n* = 11; CNO: *n* = 12. *p* = 0.0176, Mann-Whitney test. **o,** Number of prosocial behavior bouts. Saline: *n* = 11; CNO: *n* = 12. *p* = 0.0188, unpaired *t*-test with Welch’s correction. **p,** Total duration of sniffing behavior. Saline: *n* = 11; CNO: *n* = 12. *p* = 0.0003, unpaired *t*-test with Welch’s correction. **q,** Number of sniffing bouts. Saline: *n* = 11; CNO: *n* = 12. *p* = 0.0012, unpaired *t*-test with Welch’s correction. \**p* < 0.05, \*\**p* < 0.01, \*\*\**p* < 0.001, n.s., no significant difference. Data represent mean ± SEM. See Supplementary Table 1 for full statistical details.

To clarify the source of indirect inhibition onto OFC glutamatergic neurons, we sequentially applied PTX (100 μM), then CNQX (10 μM) and APV (50 μM), and vice versa. PTX reduced light-evoked EPSCs, while subsequent CNQX/APV abolished them entirely, confirming the glutamatergic identity of VM→OFC connection (Extended Data Fig. 15f, g). Reversing the order confirmed the absence of monosynaptic GABAergic inputs (Extended Data Fig. 15h, i), indicating that VM terminals likely engage local interneurons to generate feed-forward inhibition (Extended Data Fig. 15e), potentially serving to tune OFC activity for optimal empathic responses.

Finally, to test whether reactivating this pathway could reverse SI-induced behavioral deficits, we injected AAV-Retro-hSyn-Cre into the OFC and Cre-dependent AAV-CaMKIIα-DIO-hM3Dq-EGFP into the VM of SI mice (Fig. 5h, i). CNO administration restored mechanical pain contagion (baseline *vs*. post: Saline: 0.85 ± 0.06 g *vs*. 0.81 ± 0.09 g; CNO: 0.74 ± 0.10 g *vs*. 0.27 ± 0.05 g; Fig. 5j-m), increased the counts and duration of prosocial behaviors (Saline vs. CNO, duration: 24.82 ± 4.02 s *vs*. 54.17 ± 11.76 s; counts: 7.18 ± 0.87 *vs*. 13.50 ± 2.22; Fig. 5n, o), and suppressed excessive sniffing (Fig. 5p, q). These findings confirm the causal role of the VM→OFC pathway in mediating SI-evoked empathic impairment.

### SI-induced downregulation of *Grik3* leads to orbitofrontal hypoexcitability and empathic deficits

To elucidate the molecular mechanisms by which SI induces orbitofrontal neuronal hypoexcitability and consequent empathic behavioral deficits, we performed droplet-based single-nucleus RNA sequencing (snRNA-seq) on OFC tissues from GH and SI mice (Fig. 6a). From three biological replicates per group, we recovered 41,843 nuclei from GH and 38,913 nuclei from SI samples (Extended Data Fig. 16a, b), each exhibiting approximately 4,000 unique molecular identifiers (UMIs) per nucleus (Extended Data Fig. 16d, e). Principal Component Analysis (PCA) followed by t-distributed Stochastic Neighbor Embedding (t-SNE) and unsupervised clustering identified 10 distinct cell populations across all samples (Fig. 6b). These were annotated based on the expression profiles of 40 canonical cell-type markers (Extended Data Fig. 17), showing comparable distributions across GH and SI groups (Extended Data Fig. 16c), but displaying differences in UMI counts (Extended Data Fig. 16f), potentially reflecting transcriptional complexity.

**Fig. 6.**
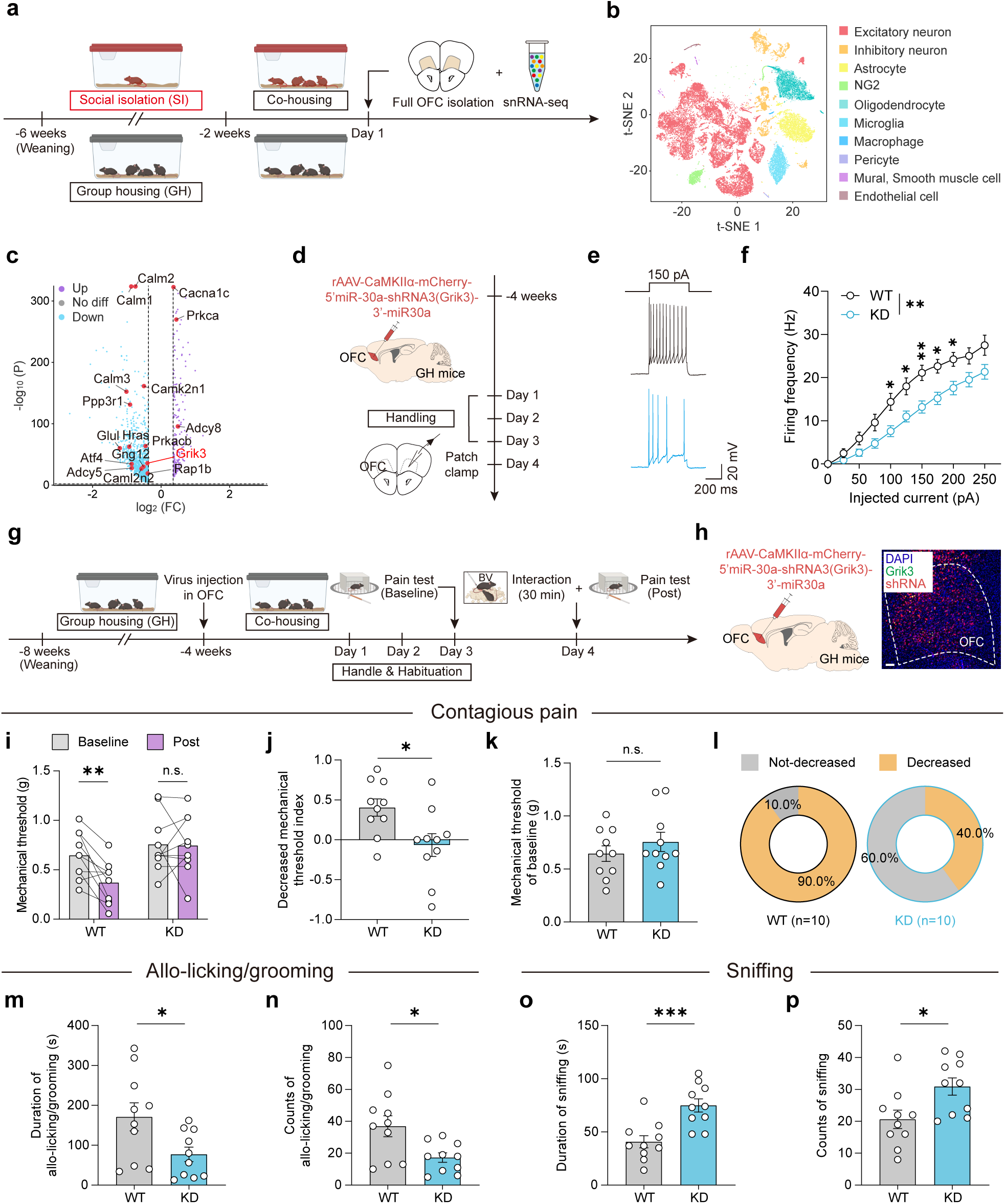
SI reduces *Grik3* expression in OFC glutamatergic neurons, leading to hypoexcitability and impaired contagious pain and prosocial behaviors. **a**, Timeline for single-nucleus RNA sequencing (snRNA-seq) of OFC samples from GH and SI mice. **b,** t-SNE visualization of ten major cell types identified from OFC samples of GH and SI mice. **c,** Volcano plot showing differentially expressed genes in OFC glutamatergic neurons between GH and SI groups. **d,** Timeline for whole-cell patch-clamp recordings following *Grik3* knockdown (KD) in OFC glutamatergic neurons. **e, f,** Representative traces (**e**) and summary data (**f**) of action potentials in response to stepwise current injections in WT (black) and *Grik3* KD (blue) neurons. WT: *n* = 25 neurons from 4 mice; KD: *n* = 30 neurons from 4 mice. Overall: *p* = 0.0024; 0 pA: *p* > 0.9999; 25 pA: *p* = 0.2936; 50 pA: *p* = 0.1284; 75 pA: *p* = 0.1003; 100 pA: *p* = 0.0254; 125 pA: *p* = 0.0254; 150 pA: *p* = 0.0095; 175 pA: *p* = 0.0137; 200 pA: *p* = 0.0335; 225 pA: *p* = 0.1306; 250 pA: *p* = 0.1637. Two-way RM ANOVA with Mann-Whitney *post hoc* tests. **g,** Experimental timeline for *Grik3* KD prior to social interaction in GH male mice. **h,** Schematic diagram (left) and representative image (right) for virus injection sites in the OFC. Scale bar, 100 μm. **i,** Changes in mechanical pain thresholds. *n* = 10 mice per group. WT: *p* = 0.0081; KD: *p* > 0.9999; two-way RM ANOVA with Bonferroni *post hoc* tests. **j,** Percentage reduction in mechanical threshold. *n* = 10 mice per group. *p* = 0.0171, unpaired *t*-test. **k,** Baseline mechanical thresholds. *n* = 10 mice per group. *p* = 0.3559, unpaired *t*-test. **l,** Proportion of mice with or without contagious pain. **m,** Total duration of prosocial behaviors. *n* = 10 mice per group. *p* = 0.0293, unpaired *t*-test. **n,** Number of prosocial behavior bouts. *n* = 10 mice per group. *p* = 0.0178, unpaired *t*-test with Welch’s correction. **o,** Total duration of sniffing behavior. *n* = 10 mice per group. *p* = 0.0008, unpaired *t*-test. **p,** Number of sniffing bouts. *n* = 10 mice per group. *p* = 0.0177, unpaired *t*-test. \**p* < 0.05, \*\**p* < 0.01, \*\*\**p* < 0.001, n.s., no significant difference. Data are presented as mean ± SEM. See Supplementary Table 1 for full statistical details.

Focusing on excitatory neuron clusters, we identified differentially expressed genes (DEGs) between SI and GH conditions (Fig. 6c). Kyoto Encyclopedia of Genes and Genomes (KEGG) pathway analysis revealed significant downregulation of genes involved in long-term potentiation and glutamatergic synaptic transmission, findings further corroborated by gene set enrichment analysis (GSEA) (Extended Data Fig. 18a, b). Notably, the expression of *Grik3*, encoding a kainate receptor subunit mediating Na⁺/Ca²⁺ influx^54,55^, was markedly decreased in excitatory neurons from SI mice (Fig. 6c and Extended Data Fig. 18c-e). GRIK3 has been implicated in the pathophysiology of schizophrenia^56^ and in modulating pain-like behaviors in murine models^57^.

To directly assess the role of *Grik3* in regulating neuronal excitability, we knocked down *Grik3* specifically in OFC glutamatergic neurons of GH mice (Fig. 6d). Whole-cell patch-clamp recordings revealed significantly reduced action potential firing in *Grik3* knockdown (KD) neurons compared to wild-type (WT) controls (firing rate: WT *vs*. KD: 21.12 ± 1.77 Hz *vs*. 13.20 ± 1.42 Hz at 150 pA; Fig. 6e, f), without alterations in other intrinsic membrane properties (Extended Data Fig. 19a-e). Behaviorally, *Grik3* KD impaired the transfer of contagious pain (baseline *vs*. post: WT: 0.64 ± 0.07 g *vs*. 0.37 ± 0.07 g; KD: 0.76 ± 0.09 g *vs*. 0.74 ± 0.09 g; Fig. 6g-l), reduced the counts and duration of allo-licking/allo-grooming behaviors (WT *vs*. KD, duration: 171.00 ± 35.18 s *vs*. 77.10 ± 18.31 s; counts: 36.90 ± 6.59 *vs*. 17.20 ± 2.97; Fig. 6m, n and Extended Data Fig. 19f, g), and increased sniffing behaviors (Fig. 6o, p and Extended Data Fig. 19h, i), without affecting locomotion or anxiety-like responses (Extended Data Fig. 19j-p).

Conversely, overexpression of *Grik3* in the OFC of SI mice reversed these deficits. *Grik3*-overexpressing (OE) orbitofrontal neurons exhibited enhanced excitability (firing rate: WT *vs*. OE: 21.79 ± 1.92 Hz *vs*. 30.07 ± 1.33 Hz at 225 pA; Fig. 7a-c), and OE mice demonstrated restored contagious pain (baseline vs. post: WT: 0.66 ± 0.05 g *vs*. 0.64 ± 0.09 g; OE: 0.63 ± 0.07 g *vs*. 0.22 ± 0.04 g; Fig. 7d-i) and prosocial interactions (WT *vs*. OE, duration: 74.69 ± 12.39 s *vs*. 81.31 ± 13.45 s; counts: 14.77 ± 1.76 *vs*. 25.23 ± 2.24; Fig. 7j, k). Furthermore, *Grik3* OE significantly reduced both the counts and duration of sniffing behaviors compared to controls (Fig. 7l, m).

**Fig. 7.**
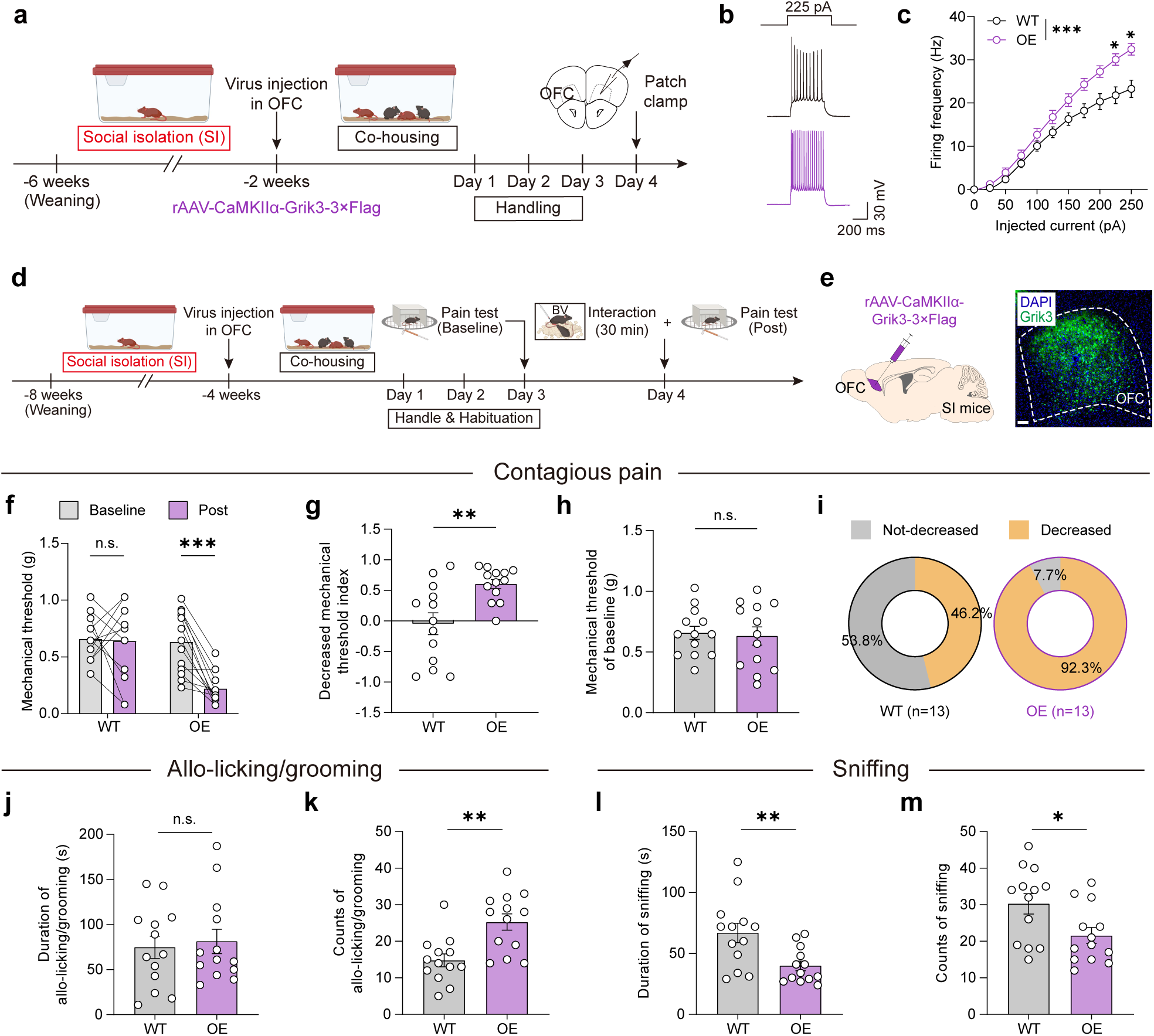
Overexpression of *Grik3* in OFC glutamatergic neurons rescues neuronal hypoexcitability and deficits in contagious pain and prosocial behaviors in SI mice. **a**, Experimental timeline depicting whole-cell patch-clamp recordings of OFC glutamatergic neurons following *Grik3* overexpression (OE) in SI mice. **b, c,** Representative traces (**b**) and quantification (**c**) of action potentials evoked by step current injections in OFC glutamatergic neurons from WT (black) and OE (purple) mice. (**c**) WT: *n* = 28 neurons from 4 mice; OE: *n* = 27 neurons from 5 mice. WT vs. OE: *p* < 0.0001; 0 pA: *p* > 0.9999; 25 pA: *p* = 0.7391; 50 pA: *p* = 0.7391; 75 pA: *p* = 0.7391; 100 pA: *p* = 0.7391; 125 pA: *p* = 0.3982; 150 pA: *p* = 0.3649; 175 pA: *p* = 0.1422; 200 pA: *p* = 0.1105; 225 pA: *p* = 0.0329; 250 pA: *p* = 0.0150. Two-way RM ANOVA with Mann-Whitney *post hoc* tests. **d,** Experimental timeline for *Grik3* overexpression in OFC glutamatergic neurons prior to social interaction in SI male mice. **e,** Schematic (left) and representative image (right) showing viral injection sites within the OFC. Scale bar, 100 μm. **f,** Changes in mechanical pain thresholds. *n* = 13 mice per group. WT: *p* > 0.9999; OE: *p* = 0.0003; two-way RM ANOVA with Bonferroni *post hoc* tests. **g,** Percentage decrease in mechanical pain thresholds. *n* = 13 mice per group. *p* = 0.0039, unpaired *t*-test with Welch’s correction. **h,** Baseline mechanical pain thresholds. *n* = 13 mice per group. *p* = 0.7802, unpaired *t*-test. **i,** Percentage of mice exhibiting or not exhibiting contagious pain. **j,** Total duration of prosocial behaviors. *n* = 13 mice per group. *p* = 0.7206, unpaired *t*-test. **k,** Number of prosocial behavior bouts. *n* = 13 mice per group. *p* = 0.0012, unpaired *t*-test. **l,** Total duration of sniffing behavior. *n* = 13 mice per group. *p* = 0.0069, unpaired *t*-test with Welch’s correction. **m,** Number of sniffing bouts. *n* = 13 mice per group. *p* = 0.0215, unpaired *t*-test. \**p* < 0.05, \*\**p* < 0.01, \*\*\**p* < 0.001, n.s., no significant difference. Data are shown as means ± SEM. See Supplementary Table 1 for full statistical details.

Together, these findings identify *Grik3* downregulation as a key molecular mechanism underlying SI-induced hypoexcitability in OFC glutamatergic neurons and associated deficits in empathic behaviors, both of which are reversible through targeted restoration of *Grik3* expression.

## Discussion

Empathy—the ability to perceive and share the emotional states of others—is fundamental to social life, promoting prosocial behaviors and reinforcing group cohesion^3,18,58^. While both intrinsic and environmental influences shape the empathic capacity, the underlying neurophysiological mechanisms remain incompletely understood. In this study, we identify the VM→ OFC excitatory projection as a critical neural substrate for empathy-like behaviors in mice. This pathway is essential for both contagious pain and prosocial behaviors, such as allo-licking and allo-grooming. Post-weaning SI disrupts these behaviors by inducing OFC hypoexcitability via downregulation of *Grik3* and attenuation of VM→OFC synaptic input. Crucially, restoring OFC excitability—through *Grik3* overexpression or chemogenetic activation of the OFC or its VM afferents—rescues these deficits (Fig. 8). These findings establish the VM→OFC circuit and its molecular components as key substrates for empathy-like behaviors, with translational relevance for understanding and treating social-affective impairments in neurodevelopmental and psychiatric conditions.

**Fig. 8.**
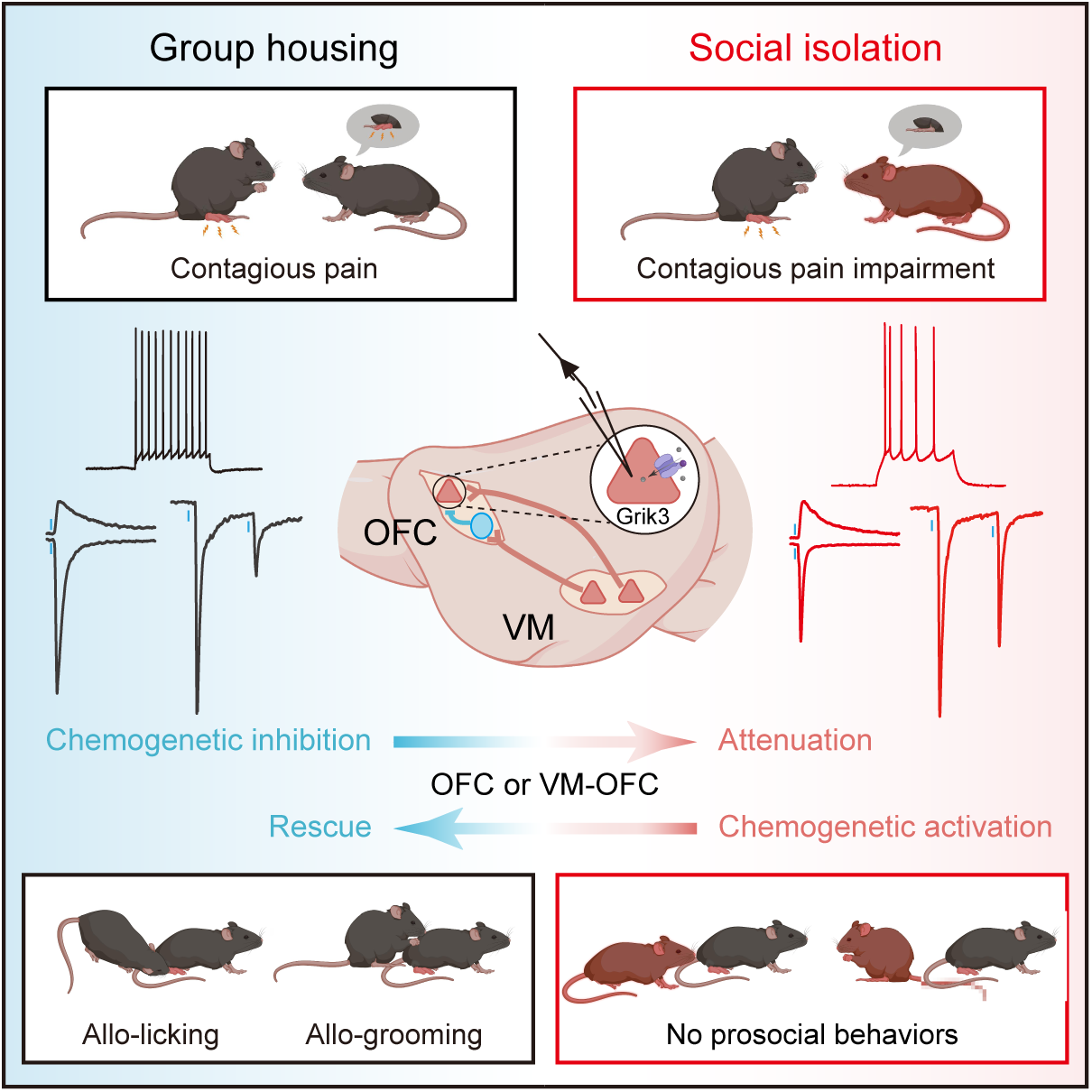
Schematic model of the molecular and circuit mechanisms underlying SI-induced empathic behavioral deficits. A balanced excitability of OFC glutamatergic neurons and intact synaptic transmission from the VM to OFC are essential for the proper expression of contagious pain and allo-grooming/allo-licking behaviors during observer-demonstrator interactions. Post-weaning SI markedly reduces OFC neuronal activity and weakens VM→OFC synaptic connectivity, leading to impairments in both pain contagion and prosocial responses. Chemogenetic activation of OFC glutamatergic neurons or the VM→OFC circuit restores these behaviors in SI-treated observers. *Grik3* emerges as a key molecule calibrating the orbitofrontal neuronal excitability and the empathic behavioral output.

Rodent models have significantly advanced our understanding of the neural basis of empathy-like behaviors, including contagious pain^11^, observational fear^22,23^, consolation^13–15^, and helping behavior^17,21,59^. Although several regions, such as the anterior cingulate cortex, insular cortex, and amygdala, have been implicated^60,61^, our results highlight the OFC as a previously underappreciated yet essential node in empathy-related processing. Using *in vivo* fiber photometry, we observed that OFC glutamatergic neurons are activated approximately one second before the initiation of social investigation or prosocial actions, suggesting a predictive role in detecting conspecific distress and initiating behavioral responses. Chemogenetic silencing of these neurons abolished both contagious pain and prosocial behaviors in a sex-independent manner. Previous work proposed that allo-grooming reflects consolation^13,62^, while allo-licking may represent targeted helping^17^. Our study suggests that OFC activity is required for both types of prosocial actions. Given the functional heterogeneity of the OFC in value encoding, multisensory integration, and affective regulation^63–65^, it is likely that distinct OFC subpopulations contribute to different facets of empathy. Future studies using projection- and cell-type-specific approaches will be instrumental in dissecting whether contagious pain and prosocial behaviors are mediated by discrete OFC outputs.

Beyond identifying the OFC as a functional hub for empathic processing in contagious pain and prosocial behaviors, we characterize upstream afferents that convey social-affective information. Prior studies have shown that negative valence cues related to others’ distress— delivered through visual or olfactory modalities—are integrated in higher-order cortical regions^66–68^. Using activity-dependent labeling and monosynaptic retrograde tracing, we identified significant input to OFC neurons engaged during social interaction from both the VM and the submedius thalamic nucleus (Sub). Projection-specific chemogenetic inhibition revealed that the VM→OFC pathway, but not the Sub→OFC projection, is necessary for empathic behaviors. Although the VM→OFC projection has previously been associated with analgesia following peripheral nerve injury^31^, our findings assign it a distinct role in socially transferred pain, underscoring the context-dependent function of this circuit in tuning pain sensitivity. Despite its anatomical connectivity and established role in nociceptive modulation^53,69^, the Sub→OFC pathway was dispensable in our empathy paradigm, highlighting the specificity of the VM→OFC circuit.

While adolescent social deprivation is known to increase the risk of anxiety, cognitive deficits, and impaired social behaviors^41–43,70,71^, its effects on higher-order social functions like empathy are less well explored. Although impairments in observational and vicarious fear learning after SI have been reported^72,73^, the impacts on contagious pain and prosocial behaviors remained unclear. Here, we show that four weeks of post-weaning SI, but not acute SI in adulthood, robustly impairs both pain contagion and prosocial behaviors, independent of sex. These behavioral deficits were not accompanied by alterations in anxiety-like behavior, locomotion, sociability, or social memory, indicating a selective disruption of empathy-related circuits. Moreover, the brief (2-week) re-grouping with naïve cage mates before testing may have minimized generalized stress, helping isolate the specific empathic phenotype. Mechanistically, SI reduced OFC neuronal excitability, dampened excitatory synaptic drive, and weakened VM→OFC connectivity—all of which were reversible through chemogenetic reactivation of this circuit. Thus, blunted orbitofrontal excitability and diminished synaptic strength represent key mechanisms through which SI disrupts empathy-like behaviors.

To uncover the molecular drivers of SI-induced OFC hypoexcitability, we performed single-nucleus RNA sequencing of OFC tissue. We identified significant downregulation of *Grik3*, which encodes the kainate receptor subunit GRIK3. GRIK3 has previously been implicated in psychiatric disorders such as schizophrenia and depression^56,74^. In our model, *Grik3* knockdown mimicked the physiological and behavioral deficits caused by SI, whereas overexpression of *Grik3* restored OFC excitability and rescued empathic behaviors. Although kainate receptors are known to modulate both presynaptic release and postsynaptic responses, the specific intracellular mechanisms by which GRIK3 regulates OFC excitability and social-affective behaviors remain to be elucidated.

In summary, we identify a previously unrecognized VM→OFC circuit and a *Grik3*-dependent molecular mechanism as central to empathy-like behaviors in mice. Post-weaning SI disrupts these processes by reducing OFC excitability and VM→OFC synaptic strength, resulting in selective impairments in contagious pain and prosocial behaviors. These findings establish a mechanistic framework for understanding how early-life social adversity alters social-affective brain circuits and may inform future therapeutic strategies targeting empathy-related deficits featured by several neuropsychiatric disorders.

## Methods

### Animals

Wild-type C57BL/6J mice (both sexes, aged 3 to 8 weeks) were obtained from the Shanghai Laboratory Animal Center (SLAC), Chinese Academy of Sciences. Transgenic mouse lines included Fos-CreER^T2^ (TRAP2; JAX Stock No. 030323) and CaMKIIα-Cre (JAX Stock No. 005359), purchased from The Jackson Laboratory. Mice were housed in groups of 4–5 per cage, or singly for SI experiments, under standard conditions (12-hour light/dark cycle, 21–25 °C) with ad libitum access to food and water. All behavioral testing was conducted during the light phase. Experimental procedures complied with the ethical guidelines approved by the Animal Ethics Committee of Shanghai Jiao Tong University School of Medicine (Policy Number: DLAS-MP-ANIM. 01-05).

## METHOD DETAILS

### Social isolation

The post-weaning social isolation (PWSI) paradigm was adapted from established protocols^42,45,75^. Three-week-old mice were randomly assigned to either group housing (GH) or social isolation (SI). GH mice were housed in same-sex groups, while SI mice were singly housed with a plastic shelter toy for four weeks. Both groups were then co-housed with same-sex, unfamiliar wild-type mice for a minimum of two weeks prior to behavioral testing to establish social familiarity. For acute social isolation (ASI), 7-week-old mice were initially co-housed with same-sex strangers for two weeks, after which ASI mice were isolated from their home cages and individually housed for three days before testing^49,76^.

### Virus constructs

The following AAVs were used: rAAV-CaMKIIα-GCaMP6m, rAAV-CaMKIIα-hM4D(Gi)-mCherry, rAAV-CaMKIIα-hM3D(Gq)-mCherry, rAAV-CaMKIIα-DIO-hM4D(Gi)-EGFP, rAAV2-retro-hSyn-Cre-mCherry, rAAV-EF1α-DIO-RVG, rAAV-EF1α-DIO-GFP-TVA, RV-EnvA-ΔG-DsRed, rAAV2/1-EF1α-Flp, rAAV-nEf1ɑ-fDIO-mCherry, rAAV-EF1α-DIO-H2B-EGFP, rAAV-CaMKIIα-ChR2-mCherry, rAAV-CaMKIIα-DIO-hM3D(Gq)-EGFP, rAAV-CaMKIIα-mCherry-5’miR-30a-shRNA3(*Grik3*)-3’-miR30a, and rAAV-CaMKIIα-*Grik3*-3×Flag. All vectors were serotype 2/9 unless otherwise specified and obtained from Brain VTA Co., Ltd (Wuhan, China), stored at –80 °C in aliquots (∼5.0 × 10¹² viral particles/mL). *Grik3* shRNA sequence: 5′-GGCTCAATGGGAAGGATTAAC-3′; Scrambled control: 5′-CCTAAGGTTAAGTCGCCCTCGA-3′.

### Stereotaxic surgeries

Stereotaxic injections were performed in 7–8-week-old mice under 1–1.5% isoflurane anesthesia, as previously described^52,77^. Animals were fixed in a stereotactic frame (RWD Life Science, Shenzhen, China), with ocular protection via erythromycin ointment and body temperature maintained at 37 °C. After disinfection and midline incision, bilateral craniotomies were made using a 0.5 mm microdrill. Glass pipettes (10–20 μm tip diameter, pulled using a P-1000 Micropipette Puller, Sutter Instrument) were used for injections. Pipettes were filled with AAV solution using silicone oil and connected to a microinjector (RWD Life Science) for precise delivery. Injection coordinates (from bregma, in mm) were as follows: OFC (AP, +2.58; ML, ±1.10; DV, –2.30), VM (AP, –1.40; ML, ±0.80; DV, –4.30), Sub (AP, –1.40; ML, ±0.40; DV, –4.10). Volumes injected per site were OFC (0.2 μL), VM (0.25 μL), and Sub (0.15 μL) bilaterally at a rate of 0.1 μL/min. Pipettes remained in place for 10 min post-injection to facilitate diffusion. For fiber photometry, a 200 μm diameter optical cannula (NA 0.37, Inper, Hangzhou, China) was implanted 0.15 mm above the OFC injection site and secured with dental cement and skull screws. Mice recovered for at least two weeks before testing. Injection accuracy was confirmed *post hoc* via expression of fluorescent markers (*e.g*., EGFP, mCherry), and animals with mistargeted or widespread expression were excluded.

### Behavior assays

#### Empathic social interaction test

Empathic social interaction was assessed using a previously validated paradigm^46^. Testing occurred in medium-sized, non-transparent acrylic boxes (19 × 19 × 30 cm) with 100 g of bedding. Observer-demonstrator pairs (matched for sex and body weight difference <10%) were co-housed for at least two weeks prior to testing to establish social familiarity. Mice were then acclimated to handling and the testing environment over three consecutive days, during which baseline mechanical pain thresholds of observers were also recorded.

On the test day, pairs were habituated in the interaction box for 10 minutes. Then, demonstrators received an intraplantar injection of 0.4% bee venom (lyophilized *Apis mellifera* venom in physiological saline; 25 μL) into the left hind paw to induce robust inflammatory pain.

Once visible paw flinching/lifting/licking confirmed the nociceptive activation, demonstrators were gently returned to the box for a 30-minute free interaction session, recorded from above using a Logitech C920e camera (Switzerland). Immediately after the social interaction period, observers were tested again to examine possible changes in mechanical pain sensitivity.

For empathic social interaction assessment, the sequential behaviors displayed by the observers during interaction with the demonstrators were manually scored from the videos. Prosocial behaviors included allo-licking (licking the injured paw) and allo-grooming (grooming the face, ears, or back of the demonstrator)^14,15^. Sniffing was defined as investigation of the demonstrator’s tail, anus, or body. Self-grooming involved face washing or body licking, consistent with previous reports^78,79^. Pairs exhibiting aggressive behavior were excluded from analysis.

For chemogenetic manipulations, mice received intraperitoneal injections of either saline or clozapine-N-oxide (CNO; 2 mg/kg; ENZO, BML-NS105-0025), prepared at 0.25 mg/mL in 0.9% sterile saline and administered at least 30 minutes before testing.

#### Mechanical pain threshold measurement

Mechanical nociceptive thresholds were assessed using von Frey filaments (North Coast Medical, Inc., USA) following the up-down paradigm^80^. Mice were individually placed on a metal mesh platform with 0.5 × 0.5 cm openings and covered by opaque acrylic enclosures (10.5 × 10.5 × 5 cm). Calibrated filaments were applied perpendicularly to the central plantar surface of the hind paw for a maximum of 5 seconds. A positive response—defined as paw withdrawal, flinching, or licking—was recorded as indicative of nociceptive response. Inter-stimulus intervals were set at a minimum of 20 seconds to prevent sensitization.

#### Open field test

The open field test (OFT) was performed in a Plexiglas arena (40 × 40 × 35 cm) subdivided into a central zone (20 × 20 cm) and a surrounding periphery. Mice were gently placed in the center and allowed to explore freely for 15 minutes. Locomotor activity, including time spent in the central zone, distance traveled within it, total distance moved and velocity, was recorded and analyzed using Noldus EthoVision XT (v17.0, Noldus Information Technology, Netherlands). Chambers were thoroughly cleaned with 75% ethanol between trials to eliminate olfactory cues.

#### Elevated plus maze test

The elevated plus maze (EPM) consisted of two open arms (30 × 5 cm) and two enclosed arms (30 × 5 × 15 cm) arranged orthogonally and connected by a central platform (5 × 5 cm). The apparatus was elevated 50 cm above the floor. Mice were placed at the central junction and allowed to explore for 8 minutes. Time spent in the open arms and the number of entries into each arm type were recorded using Noldus EthoVision XT (v17.0).

#### Three-chamber test

Sociability and social novelty were evaluated using a three-chambered apparatus (65 × 20 × 25 cm) with two side chambers (25 × 20 × 25 cm) and a central chamber (15 × 20 × 25 cm)^81^. Cylindrical wire cages (10 × 10 × 10 cm) were positioned in the corners of the side chambers. During the first 10-minute habituation session, mice explored the apparatus freely. In the second session, a novel, same-sex conspecific (stranger 1) was placed under one wire cage while the opposite chamber remained empty. In the third session, another novel conspecific (stranger 2) was introduced into the previously unoccupied cage. The assignment of strangers to chambers was counterbalanced. Exploration time around each cage was quantified using Noldus EthoVision XT (v17.0). Sociability and social novelty preference indexes were computed as follows: Sociability index = (Time with stranger 1 − Time with empty cage) / (Time with stranger 1 + Time with empty cage); Social novelty index = (Time with stranger 2 − Time with stranger 1) / (Time with stranger 1 + Time with stranger 2).

### Fiber photometry

Fiber photometry was performed using a dual-color system (ThinkerTech, Nanjing, China), following established protocols with minor modifications^77^. Implanted optical fibers were coupled to the recording system through patch cords. A 488-nm laser (OBIS 488LS, Coherent, USA) was reflected by a dichroic mirror (MD498, Thorlabs, USA), focused through a 10× objective lens (NA = 0.3, Olympus, Japan), and directed into an optical commutator (FRJ_1 × 1_FC-FC, Doric Lenses, Canada). A 580-nm laser served as an internal control. Light was transmitted via a 200-μm core, 0.37 NA optical fiber (1 m, Inper, Hangzhou, China), with power at the tip adjusted to 40–60 μW to minimize GCaMP6m photobleaching. Emitted signals were filtered through a bandpass filter (MF525-39, Thorlabs, USA) and detected by a photomultiplier tube (R3896, Hamamatsu, Japan), with output current converted to voltage (C7319, Hamamatsu, Japan) and filtered through a 40-Hz low-pass filter (Brownlee 440). Analog signals were digitized at 100 Hz using a Power 1401 interface and Spike2 software (CED, UK). For empathic interaction recordings, observer-demonstrator dyads were habituated to the test box over a period of three days. During this period, observers were connected to the photometry system via optical fibers. During the 30-minute test, GCaMP6m fluorescence was recorded concurrently with behavioral video footage. Behavioral scoring—including sniffing, allo-licking, and allo-grooming—was performed manually in a double-blind manner.

Calcium signal data were exported to MATLAB for analysis. Z-scores were computed for each fluorescence value (F) using the equation: Z = (F − F_baseline) / SD_baseline, where F_baseline and SD_baseline represent the mean and standard deviation of fluorescence during the 1-second pre-interaction baseline (from −2 to −1 s before behavior onset). Z-score traces were aligned to behavioral events and averaged across trials within animals and across animals within groups. Resulting Z-score traces were visualized as peri-event plots and heatmaps, with shaded error bars representing the SEM. Quantitative measures included the peak Z-score within 3 seconds post-behavior onset and the area under the curve (AUC) over the same interval.

### *In vitro* electrophysiological recordings

Whole-cell patch-clamp recordings were performed on acute brain slices following established protocols^69,82^. Mice were deeply anesthetized with isoflurane and rapidly perfused with ∼15 mL of ice-cold N-methyl-D-glucamine-based artificial cerebrospinal fluid (NMDG-aCSF), containing: 93 mM NMDG, 93 mM HCl, 2.5 mM KCl, 1.25 mM NaH₂PO₄, 10 mM MgSO₄, 30 mM NaHCO₃, 0.5 mM CaCl₂, 25 mM glucose, 20 mM HEPES, 5 mM sodium ascorbate, 3 mM sodium pyruvate, and 2 mM thiourea (pH 7.3–7.5; osmolarity 295–305 mOsm). Coronal slices (300 μm thick) encompassing the OFC were prepared using a vibratome (VT-1000S, Leica, Germany) in ice-cold NMDG-aCSF. Slices were initially incubated at 31 °C in the same solution for 5–10 minutes and subsequently transferred to room temperature standard aCSF for recovery (≥ 45 minutes). The aCSF consisted of 125 mM NaCl, 2.5 mM KCl, 12.5 mM glucose, 1 mM MgCl₂, 2 mM CaCl₂, 1.25 mM NaH₂PO₄, and 25 mM NaHCO₃ (pH 7.3–7.5; osmolarity 280–300 mOsm), continuously bubbled with 95% O₂/5% CO₂ to maintain physiological pH and oxygenation.

Slices were transferred to a perfusion chamber maintained at 32 °C and superfused with aCSF at 2–3 mL/min. Neurons in the OFC were visualized using an upright fluorescence microscope (BX51WI, Olympus, Japan) equipped with differential interference contrast optics and a 40× water-immersion objective. Patch pipettes (3–5 MΩ) were pulled from borosilicate glass (Sutter Instrument, USA) using a P-2000 micropipette puller (Sutter Instrument). Recordings were obtained using an Axon 200B amplifier (Molecular Devices, USA), low-pass filtered at 2 kHz, and digitized at 10 kHz via a Digidata 1440A interface. Series resistance and input resistance were monitored throughout the experiments. Recordings were discarded if a substantial change in series resistance (>20%) was detected.

To assess the intrinsic excitability of OFC pyramidal neurons following SI, mice were single-housed for 4 weeks and then re-socialized for 2 weeks prior to slice preparation. Current-clamp recordings were performed using an internal solution composed of 145 mM potassium gluconate, 5 mM NaCl, 10 mM HEPES, 2 mM Mg-ATP, 0.1 mM Na₂-GTP, 0.2 mM EGTA, and 1 mM MgCl₂ (pH 7.3, adjusted with KOH; osmolarity 280–290 mOsm). Neuronal firing was evoked by 500-ms current steps from 0 to 250 pA (25 pA increments). Parameters such as action potential threshold, amplitude, and rheobase were analyzed as described previously^52^. To determine the role of *Grik3* in OFC excitability, we delivered rAAV-CaMKIIα-mCherry-5′miR-30a-shRNA3(*Grik3*)-3′-miR30a to knock down *Grik3* in GH mice, and rAAV-CaMKIIα-*Grik3*-3×Flag to overexpress *Grik3* in SI mice. Brain slices were prepared 3–4 weeks post-injection, and OFC neuronal activity was recorded as above.

To evaluate synaptic input changes, spontaneous excitatory postsynaptic currents (sEPSCs) and spontaneous inhibitory postsynaptic currents (sIPSCs) were recorded in voltage-clamp mode at –70 mV and 0 mV, respectively, using a Cs-based internal solution containing 132.5 mM Cs-gluconate, 17.5 mM CsCl, 2 mM MgCl₂, 0.5 mM EGTA, 10 Mm HEPES, and 4 mM Mg-ATP (pH 7.3, adjusted with CsOH; osmolarity 280–290 mOsm). The baseline sEPSCs or sIPSCs were recorded for 5 min and were analyzed from 100 to 200 s after the establishment of the recording.

To interrogate synaptic transmission and plasticity in the VM→OFC projection, rAAV-CaMKIIα-ChR2-mCherry was injected into the VM, and OFC recordings were performed under blue-light stimulation three weeks after virus injection. Photostimulation (473 nm, 15 s, 0.66 Hz, 5-ms pulse) was delivered through a 40× objective using a collimated LED (Lumen Dynamics Group Inc, USA) interfaced with the Axon 200B amplifier. The intensity of photostimulation was directly controlled by the stimulator, while the duration was set by the pClamp 10.5 software. Light-evoked EPSCs/IPSCs were recorded and pharmacologically validated using CNQX (10 μM) and APV (50 μM) for glutamatergic currents, and PTX (100 μM) for GABAergic currents. Local inhibition within the OFC was dissected by applying PTX or CNQX plus APV.

Monosynaptic connectivity was verified using sequential bath application of TTX (0.5 μM) and 4-aminopyridine (4-AP, 100 μM). Paired-pulse ratio (PPR) was measured at –70 mV or 0 mV by delivering paired photostimulation pulses (5 ms duration) at inter-stimulus intervals of 25, 50, 100, 200, and 500 ms. PPR was calculated as the amplitude ratio of the second to the first evoked current. AMPAR/NMDAR ratios were determined from EPSC peak amplitudes recorded at –70 mV (AMPAR, in the presence of 100 μM PTX) and +40 mV (NMDAR, in the presence of 100 μM PTX and 20 μM CNQX). Data were analyzed offline using Clampfit v10.6 (Molecular Devices) and Mini Analysis v6.03 (Synaptosoft).

### Activity-dependent neuronal labeling

Activity-dependent labeling of neurons was performed using TRAP2 mice as previously described^11,52,83^. To induce recombination, 4-hydroxytamoxifen (4-OHT; Sigma-Aldrich, H6278, USA) was freshly prepared at 10 mg/mL. Briefly, a 20 mg/mL stock was dissolved in ethanol at 37 °C for 30 minutes, followed by the addition of twice the volume of corn oil (Sigma-Aldrich, C8267, USA). Ethanol was then removed by vacuum centrifugation. The resulting solution was stored at 4 °C and used within 24 hours. To reduce immediate early gene activation due to handling, mice were relocated from the vivarium to a nearby room at least 3 hours before 4-OHT injection. All injections were administered intraperitoneally. Prior to FosTRAP labeling, observer-demonstrator pairs were handled and habituated for 3 days. Observer mice were then administered 4-OHT (50 mg/kg) at least 30 minutes before engaging in a 120-minute social interaction with a demonstrator in pain. Afterward, mice were returned to the vivarium.

### Subpopulation-specific input tracing

To map monosynaptic inputs to socially activated neurons, we employed rabies virus retrograde tracing in TRAP2 mice. On Day 1, a Cre-dependent AAV cocktail (1:1 ratio of rAAV-EF1α-DIO-RVG and rAAV-EF1α-DIO-EGFP-TVA) was injected into the OFC. FosTRAP labeling was performed 16 days (including the 14 days co-housing time) after the virus injection. After three additional weeks, RV-EnvA-ΔG-DsRed was injected into the same site, and mice were perfused one week later. Brains were post-fixed overnight and coronally sectioned at 40 μm (VT-1000S, Leica, Germany) with a slicing speed of 0.6–0.8 mm/s. Starter cells (GFP^+^/DsRed^+^) were identified, and the distribution of DsRed^+^ monosynaptic inputs was examined among all brain sections.

Imaging was conducted on an Olympus VS200 microscope using 10× objectives with consistent settings across experimental groups. Subsequent analyses were performed using the Aligning Big Brains & Atlases (ABBA) plugin in ImageJ (Fiji edition, National Institutes of Health, USA) and QuPath software^84^.

### Anterograde tracing of VM→OFC projections

To identify the specific OFC cell types receiving inputs from the VM, a combinatorial viral tracing strategy was employed. rAAV2/1-EF1α-Flp was injected into the VM, while a mixture of rAAV-nEF1α-fDIO-mCherry and rAAV2/9-EF1α-DIO-H2B-EGFP was delivered into the OFC of CaMKIIα-Cre mice. This strategy selectively labeled VM-innervated OFC neurons with mCherry, whereas all OFC glutamatergic neurons expressed EGFP. After 3–4 weeks of viral expression, mice were transcardially perfused, and 40 μm coronal brain sections were collected. Fluorescence images were captured using an Olympus VS200 slide scanner equipped with a 10× objective, applying uniform acquisition parameters across all samples. Cell counting and colocalization analysis were conducted using the ABBA plugin in Fiji and QuPath software. The proportion of mCherry^+^/EGFP^+^ double-labeled cells was quantified relative to the total mCherry^+^ population.

### Single-nucleus RNA sequencing

#### Sample collection

Following weaning, mice underwent a four-week period of social isolation, followed by two weeks of co-housing with age- and sex-matched wild-type strangers. At the conclusion of the paradigm, mice were deeply anesthetized with isoflurane and rapidly decapitated. Brains were immediately harvested and placed in a chilled brain matrix (RWD, China) on ice. Coronal sections (1-mm thick) encompassing the OFC were obtained using precooled blades, and the OFC was microdissected using sterile, chilled forceps. Dissected tissues were transferred into precooled microcentrifuge tubes and flash-frozen on dry ice for subsequent nuclei isolation.

#### Nuclei suspension preparation

All reagents were prechilled prior to use. The lysis buffer contained 0.25 M sucrose, 5 mM CaCl₂, 3 mM MgAc₂, 10 mM Tris-HCl (pH 8.0), 1 mM DTT, 0.1 mM EDTA, 1× protease inhibitor cocktail, and 1 U/µL RiboLock RNase Inhibitor. Iodixanol solutions (30%, 33%, 50%) were composed of 0.16 M sucrose, 10 mM NaCl, 3 mM MgCl₂, 10 mM Tris-HCl (pH 7.4), 1 mM DTT, 0.1 mM PMSF, and 1 U/µL RiboLock. The nuclei wash buffer consisted of PBS with 0.04% BSA, 0.2 U/µL RiboLock, 500 mM mannitol, and 0.1 mM PMSF.

Frozen brain tissue samples (≥ 0.2 g) were homogenized in 500 µL lysis buffer using a Dounce homogenizer (15–20 strokes with a loose pestle, followed by 5–10 strokes with a tight pestle). The homogenate was combined with 700 µL wash buffer and filtered through a 70 µm cell strainer. Equal volumes of the filtrate and 50% iodixanol were mixed and layered onto a step gradient (1 mL 33% and 2 mL 30% iodixanol) in a 15 mL tube. After centrifugation at 1,000 *×* g for 10 minutes at 4 °C, nuclei at the 30–33% interface were collected, diluted in wash buffer, and pelleted at 500 × g for 5 minutes at 4 °C. The resulting pellet was gently resuspended, passed through a 40 µm cell strainer, and stained with Trypan Blue to evaluate nuclei integrity. The final suspension was adjusted to 700–1200 nuclei/μL for downstream applications.

#### Single-nucleus library construction and sequencing

Nuclei were processed using the 10× Genomics Chromium Single-Nucleus 3’ platform to generate Gel Bead-In-Emulsion (GEM) droplets. Libraries were constructed using the Chromium Next GEM Single Cell 3’ Reagent Kits v3.1. Upon GEM dissolution, primers containing the Illumina Read 1 sequence, a 16-nt 10× cell barcode, a 10-nt unique molecular identifier (UMI), and a poly-dT sequence were released and mixed with the nuclei lysate. Barcoded, full-length cDNAs were synthesized from polyadenylated mRNA via reverse transcription and then amplified by PCR. Library construction steps involved end repair, A-tailing, adapter ligation, and indexing PCR, incorporating the Illumina P5 and P7 sequences. The final libraries, compatible with Illumina paired-end sequencing, incorporated the cell barcodes and UMIs in Read 1, and captured cDNA sequences in Read 2. The sample index was incorporated as the i7 index read. Read 1 and Read 2 were standard Illumina sequencing primer sites used in paired-end sequencing.

#### Bioinformatic analysis

Transcriptomic data analysis was conducted on the Omicsmart platform (Gene Denovo Biotechnology Co., Ltd, China; http://www.omicsmart.com). Raw BCL files were processed using 10× Genomics Cell Ranger v6.1.0 to generate FASTQ files, align reads, and perform UMI quantification. Reads mapping uniquely to the transcriptome and overlapping an exon by at least 50% were retained. UMIs were corrected for sequencing errors, and valid barcodes were identified via the EmptyDrops method^85^. Cell-by-gene expression matrices were generated from these data via UMI counting and cell barcodes calling.

*Principal component analysis (PCA)*. The integrated gene expression matrix was scaled and subjected to PCA to reduce dimensionality. To identify significant principal components (PCs), we performed a resampling test based on the JackStraw approach, selecting PCs enriched for genes with low *p*-values for downstream clustering and further dimensionality reduction^86^.

*Cell clustering, visualization, and annotation.* Cell-by-gene expression matrices were analyzed using Seurat’s graph-based clustering framework. Pairwise distances between cells were computed using the previously identified significant PCs. Clustering was then performed using the Louvain algorithm to optimize modularity^87^. For visualization, we applied t-distributed stochastic neighbor embedding (t-SNE) to the same PCs. Cell clusters were manually annotated based on the expression of canonical cell-type-specific markers.

*Differentially expressed genes (DEGs) analysis.* To further analyze the DEGs between GH and SI groups, a hurdle model was applied using the Model-based Analysis of Single-cell Transcriptomics (MAST) framework, adapted for single-nucleus data. DEGs between groups were identified by the following criteria: |log_2_FC| ≥ 0.36; *p* < 0.05; percentage of nuclei where the gene was detected in a specific cluster ≥ 10%. Identified DEGs were subsequently subjected to pathway enrichment analysis.

*Pathway enrichment analysis.* To identify biologically relevant pathways enriched among DEGs, we performed enrichment analysis using the Kyoto Encyclopedia of Genes and Genomes (KEGG) database^88^. Pathway enrichment was evaluated against the genome-wide background using statistical methods described previously^89^, and *p*-values were corrected for multiple testing using the false discovery rate (FDR) method. Pathways with FDR ≤ 0.05 were deemed significantly enriched.

### Immunofluorescence staining

Mice were deeply anesthetized with isoflurane and perfused with 20 mL of ice-cold saline followed by 10 mL of ice-cold 4% paraformaldehyde in 0.1 M phosphate-buffered saline (PBS). Brains were carefully extracted and post-fixed overnight in 4% paraformaldehyde at 4 °C, and then cryoprotected in 30% sucrose until they sank to the bottom. Coronal brain sections (40 μm) were prepared using a vibratome (VT-1000S, Leica, Germany). To assess viral infection efficiency, sections were incubated with 4′,6-diamidino-2-phenylindole dihydrochloride hydrate (DAPI, 1:2000; Thermo Fisher Scientific, #D1306) for 15 minutes, followed by three 15-minute washes in PBS containing 0.1% Tween-20 (PBST). Slices were mounted on glass slides and coverslipped using mounting media. Optical fiber placements were verified using standard histological procedures.

For immunofluorescence staining, sections were washed in PBS and blocked in 0.01 M PBS containing 10% donkey serum and 0.3% Triton X-100 for 1 hour at room temperature. Sections were then incubated overnight at 4 °C in primary antibody solutions prepared in 0.01 M PBS with 2% donkey serum. The primary antibodies used were anti-c-Fos (1:500; Cat. #2250, Cell Signaling Technology, USA) and anti-GRIK3 (1:500; Cat. #180203, SYSY, Germany). After three PBST washes, sections were incubated in a dark chamber for 2 hours at room temperature with fluorophore-conjugated secondary antibody (Alexa Fluor 488, donkey anti-rabbit, 1:200; Cat. #A21206, Thermo Fisher Scientific, USA) and DAPI (1:1000; Cat. #D9542, Sigma-Aldrich). Following final washes in PBST, sections were mounted using an antifade reagent. All fluorescence images, including the verification of virus expression and fiber/cannula implantation, were acquired using Olympus VS200 system with 10× objectives. Imaging parameters were kept constant across all groups. The numbers of c-Fos^+^ neurons were counted in four slices for each mouse through different bregma planes of the OFC region. All cell counting was carried out manually offline in a blind manner using the Aligning Big Brains & Atlases (ABBA) plugin in ImageJ (Fiji) and QuPath software.

## Statistics

All statistical analyses and graph generation were conducted using GraphPad Prism 10.3 (GraphPad Software Inc., USA). Unless otherwise specified, data were first assessed for normality and homogeneity of variance. Statistical tests included unpaired and paired Student’s *t*-tests (with or without Welch’s correction), Wilcoxon matched-pairs signed rank test, Mann-Whitney test, and one-way or two-way repeated-measures ANOVA with Bonferroni *post hoc* tests. No statistical methods were used to predetermine sample size. Results are reported as mean ± SEM unless indicated otherwise. A significance threshold of *p* < 0.05 was applied. Details on sample sizes, statistical tests, and exact *p*-values are provided in Supplementary Table S1.

## Data availability

All data supporting the findings of this study are available within the paper and its Extended Data. Source data underlying all main and Extended figures are provided with the paper.

## Code availability

No original code was generated in this paper.

## Supporting information

Main text

## Acknowledgements

We thank Jun B Ding (Stanford University), Michael X. Zhu (University of Texas Health Science Center at Houston), Lu-Yang Wang (University of Toronto), Qiufu Ma (Westlake University) and Bo Li (Westlake University) for providing helpful comments on the manuscript. This study was supported by grants from the STI2030–Major Projects (2021ZD0202800, 2021ZD0203302), the National Natural Science Foundation of China (32430040, 82571405), Shanghai Natural Science Foundation (25ZR1402302), and the innovative research teams of high-level local universities in Shanghai.

## Author contributions

Y.G., S.S.G., and X.C. performed experiments, analyzed the data and prepared the manuscript. Z.Z., X.L.S., Q.J., Y.L., and T.T.Z. contributed to data collection, interpretation and analysis. D.W. and X.Z. provided helpful comments on the manuscript. Y.G., H.J., M.G.L., and T.L.X. wrote and edited the manuscript. M.G.L. and T.L.X. designed experiments, supervised the experiments, and finalized the manuscript. All authors have read and approved the paper.

