## Supplementary material for "Post-weaning social isolation impairs the orbitofrontal cortical circuit subserving contagious pain and prosocial behaviors": Main text

#### Extended Data Figure Legends

**Extended Data Fig. 1 Additional behavioral outcomes in the empathic social interaction test for male and female mice.**

**a,** Baseline mechanical pain thresholds prior to empathic social interaction in male mice. $n = 12$ mice per group. $p = 0.2322$, unpaired *t*-test. **b,** Proportion of male mice exhibiting or not exhibiting contagious pain. **c, d,** Same analyses as (**a** and **b**), but for female mice. (**c**) $n = 12$ mice per group, $p = 0.0749$, Mann-Whitney test. **e,** Latency to initiate allo-licking or allo-grooming in males during the 30-minute empathic session. $n = 12$ mice per group; $p = 0.0006$, Mann-Whitney test. **f,** Duration per bout of allo-licking/grooming in males. $n = 12$ mice per group, $p = 0.338$, unpaired *t*-test with Welch's correction. **g, h,** Same as (**e** and **f**), but for females. $n = 12$ mice per group. (**g**) $p = 0.1734$, Mann-Whitney test; (**h**) $p = 0.0084$, unpaired *t*-test with Welch's correction. **i,** Latency to initiate sniffing in males during the 30-minute session. $n = 12$ mice per group, $p = 0.2108$, Mann-Whitney test. **j,** Duration per bout of sniffing in males. $n = 12$ mice per group, $p = 0.0222$, unpaired *t*-test. **k, l,** Same as (**i** and **j**), for females. (**k**) $p = 0.0411$, Mann-Whitney test; (**l**) $p = 0.9655$, Mann-Whitney test. **m, n,** Total duration (**m**) and number of bouts (**n**) of self-licking/grooming in males. $n = 12$ mice per group; (**m**) $p = 0.4577$, (**n**) $p = 0.365$, both unpaired *t*-tests. **o, p,** Same analyses as (**m** and **n**), but for females. (**o**) $p = 0.015$, (**p**) $p = 0.2247$, both unpaired *t*-tests with Welch's correction. **q,** Latency to initiate self-licking/grooming in males. $n = 12$ mice per group. $p = 0.7553$, Mann-Whitney test. **r,** Duration per bout of self-licking/grooming in males. $n = 12$ mice per group. $p = 0.7951$, unpaired *t*-test. **s, t,** Same as (**q** and **r**), but for females. (**s**) $p = 0.9656$, Mann-Whitney test; (**t**) $p = 0.0112$, unpaired *t*-test with Welch's correction. $^{*}p < 0.05$, $^{**}p < 0.01$, $^{***}p < 0.001$, n.s., no significant difference. Data are presented as means ± SEM. See Supplementary Table 1 for detailed statistics.

**Extended Data Fig. 2 Prosocial and sniffing behaviors predominantly occur within the first 5 minutes of the 30-minute empathic interaction.**

**a,** Total duration of allo-licking/grooming in each 5-minute interval during the 30-minute session in male mice. $n$ = 12 mice per group. Group comparison: GH vs. SI, $p = 0.2095$; by interval: 0–5 min: $p = 0.1129$, 5–10 min: $p = 0.5332$, 10–15 min: $p = 0.0084$, 15–20 min: $p = 0.2200$, 20–25 min: $p = 0.0001$, 25–30 min: $p = 0.5332$. Two-way RM ANOVA with Mann-Whitney *post hoc* tests. **b,** Number of allo-licking/grooming bouts per interval. GH vs. SI, $p = 0.0234$; 0–5 min: $p = 0.0149$, 5–10 min: $p = 0.4162$, 10–15 min: $p = 0.0053$, 15–20 min: $p = 0.1037$, 20–25 min: $p = 0.0003$, 25–30 min: $p = 0.4162$. Two-way RM ANOVA with Mann-Whitney *post hoc* tests. **c,** Cumulative percentage of allo-licking/grooming duration over time. GH vs. SI, $p = 0.1608$; at 5 min: $p = 0.4514$, 10 min: $p = 0.3805$, 15 min: $p = 0.4562$, 20 min: $p = 0.0094$, 25 min: $p = 0.6811$, 30 min: $p > 0.9999$. Two-way RM ANOVA with Mann-Whitney *post hoc* tests. **d,** Total sniffing duration per 5-minute interval. GH vs. SI, $p < 0.0001$; 0–5 min: $p = 0.0023$, 5–10 min: $p = 0.1956$, 10–15 min: $p = 0.1073$, 15–20 min: $p = 0.1956$, 20–25 min: $p = 0.3599$, 25–30 min: $p = 0.2028$. Two-way RM ANOVA with Mann-Whitney *post hoc* tests. **e,** Number of sniffing bouts per interval. GH vs. SI, $p < 0.0001$; 0–5 min: $p = 0.0017$, 5–10 min: $p = 0.2478$, 10–15 min: $p = 0.1926$, 15–20 min: $p = 0.0933$, 20–25 min: $p = 0.3870$, 25–30 min: $p = 0.2478$. Two-way RM ANOVA with Mann-Whitney *post hoc* tests. **f,** Cumulative percentage of sniffing duration across time intervals. GH vs. SI, p = 0.6853; 5–30 min: all $p > 0.97$. Two-way RM ANOVA with Mann-Whitney *post hoc* tests. **g-i,** Same analyses as (**d-f**), applied to self-licking/grooming behaviors. $n$ = 12 mice per group. All two-way RM ANOVA with Mann-Whitney *post hoc* tests. **g,** Total duration: GH vs. SI, $p = 0.4002$; 0–5 min: $p = 0.9438$, 5–10 min: $p = 0.5344$, 10–15 min: $p = 0.9438$, 15–20 min: $p = 0.9438$, 20–25 min: $p = 0.3031$, 25–30 min: $p = 0.9250$. **h,** Number of bouts: GH vs. SI, $p = 0.2227$; all individual intervals $p > 0.38$. **i,** Cumulative duration: GH vs. SI, $p = 0.0561$; 5–30 min: all $p > 0.21$. *$p < 0.05$, **$p < 0.01$, ***$p < 0.001$, n.s., no significant difference. Data are presented as means ± SEM. See Supplementary Table 1 for detailed statistics.

**Extended Data Fig. 3 SI does not alter anxiety-like behavior or locomotion.**

**a,** Experimental timeline for anxiety-related behavioral test following GH or SI in male and female mice. **b,** Representative locomotor traces in the open field test (OFT) for GH (dark grey) and SI (orange) male mice. **c-e,** Quantification of OFT in males. GH: $n = 16$; SI: $n = 10$. (**c**) Time spent in the center zone: $p = 0.9902$. (**d**) Distance traveled in the center: $p = 0.5457$. (**e**) Total distance traveled: $p = 0.7898$. All unpaired *t*-tests. **f,** Representative traces from the elevated plus maze (EPM) for GH and SI male mice. **g-i,** EPM results in males. GH, $n = 12$; SI, $n = 11$. (**g**) Time spent in open arms: $p = 0.2566$. (**h**) Open arm entries: $p = 0.1126$. (**i**) Total arm entries: $p = 0.2386$. All unpaired *t*-tests. **j-m,** OFT results in females. $n = 12$ mice per group. (**k**) Center time: $p = 0.2594$. (**l**) Center distance: $p = 0.9083$. (**m**) Total distance: $p = 0.92$. All unpaired *t*-tests. **n-q,** EPM results in females: GH, $n = 11$; SI, $n = 12$. (**o**) Time in open arms: $p = 0.4973$. (**p**) Open arm entries: $p = 0.5763$. (**q**) Total entries: $p = 0.8506$. All unpaired *t*-tests. n.s., no significant difference. Data are shown as means ± SEM. See Supplementary Table 1 for detailed statistics.

**Extended Data Fig. 4 SI does not affect sociability or social novelty behavior.**

**a,** Timeline for the three-chamber test following GH or SI in male and female mice. **b,** Heatmaps showing exploratory behavior in the sociability phase for GH (black) and SI (orange) male mice. E: empty cup; S1: stranger 1 mouse. **c,** Time spent investigating the empty vs. stranger 1 cup. $n =$

10 mice per group. GH: $p = 0.0002$; SI: $p = 0.0005$. Two-way RM ANOVA with Bonferroni *post hoc* tests. **d,** Sociability preference index: (stranger 1 − empty)/(stranger 1 + empty). $n = 10$ mice per group. $p = 0.7103$, unpaired *t*-test. **e,** Heatmaps during the social novelty phase. S1: stranger 1; S2: stranger 2. **f,** Time spent investigating the S1 vs. S2 cup. $n = 10$ mice per group. GH: $p = 0.0209$; SI: $p = 0.0044$. Two-way RM ANOVA with Bonferroni *post hoc* tests. **g,** Social novelty preference index: (stranger 2 − stranger 1)/(stranger 2 + stranger 1). $p = 0.7142$, unpaired *t*-test. **h-j,** Sociability data for females. $n = 12$ mice per group. (**i**) GH: $p = 0.0002$; SI: $p = 0.0012$. Two-way RM ANOVA with Bonferroni *post hoc* tests. (**j**) Preference index: $p = 0.5599$, unpaired *t*-test. **k-m,** Social novelty data for females. $n = 12$ mice per group. (**l**) GH: $p = 0.0070$; SI: $p = 0.1134$. Two-way RM ANOVA with Bonferroni *post hoc* tests. (**m**) Preference index: $p = 0.3819$, unpaired *t*-test. *$p < 0.05$, **$p < 0.01$, ***$p < 0.001$, n.s., no significant difference. Data are presented as means ± SEM. See Supplementary Table 1 for detailed statistics.

**Extended Data Fig. 5 ASI in adulthood does not influence contagious pain or prosocial behaviors.**

**a,** Schematic timeline of the 3-day ASI protocol followed by the empathic social interaction test. **b,** Change in mechanical pain thresholds after interacting for 30 min with a painful demonstrator. $n = 12$ mice per group. GH: $p = 0.0005$, Wilcoxon signed-rank test; ASI: $p = 0.0038$, paired *t*-test. **c,** Percentage decrease in mechanical pain thresholds. $n = 12$ mice per group. $p = 0.0074$, unpaired *t*-test. **d,** Baseline pain thresholds prior to social interaction. $n = 12$ mice per group. $p = 0.4943$, unpaired *t*-test. **e,** Proportion of mice exhibiting or not exhibiting contagious pain. **f-i,** Allo-licking and allo-grooming metrics during the 30-minute session. $n = 12$ mice per group. (**f**) Total duration: $p = 0.7659$, Mann-Whitney test. (**g**) Number of bouts: $p = 0.3275$, unpaired *t*-test. (**h**) Latency to first bout: $p = 0.8549$, Mann-Whitney test. (**i**) Duration per bout: $p = 0.5605$, Mann-Whitney test. **j-m,** Sniffing behavior analysis. $n = 12$ mice per group. (**j**) Total duration: $p = 0.2684$, unpaired *t*-test. (**k**) Bout number: $p = 0.1718$, Mann-Whitney test. (**l**) Latency to first bout: $p = 0.5034$, Mann-Whitney test. (**m**) Duration per bout: $p = 0.899$, Mann-Whitney test. $^{**}p < 0.01$, $^{***}p < 0.001$, n.s., no significant difference. Data are shown as means ± SEM. See Supplementary Table 1 for detailed statistics.

**Extended Data Fig. 6 SI impairs OFC neuronal activation during empathic social interaction.**

**a,** Experimental timeline for c-Fos immunostaining in GH and SI mice. **b,** Representative c-Fos immunostaining images across bregma levels in the orbitofrontal cortex (OFC), highlighting medial (mOFC), ventral (vOFC), and lateral (lOFC) subregions in GH and SI mice. Scale bar, 100 μm. **c,** Quantification of c-$Fos^+$ cells in vOFC and lOFC at different anterior-posterior coordinates. $n = 4$ mice per group. +2.80 mm: $p = 0.0013$, +2.58 mm: $p = 0.0001$, +2.34 mm: $p = 0.0062$, +2.10 mm: $p = 0.1577$. Two-way ANOVA with Bonferroni *post hoc* tests. **d,** Quantification of c-$Fos^+$ cells in the mOFC. $n = 4$ mice per group. +2.80 mm: $p = 0.4305$, +2.58 mm: $p = 0.0907$, +2.34 mm: $p = 0.0436$, +2.10 mm: $p = 0.9695$. Two-way ANOVA with Bonferroni *post hoc* tests. $^{*}p < 0.05$, $^{**}p < 0.01$, $^{***}p < 0.001$, n.s., no significant difference. Data are presented as mean ± SEM. See Supplementary Table 1 for detailed statistics.

**Extended Data Fig. 7 Effects of chemogenetic inhibition of OFC glutamatergic neurons on other behavioral parameters of the empathic social interaction test in GH mice.**

**a,** Baseline mechanical pain thresholds before the empathic social interaction in male mice. Saline: *n* = 12; CNO: *n* = 13. *p* = 0.7084, unpaired *t*-test. **b,** Proportion of mice with or without contagious pain. **c, d,** Corresponding results in female mice. *n* = 10 mice per group. (**c**) *p* = 0.3603, unpaired *t*-test. **e,** Latency to initiate allo-licking or allo-grooming behavior. Saline: *n* = 12; CNO: *n* = 13. *p* = 0.0442, Mann-Whitney test. **f,** Duration of each bout of allo-licking or allo-grooming behavior. Saline: *n* = 12; CNO: *n* = 13. *p* = 0.8518, unpaired *t*-test. **g, h,** Similar to (**e** and **f**) but for data in the female mice. *n* = 10 mice per group. (**g**) *p* = 0.0002, Mann-Whitney test. (**h**) p = 0.3369, unpaired *t*-test with Welch's correction. **i,** Total duration of sniffing behavior during a 30-min empathic social interaction session in male mice. Saline: *n* = 12; CNO: *n* = 13. *p* = 0.0025, Mann-Whitney test. **j,** Bout number of sniffing behavior. Saline: *n* = 12; CNO: *n* = 13. *p* = 0.1087, unpaired *t*-test. **k, l,** Similar to (**i** and **j**) but for data in the female mice. *n* = 10 mice per group. (**k**) *p* = 0.001, unpaired *t*-test with Welch's correction. (**l**) *p* = 0.0011, unpaired *t*-test. **m,** Latency to initiate sniffing behavior. Saline: *n* = 12 mice; CNO: *n* = 13 mice. *p* = 0.5286, Mann-Whitney test. **n,** Duration of each bout of sniffing behavior. Saline: *n* = 12; CNO: *n* = 13. *p* = 0.0787, unpaired *t*-test. **o, p,** Similar to (**m** and **n**) but for data in the female mice. *n* = 10 mice per group. (**o**) *p* = 0.4444, Mann-Whitney test. (**p**) *p* = 0.0711, unpaired *t*-test. $^*p < 0.05$, $^{**}p < 0.01$, $^{***}p < 0.001$, n.s., no significant difference. Data are shown as mean ± SEM. See Supplementary Table 1 for detailed statistics.

**Extended Data Fig. 8 Chemogenetic inhibition of OFC glutamatergic neurons does not alter anxiety-like behavior or locomotion.**

**a,** Experimental timeline for the open field test (OFT) following chemogenetic inhibition in male and female mice. **b,** Representative locomotor traces in OFT from saline-treated (dark grey) and

CNO-treated (red) male mice. **c,** Total duration spent in the center area of the arena. $n = 12$ mice per group. $p = 0.6636$, unpaired $t$-test. **d,** Distance traveled in the center area of the arena. $n = 12$ mice per group. $p = 0.811$, Mann-Whitney test. **e,** Average velocity during traveling in the arena. $n = 12$ mice per group. $p = 0.7335$, unpaired $t$-test. **f,** Total distance traveled in the arena. $n = 12$ mice per group. $p = 0.7334$, unpaired $t$-test. **g-k,** Similar to (**b-f**) but for data in the female mice. $n = 10$ mice per group. (**h**) $p = 0.6261$; (**i**) $p = 0.9774$; (**j**) $p = 0.1346$; (**k**) $p = 0.1344$. All unpaired $t$-tests. n.s., no significant difference. All data are presented as mean ± SEM. See Supplementary Table 1 for detailed statistics.

**Extended Data Fig. 9 OFC glutamatergic neuronal activity is coupled with prosocial behaviors in SI male mice.**

**a,** Timeline of fiber photometry recording of OFC glutamatergic neurons in SI male mice. **b,** Schematic (left) and representative fluorescence image (right) showing viral expression and fiber implantation targeting the OFC. Scale bar, 100 μm. **c, d,** Heatmaps of $Ca^{2+}$ activity aligned to the onset of allo-licking/grooming (**c**) and sniffing (**d**) behaviors. Each row represents changes in $Ca^{2+}$ for one trial of prosocial or sniffing behavior. The color scale indicates the range of z-score. $n = 8$ mice. **e,** Peri-event plots of Z-scored $Ca^{2+}$ signals during prosocial and sniffing behaviors. Solid lines represent the mean; shaded areas indicate SEM. $n = 8$ mice. **f,** Example trace of Z-scored $Ca^{2+}$ signals during prosocial interactions and sniffing behaviors. **g, h,** Quantification of $Ca^{2+}$ activity changes as area under the curve (AUC; **g**) and peak Z-score (**h**). $n = 8$ mice per group. (**g**) $p = 0.0114$; (**h**) $p = 0.0011$. Both unpaired $t$-tests. $^{*}p < 0.05$, $^{**}p < 0.01$, n.s., no significant difference. Data are shown as mean ± SEM. See Supplementary Table 1 for detailed statistics.

**Extended Data Fig. 10 SI alters the membrane properties and synaptic transmission of OFC glutamatergic neurons.**

**a,** Resting membrane potential of OFC glutamatergic neurons. GH: $n = 26$ neurons from 8 mice; SI: $n = 20$ neurons from 3 mice. $p = 0.0015$, unpaired *t*-test. **b, c,** Threshold (**b**) and rheobase (**c**) of action potential (AP) firing. GH: $n = 26$ neurons from 8 mice; SI: $n = 20$ neurons from 3 mice. (**b**) $p < 0.0001$, unpaired *t*-test with Welch's correction. (**c**) $p < 0.0001$, Mann-Whitney test. **d, e,** Rise tau (**d**) and decay tau (**e**) of AP. GH: $n = 26$ neurons from 8 mice; SI: $n = 20$ neurons from 3 mice. (**d**) $p = 0.0035$, unpaired *t*-test. (**e**) $p = 0.0977$, Mann-Whitney test. **f,** Quantification of AP amplitude. GH: $n = 26$ neurons from 8 mice; SI: $n = 20$ neurons from 3 mice. $p = 0.0764$, unpaired *t*-test. **g,** Input resistance of OFC glutamatergic neurons. GH: $n = 26$ neurons from 8 mice; SI: $n = 20$ neurons from 3 mice. $p = 0.119$, Mann-Whitney test. **h, i,** Averaged sEPSC/sIPSC amplitude (**h**) and frequency (**i**) of OFC glutamatergic neurons. GH: $n = 18$ neurons from 6 mice; SI: $n = 22$ neurons from 4 mice. (**h**) $p = 0.0171$, unpaired *t*-test. (**i**) $p = 0.0168$, Mann-Whitney test. $^{*}p < 0.05$, $^{**}p < 0.01$, $^{***}p < 0.001$, n.s., no significant difference. Data are presented as means ± SEM. See Supplementary Table 1 for statistical details.

**Extended Data Fig. 11 Effects of chemogenetic activation of OFC glutamatergic neurons on other behavioral parameters of the empathic social interaction test in SI mice.**

**a,** Baseline mechanical pain thresholds. Saline: $n = 8$; CNO: $n = 11$. $p = 0.2438$, unpaired *t*-test. **b,** Proportion of male mice exhibiting or not exhibiting contagious pain. **c, d,** Similar to (**a** and **b**) but in female mice. $n = 11$ mice per group. $p = 0.985$, unpaired *t*-test. **e, f,** Latency to initiate allo-licking/grooming (**e**) and duration per bout (**f**) in male mice. Saline: $n = 8$; CNO: $n = 11$. (**e**) $p = 0.5597$, (**f**) $p = 0.5298$. Both Mann-Whitney tests. (**g** and **h**) Similar to (**e** and **f**) but in female mice.

*n* = 11 mice per group. (**g**) $p = 0.2106$, Mann-Whitney test. (**h**) $p = 0.3128$, unpaired *t*-test. **i, j,** Total duration (**i**) and bout count (**j**) of sniffing behavior in male mice. Saline: $n = 8$; CNO: $n = 11$. (**i**) $p = 0.0005$, (**j**) $p = 0.0235$. Both unpaired *t*-tests. **k, l,** Similar to (**i** and **j**) but in female mice. $n = 11$ mice per group. (**k**) $p = 0.104$, (**l**) $p = 0.1625$. Both Mann-Whitney tests. **m, n,** Latency to initiate sniffing behavior (**m**) and duration of each bout (**n**) in male mice. Saline: $n = 8$; CNO: $n = 11$. (**m**) $p = 0.8558$, Mann-Whitney test. (**n**) $p = 0.0002$, unpaired *t*-test. **o, p,** Similar to (**m** and **n**) but in female mice. $n = 11$ mice per group. (**o**) $p = 0.2485$, (**p**) $p = 0.3074$. Both Mann-Whitney tests. *$p < 0.05$, ***$p < 0.001$, n.s., no significant difference. Data are presented as means ± SEM. See Supplementary Table 1 for statistical details.

**Extended Data Fig. 12 Whole-brain input mapping of social interaction-TRAPed OFC neurons and anterograde tracing of the VM→OFC pathway.**

**a,** Timeline illustrating the strategy for whole-brain input mapping to social interaction-TRAPed OFC neurons. **b,** Schematic of rabies virus-mediated monosynaptic retrograde tracing of presynaptic inputs to TRAPed OFC neurons. **c,** Representative image showing dual-labeled OFC starter neurons infected by both the helper virus ($GFP^+$) and rabies virus ($DsRed^+$). Scale bar, 100 μm. **d,** Quantification of DsRed-labeled neurons projecting to the OFC in selected brain regions. $n = 5$ mice. M1/M2: motor cortex; IC: insular cortex; S1/S2: somatosensory cortex; Pir: piriform cortex; MS: medial septal nucleus; SI: substantia innominate; PVT: paraventricular thalamic nucleus; PT: paratenial thalamic nucleus; AM: anteromedial thalamic nucleus; MD: mediodorsal thalamic nucleus; Re: reuniens thalamic nucleus; VL: ventrolateral thalamic nucleus; VM: ventromedial thalamic nucleus; BLA: basolateral amygdaloid nucleus; CM: central medial thalamic nucleus; Sub: submedius thalamic nucleus; Po: posterior thalamic nucleus; PF:

parafascicular thalamic nucleus; Ect: ectorhinal cortex; PRh: perirhinal cortex. **e,** Representative images of DsRed-labeled neurons in selected brain regions. Scale bars: 100 μm. **f,** Schematic of the viral tracing strategy to identify OFC glutamatergic neurons receiving direct input from VM. **g,** Representative immunofluorescence showing EGFP-labeled glutamatergic cells, mCherry-labeled VM-innervated cells, and merged signals (yellow, white arrowheads). Scale bars: 100 μm. **h,** Pie chart indicating that a substantial proportion of VM-innervated OFC neurons are glutamatergic. $n = 4$ mice. Data are presented as means ± SEM.

**Extended Data Fig. 13 Chemogenetic inhibition of VM→OFC projection alters empathic interaction behavior without affecting anxiety-like traits.**

**a,** Latency to initiate allo-licking/grooming behavior. $n = 13$ mice per group. $p = 0.0221$, Mann-Whitney test. **b,** Duration of each bout of allo-licking/grooming behavior. $n = 13$ mice per group. $p = 0.7445$, unpaired $t$-test. **c, d,** Latency to initiate sniffing behavior (**c**) and duration of each bout (**d**). $n = 13$ mice per group. (**c**) $p = 0.2266$, Mann-Whitney test. (**d**) $p = 0.0591$, unpaired $t$-test. **e, f,** Timeline (**e**) and schematic (**f**) for OFT following chemogenetic inhibition of the VM→OFC pathway in GH male mice. **g,** Representative OFT traces comparing saline- (dark grey) and CNO-treated (green) mice. **h-k,** Total time spent in the center (**h**), distance traveled in the center (**i**), average velocity (**j**) and total distance traveled in the arena (**k**). $n = 14$ mice per group. (**h**) $p = 0.3648$, unpaired $t$-test. (**i**) $p = 0.1139$, Mann-Whitney test. (**j**) $p = 0.1049$, unpaired $t$-test. (**k**) $p = 0.3483$, unpaired $t$-test. $^*p < 0.05$, n.s., no significant difference. Data are presented as means ± SEM. See Supplementary Table 1 for statistical details.

**Extended Data Fig. 14 Chemogenetic inhibition of glutamatergic projections from the submedius thalamic nucleus (Sub) to the OFC does not affect contagious pain or prosocial behaviors.**

**a,** Experimental timeline illustrating the chemogenetic suppression of Sub→OFC projections during empathic social interactions in GH male mice. **b,** Schematic (left) and representative image (right) showing viral injection sites in the Sub and OFC. Scale bar, 100 μm. **c,** Change in mechanical pain thresholds. Saline: $n = 10$; CNO: $n = 12$. Saline: $p = 0.0039$, Wilcoxon matched-pairs signed-rank test; CNO: $p = 0.0009$, paired $t$-test. **d,** Percentage decrease in mechanical pain thresholds. Saline: $n = 10$; CNO: $n = 12$. $p = 0.4562$, Mann-Whitney test. **e,** Baseline mechanical pain thresholds. Saline: $n = 10$; CNO: $n = 12$. $p = 0.4667$, unpaired $t$-test. **f,** Proportion of mice with or without contagious pain. **g,** Total duration of allo-licking and allo-grooming. Saline: $n = 10$; CNO: $n = 12$. $p = 0.1063$, unpaired $t$-test with Welch's correction. **h,** Number of bouts of allo-licking and allo-grooming. Saline: $n = 10$; CNO: $n = 12$. $p = 0.1122$, unpaired $t$-test. **i,** Latency to initiate allo-licking or allo-grooming. Saline: $n = 10$; CNO: $n = 12$. $p = 0.0237$, Mann-Whitney test. **j,** Duration of individual bouts of allo-licking or allo-grooming. Saline: $n = 10$; CNO: $n = 12$. $p = 0.8718$, Mann-Whitney test. **k-n,** Sniffing behavior analyses corresponding to panels (**g**-**j**). Saline: $n = 10$; CNO: $n = 12$. (**k**) $p = 0.372$, Mann-Whitney test; (**l**) $p = 0.2344$, unpaired $t$-test; (**m**) $p = 0.4958$, unpaired $t$-test; (**n**) $p = 0.381$, Mann-Whitney test. $^*p < 0.05$, $^{**}p < 0.01$, $^{***}p < 0.001$, n.s., no significant difference. Data are expressed as means ± SEM. See Supplementary Table 1 for detailed statistics.

**Extended Data Fig. 15 OFC glutamatergic neurons receive local inhibition from OFC interneurons, which are innervated by VM glutamatergic projections.**

**a,** Representative traces of light-evoked IPSCs recorded in OFC glutamatergic neurons under baseline conditions (aCSF), in the presence of TTX, and TTX combined with 4-AP. **b,** Quantification of IPSC amplitudes from (A). *n* = 7 neurons from 3 mice. aCSF vs. TTX: *p* = 0.0011; TTX vs. TTX+4-AP: *p* = 0.262. One-way RM ANOVA with Bonferroni *post hoc* tests. **c, d,** Representative traces (**c**) and quantification (**d**) of paired-pulse ratios (PPRs) of light-induced IPSCs in OFC glutamatergic neurons from GH (black) and SI (red) mice. GH: *n* = 20 neurons from 5 mice; SI: *n* = 22 neurons from 4 mice. 25 ms: *p* = 0.4383, 50 ms: *p* = 0.7916, 200 ms: *p* = 0.3986, 500 ms: *p* = 0.2139, All Mann-Whitney tests; 100 ms: *p* = 0.9099, unpaired *t*-test. **e,** Schematic depicting VM glutamatergic input and local inhibitory circuitry in the OFC. **f, g,** Representative traces (**f**) and quantification (**g**) of light-evoked EPSC amplitudes in OFC glutamatergic neurons under aCSF, PTX, and CNQX + APV conditions. *n* = 7 neurons from 3 mice. aCSF *vs*. PTX: *p* = 0.0126; PTX *vs*. CNQX + APV: *p* = 0.0011. One-way RM ANOVA with Bonferroni *post hoc* tests. **h, i,** Representative traces (**h**) and quantification (**i**) of light-induced IPSCs in OFC glutamatergic neurons under aCSF, CNQX + APV, and PTX conditions. *n* = 5 neurons from 3 mice. aCSF *vs*. CNQX + APV: *p* = 0.0054; PTX *vs*. CNQX + APV: *p* > 0.9999. One-way RM ANOVA with Bonferroni *post hoc* tests. **p* < 0.05, ***p* < 0.01, n.s., no significant difference. Data are presented as means ± SEM. See Supplementary Table 1 for detailed statistics.

**Extended Data Fig. 16 Quality control of single-nucleus RNA-sequencing data from the OFC of GH and SI mice.**

**a,** Total number of efficient cells obtained from the OFC of GH and SI mice. *n* = 3 mice per group. *p* = 0.2606, unpaired *t*-test with Welch's correction. **b,** Distribution of log-transformed gene counts per cell across individual mice. **c,** t-SNE plot showing cell cluster distributions from GH and SI mice. **d,** Total UMI counts per cell in GH and SI mice. $n = 3$ mice per group. $p = 0.0529$, unpaired $t$-test. **e,** Distribution of log-transformed UMI counts per cell across individual mice. **f,** t-SNE visualization of transcriptional activity across identified cell clusters in GH and SI mice. n.s., no significant difference. Data are expressed as means ± SEM. See Supplementary Table 1 for detailed statistics.

**Extended Data Fig. 17 Molecular signatures and cell type-specific gene expression profiles across OFC cell clusters.**

**a,** Molecular characteristics of identified OFC clusters, indicated by the proportion of cells expressing specific marker genes (circle size) and the corresponding average expression levels (color scale). **b,** Heatmap illustrating differential expression of selected marker genes across identified cell clusters. Each vertical line represents a distinct cell cluster.

**Extended Data Fig. 18 Gene enrichment analysis and *Grik3* expression patterns in OFC excitatory neurons.**

**a,** KEGG pathway enrichment analysis of differentially expressed genes in OFC excitatory neurons from GH *vs.* SI mice. **b,** Gene set enrichment analysis (GSEA) highlighting differences in long-term potentiation and glutamatergic synapse pathways between GH and SI groups. **c,** Lollipop plot displaying *Grik3* expression across all identified OFC cell clusters. **d,** Quantitative comparison of *Grik3* expression in excitatory neurons of GH and SI mice. **e,** Heatmap showing expression levels of genes involved in glutamatergic transmission and synaptic plasticity in OFC excitatory neurons across both conditions.

**Extended Data Fig. 19 *Grik3* knockdown in OFC glutamatergic neurons does not alter intrinsic electrophysiological properties, anxiety-like behavior, or locomotion in GH mice.**

**a,** Resting membrane potential of OFC glutamatergic neurons. WT: $n = 25$ neurons from 4 mice; KD: $n = 30$ neurons from 4 mice. $p = 0.4692$, unpaired *t*-test. **b,** Rheobase of AP firing. WT: $n = 25$ neurons from 4 mice; KD: $n = 30$ neurons from 4 mice. $p = 0.3665$, Mann-Whitney test. **c-e,** Rise tau (**c**), decay tau (**d**) and amplitude (**e**) of AP. WT: $n = 25$ neurons from 4 mice; KD: $n = 30$ neurons from 4 mice. (**c**) $p = 0.4396$, unpaired *t*-test. (**d**) $p = 0.4653$, Mann-Whitney test. (**e**) $p = 0.834$, Mann-Whitney test. **f,** Latency to initiate allo-licking or allo-grooming. $n = 10$ mice per group. $p = 0.3622$, Mann-Whitney test. **g,** Duration of each allo-licking or allo-grooming bout. $n = 10$ mice per group. $p = 0.5872$, unpaired *t*-test. **h, i,** Latency to initiate sniffing behavior (**h**) and duration of each bout (**i**). $n = 10$ mice per group. (**h**) $p = 0.7839$, unpaired *t*-test. (**i**) $p = 0.0048$, unpaired *t*-test with Welch's correction. **j,** Experimental timeline for the OFT following *Grik3* knockdown in OFC glutamatergic neurons in GH mice. **k,** Schematic showing viral injection targeting the OFC. **l,** Representative open field trajectories from WT (dark grey) and KD (blue) male mice. **m-p,** Total time spent in the center (**m**), distance traveled in the center (**n**), average velocity (**o**) and total distance traveled in the arena (**p**). $n = 10$ mice per group. (**m**) $p = 0.0928$, unpaired *t*-test with Welch's correction. (**n**) $p = 0.4905$, (**o**) $p = 0.9103$, (**p**) $p = 0.9103$, all unpaired *t*-tests. $^{**}p < 0.01$, n.s., no significant difference. Data are presented as mean ± SEM. See Supplementary Table 1 for detailed statistics.

**Supplementary videos**

Video S1: Representative examples of allo-licking and allo-grooming behaviors initiated by observer mice toward painful demonstrators.

Video S2: Video comparison showing intact prosocial behaviors in GH observers, contrasted with their absence in SI mice.

Video S3: Real-time fiber photometry recording of OFC glutamatergic neuron activity in a GH observer during interaction with a painful demonstrator.

Video S4: Chemogenetic inhibition of OFC neurons abolished prosocial behaviors in GH mice, whereas activation of OFC neurons restored prosocial responses in SI mice.

Extended Data Fig. 1

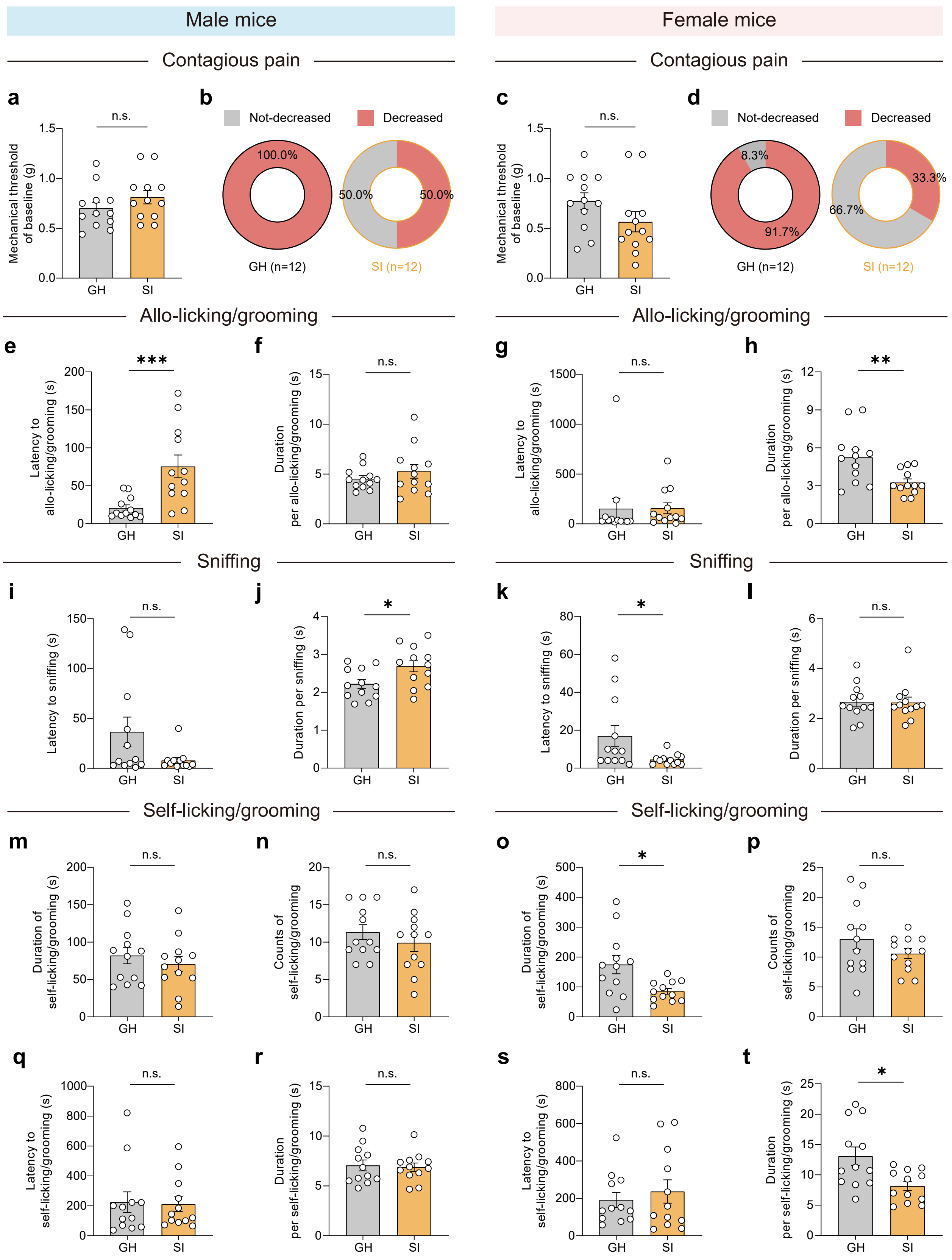

Extended Data Fig. 2

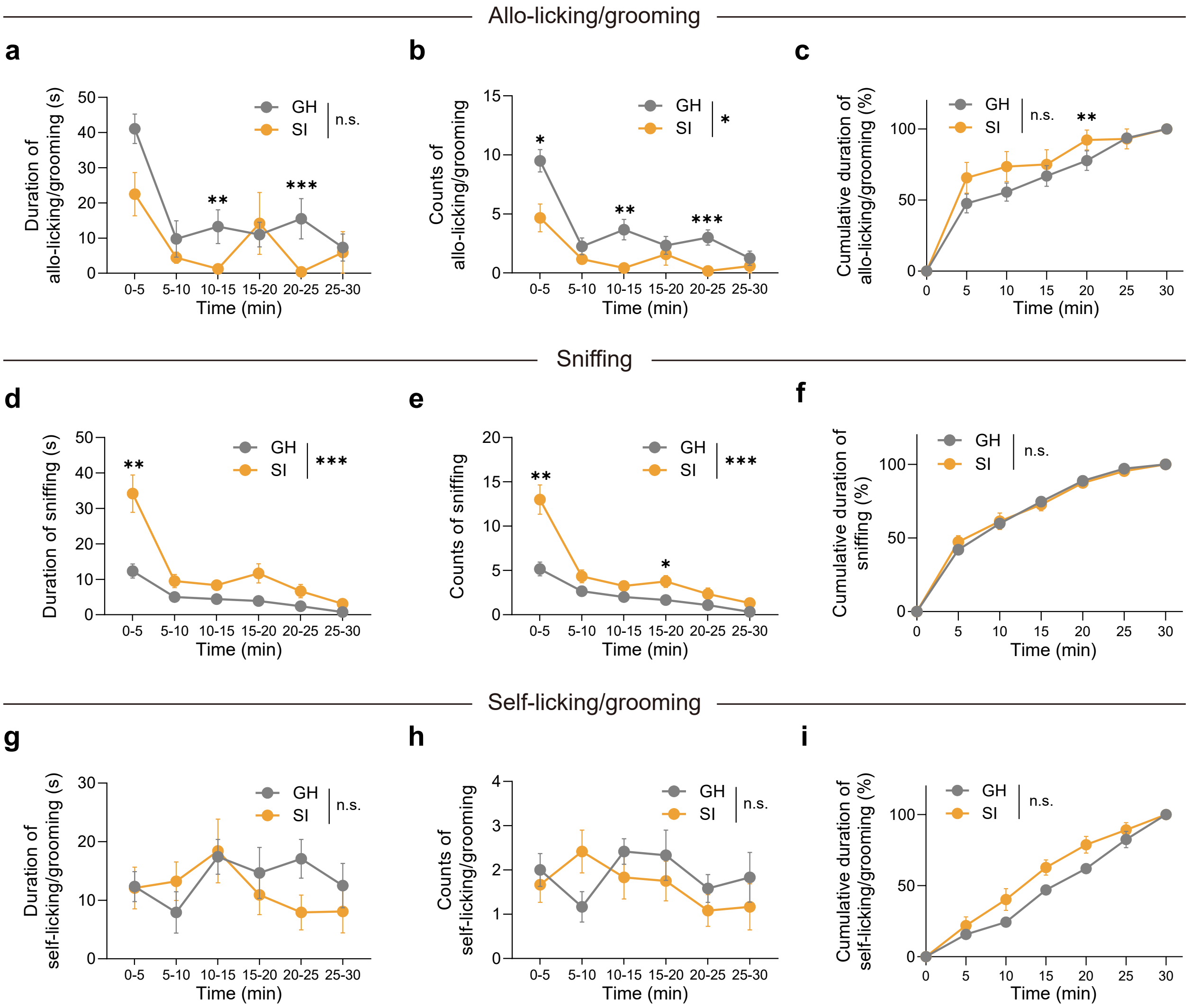

### Extended Data Fig. 3

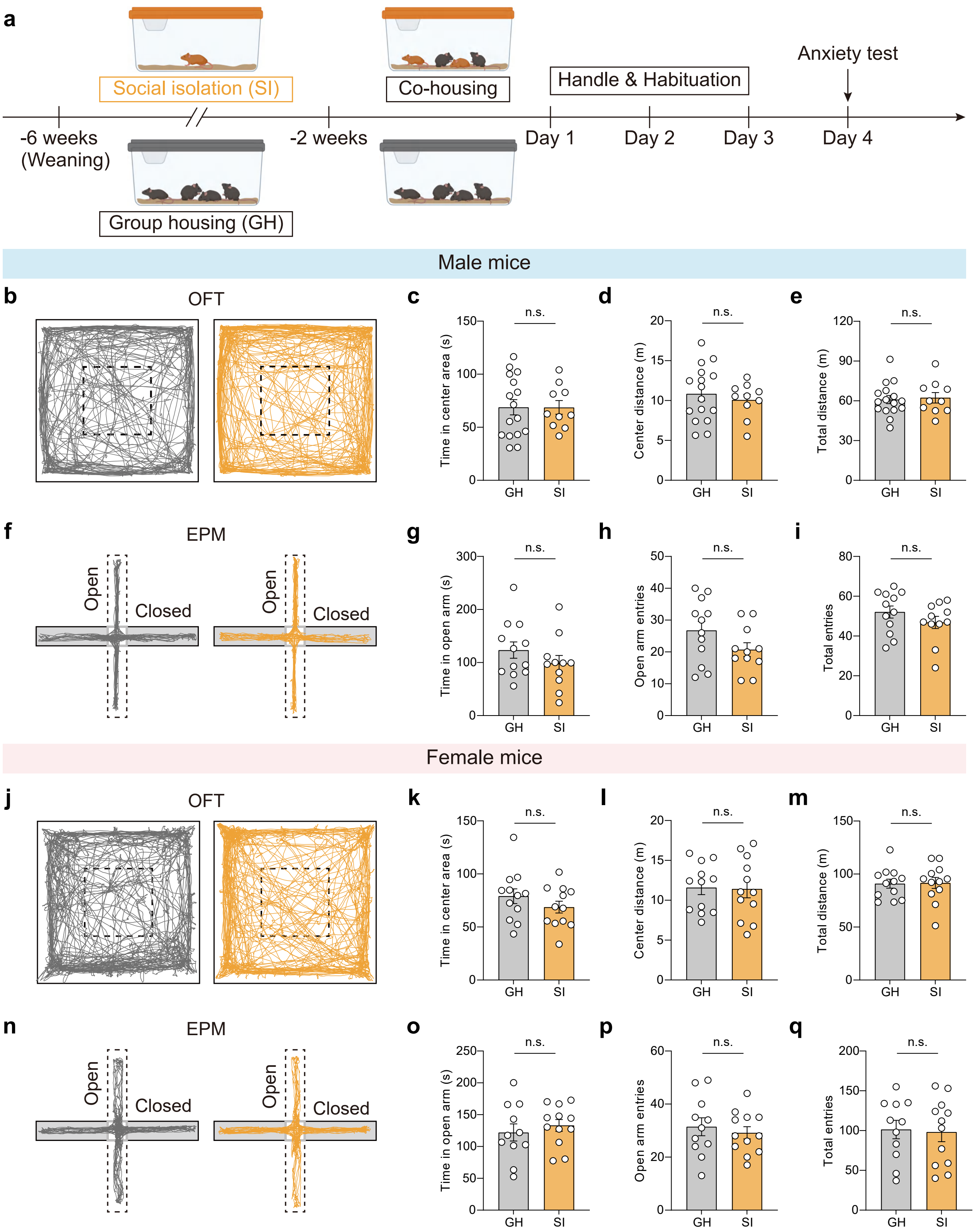

### Extended Data Fig. 4

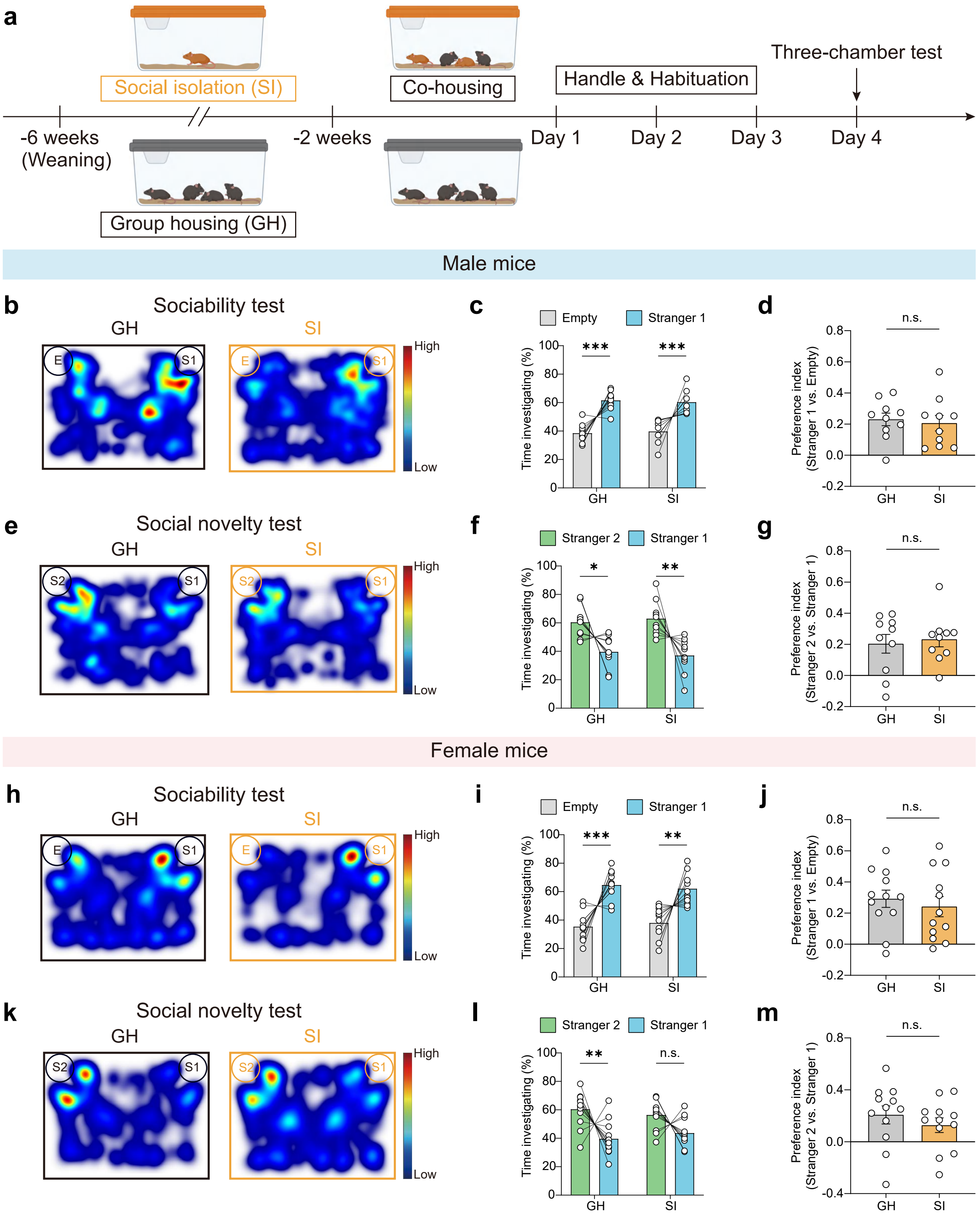

### Extended Data Fig. 5

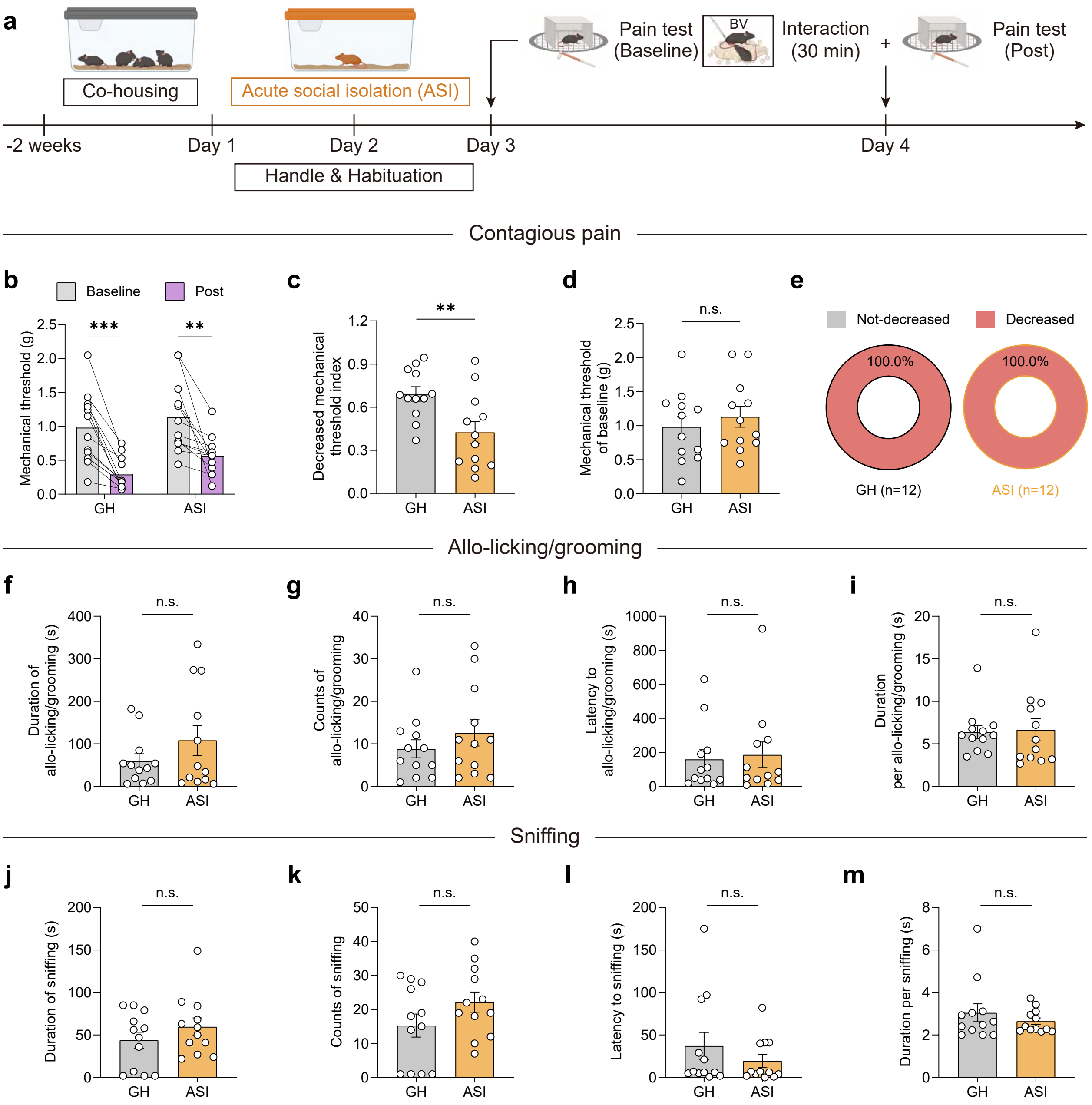

### Extended Data Fig. 6

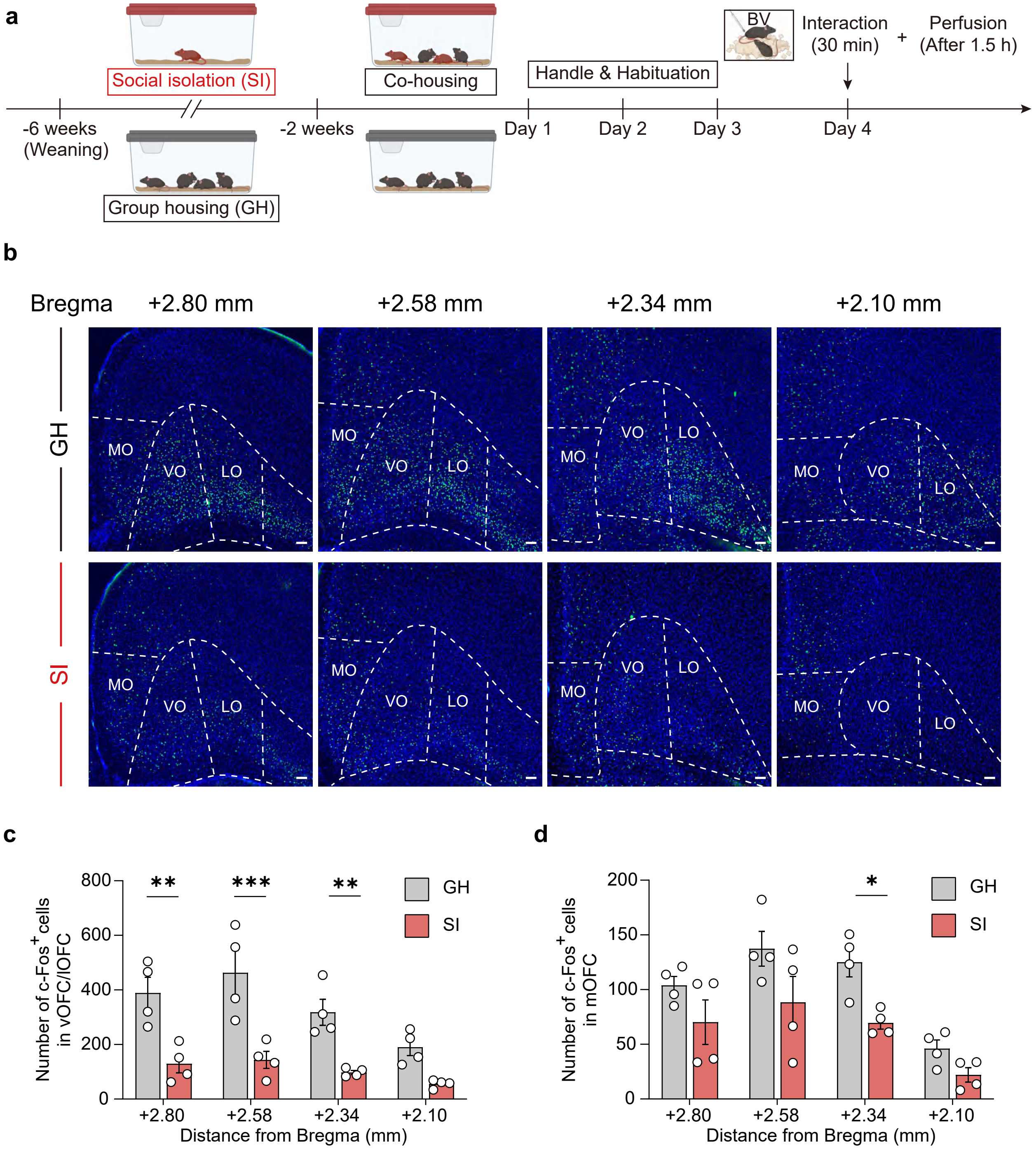

### Extended Data Fig. 7

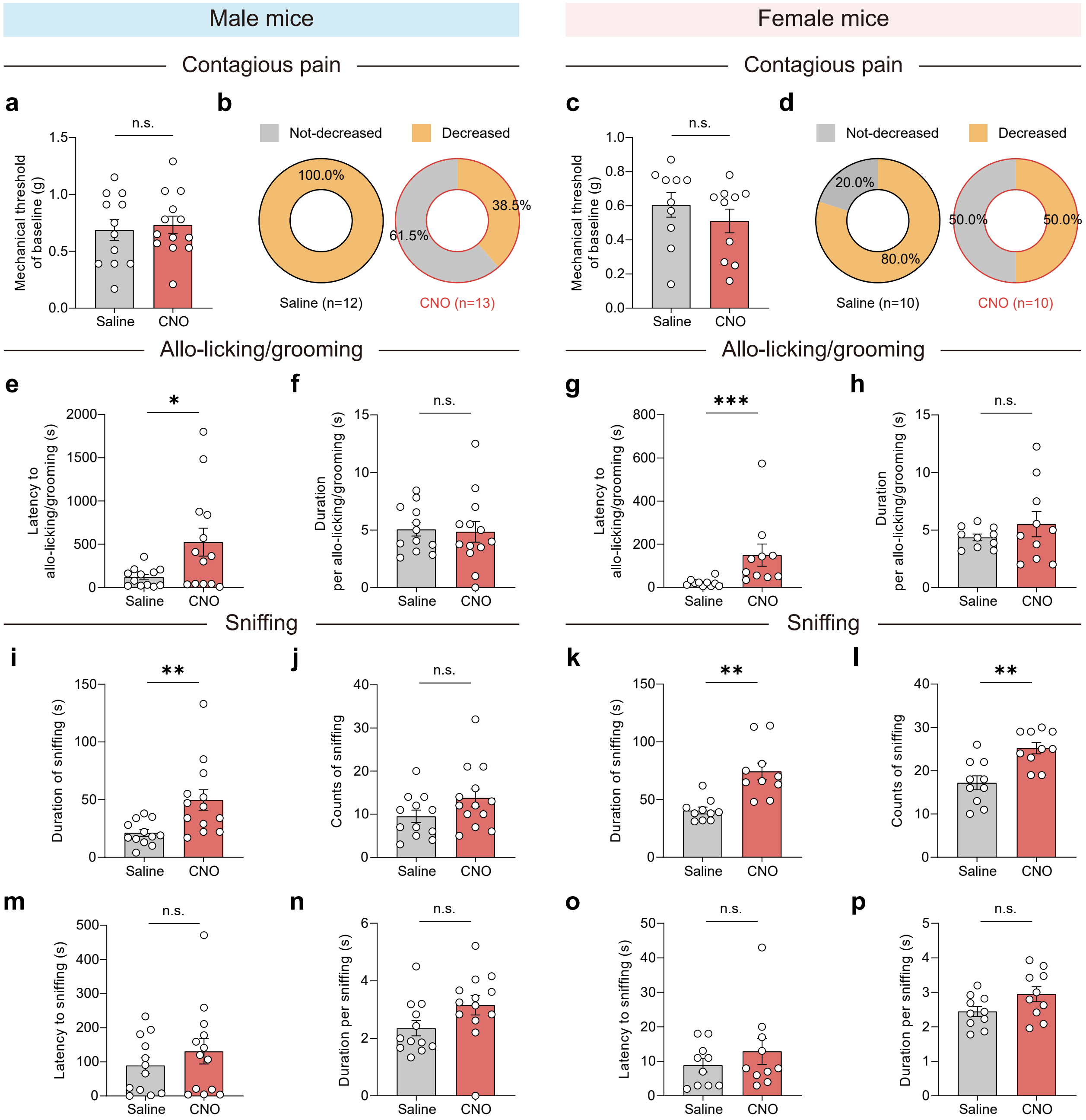

### Extended Data Fig. 8

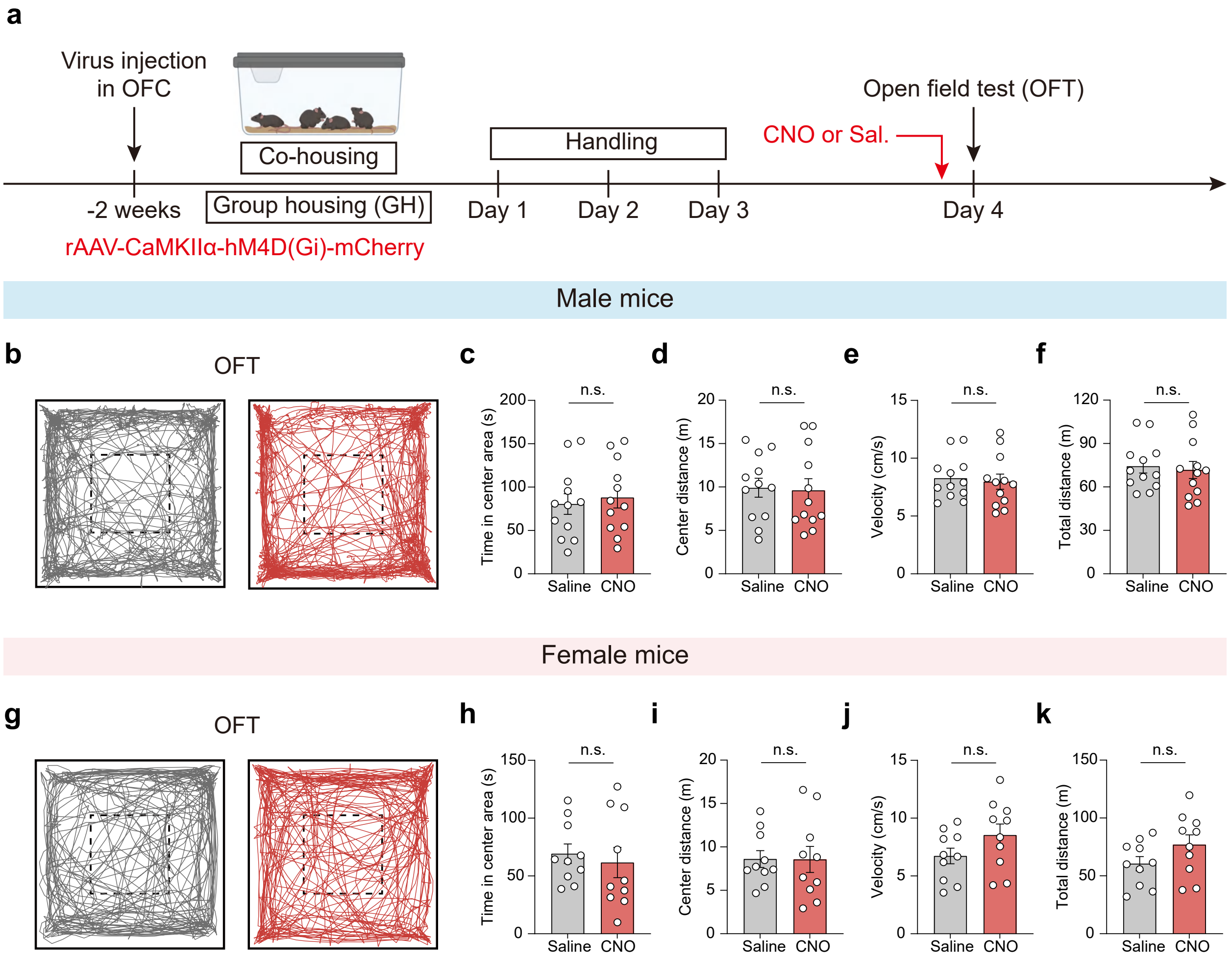

### Extended Data Fig. 9

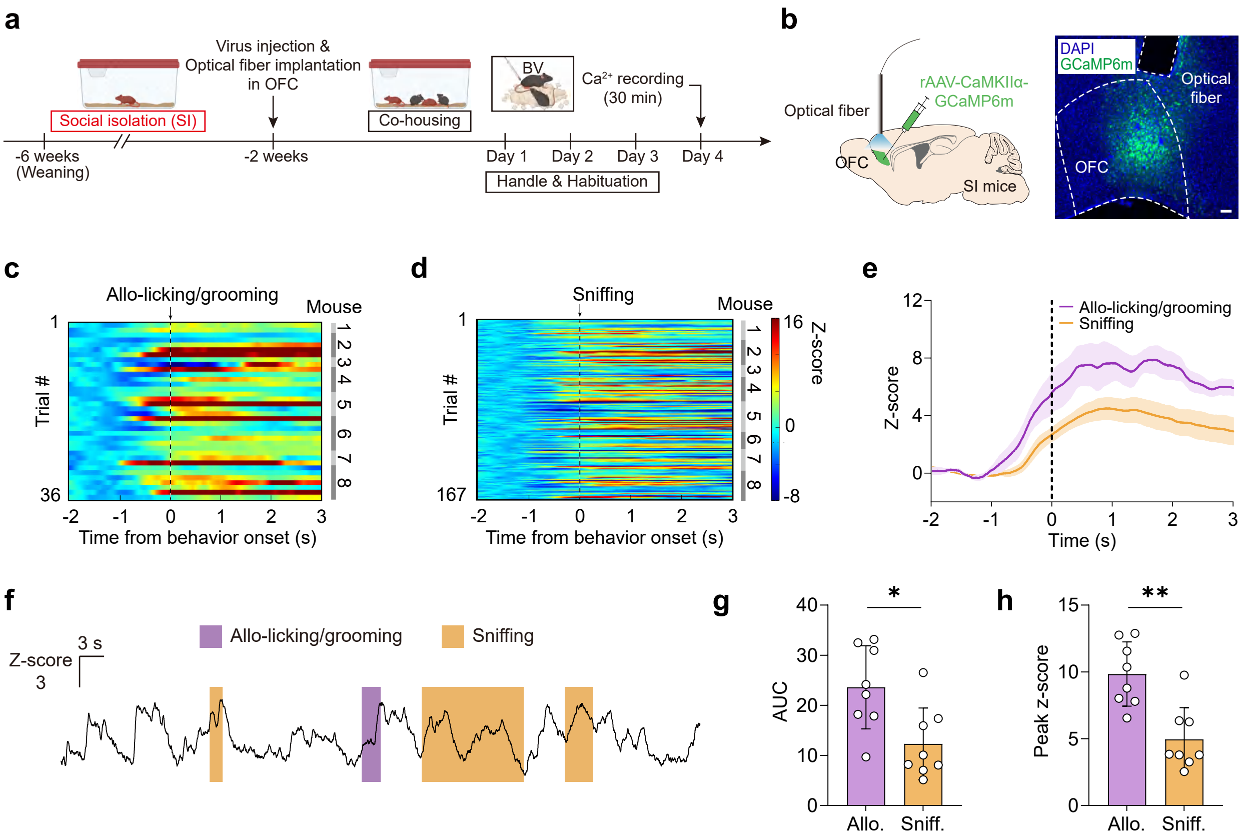

### Extended Data Fig. 10

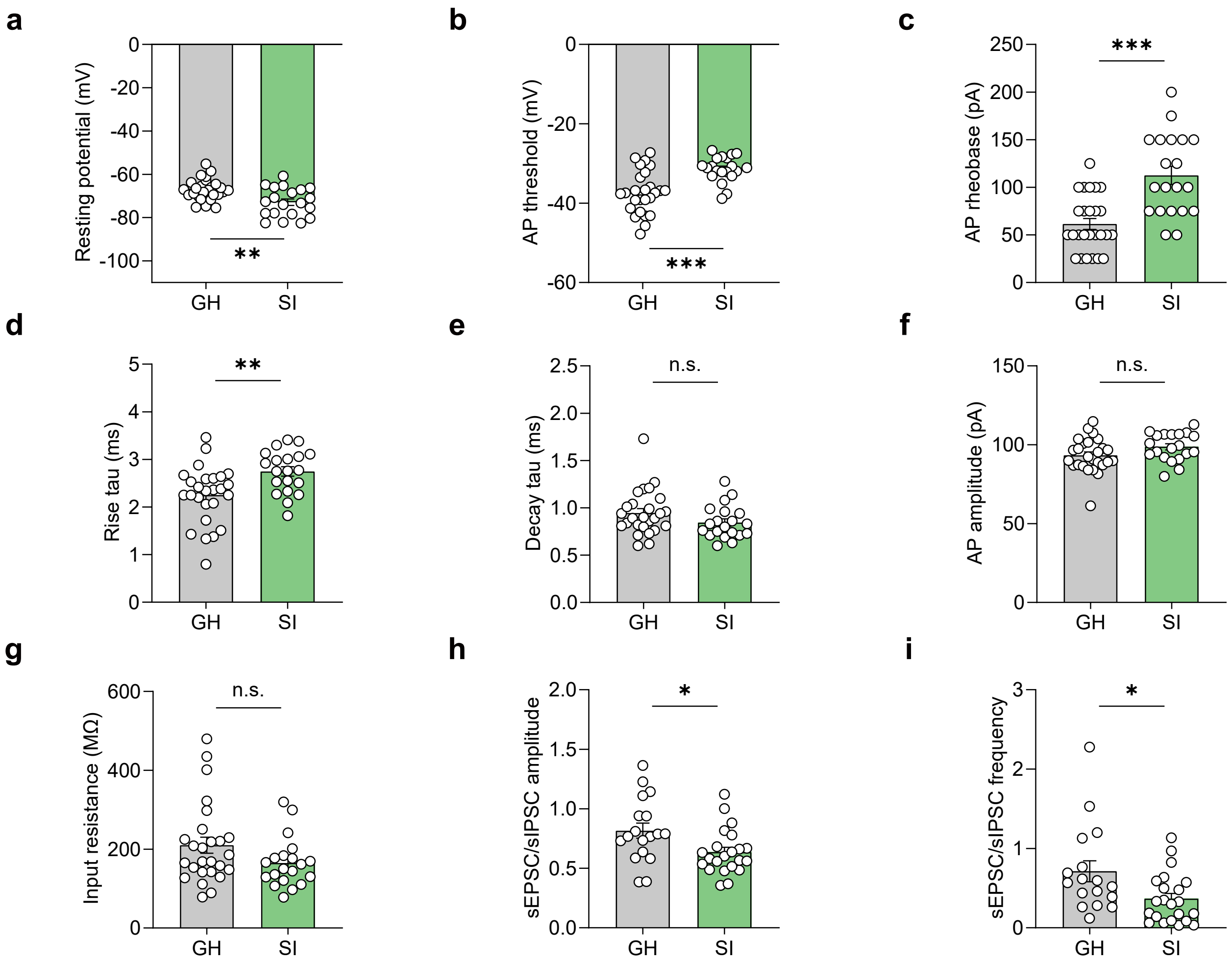

### Extended Data Fig. 11

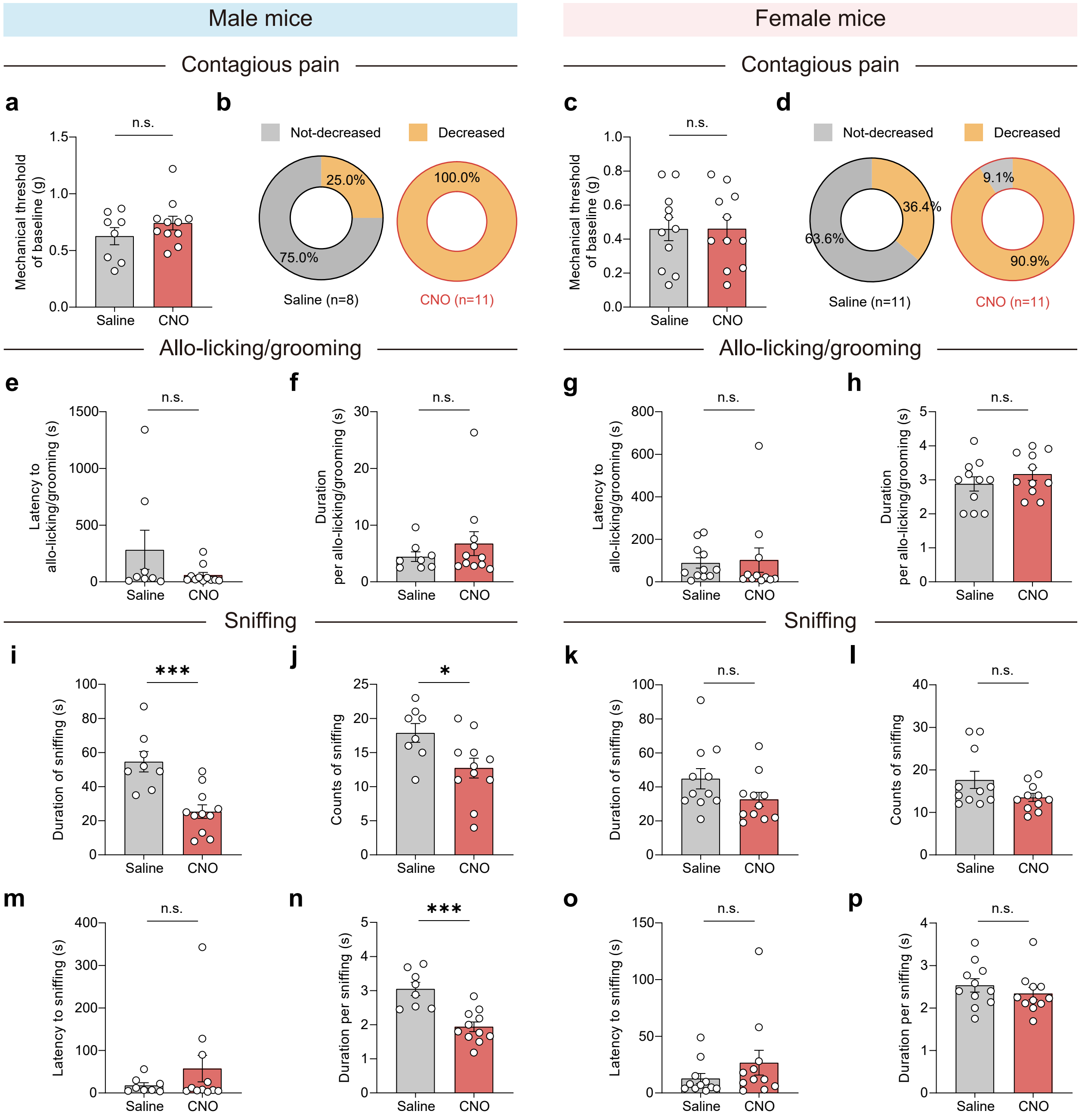

### Extended Data Fig. 12

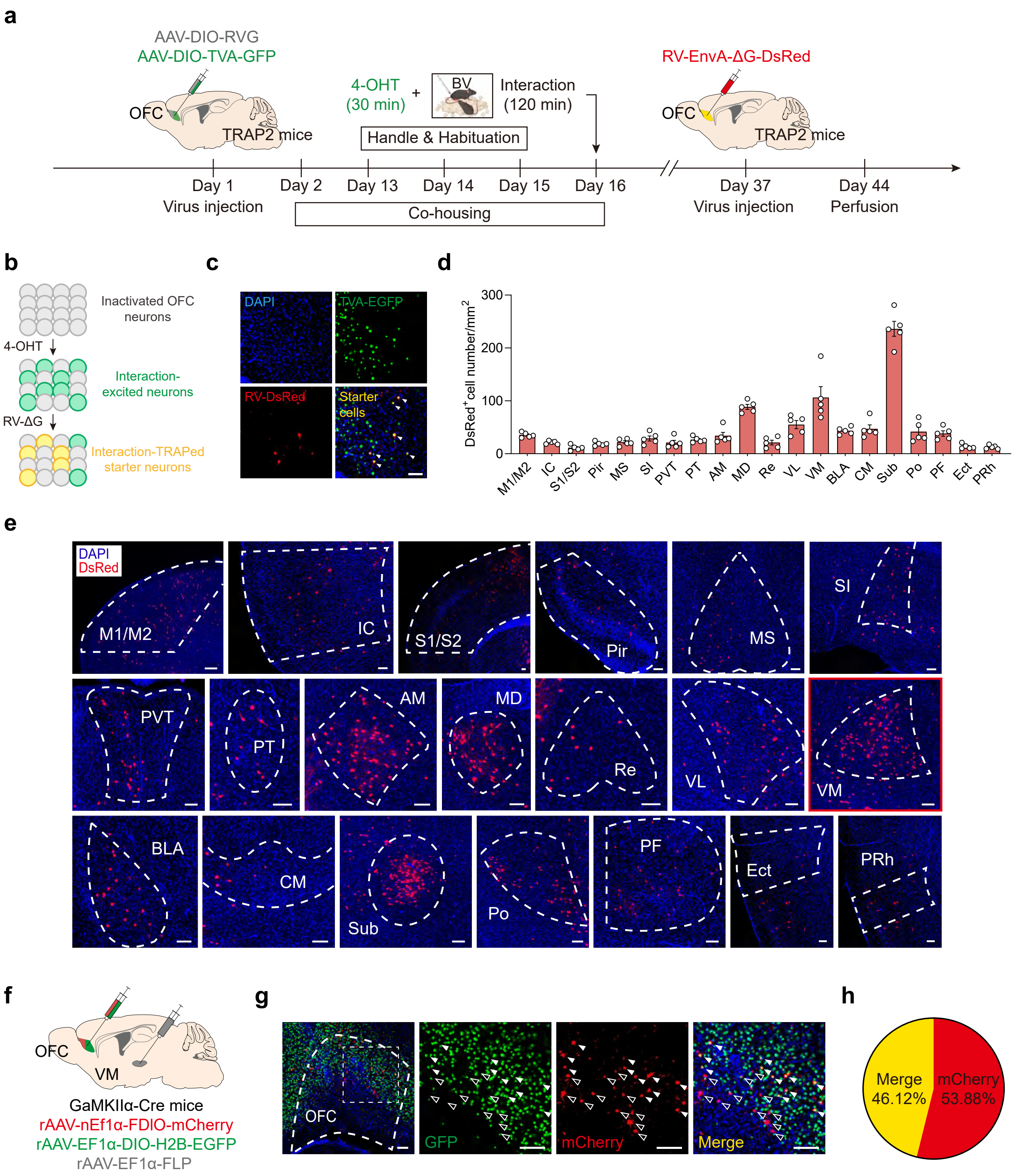

### Extended Data Fig. 13

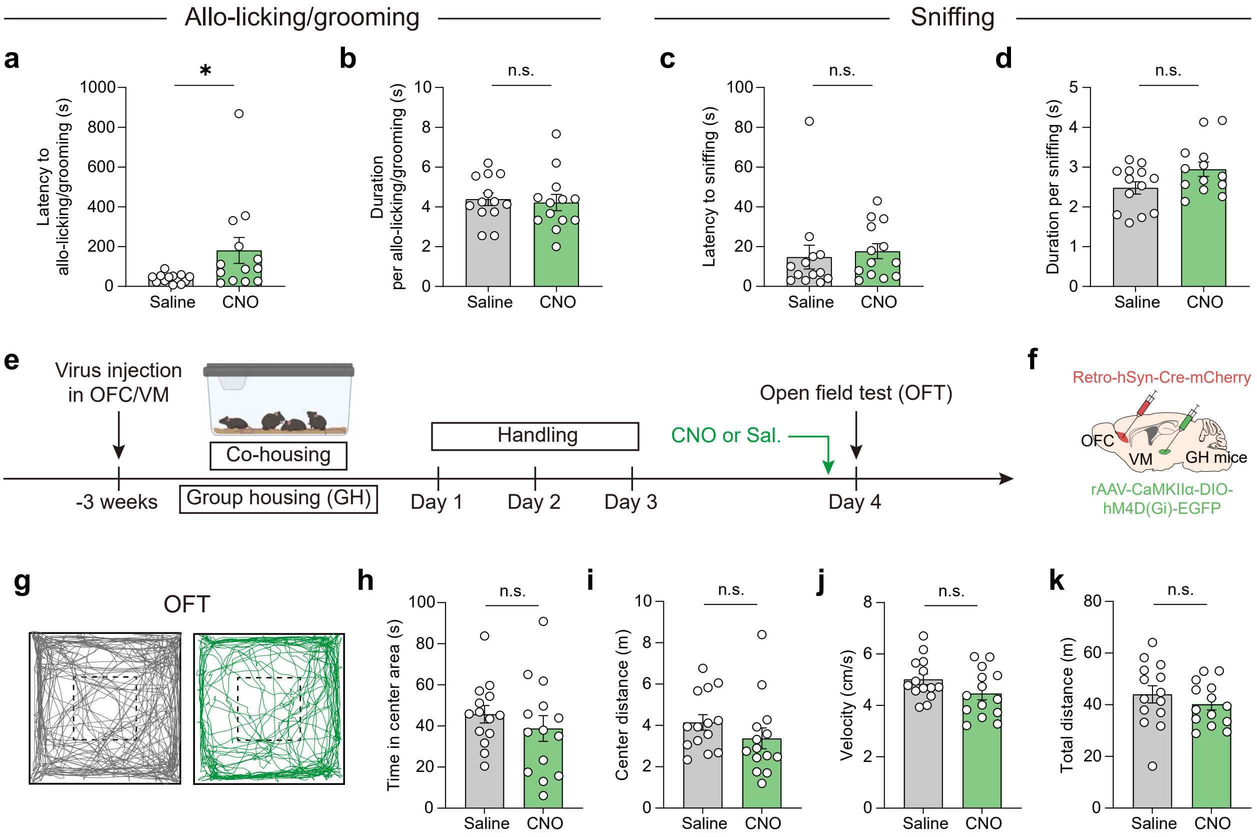

### Extended Data Fig. 14

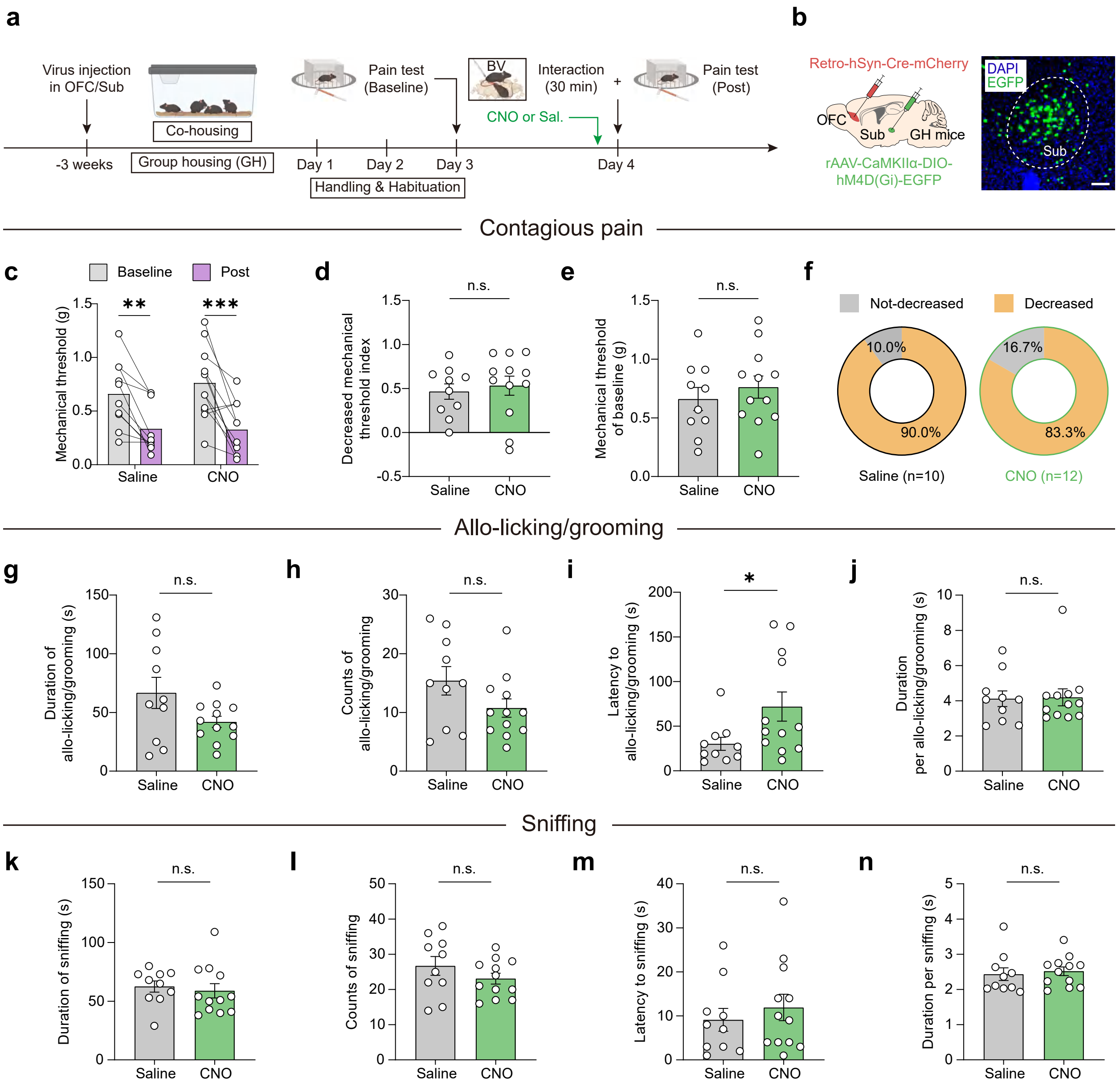

### Extended Data Fig. 15

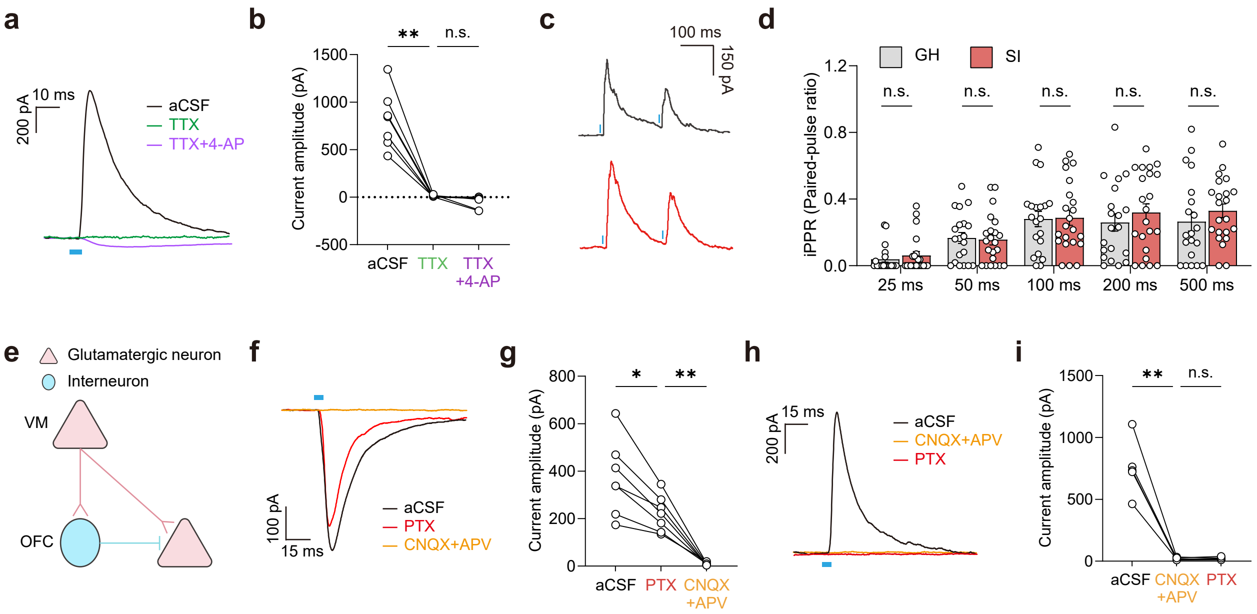

### Extended Data Fig. 16

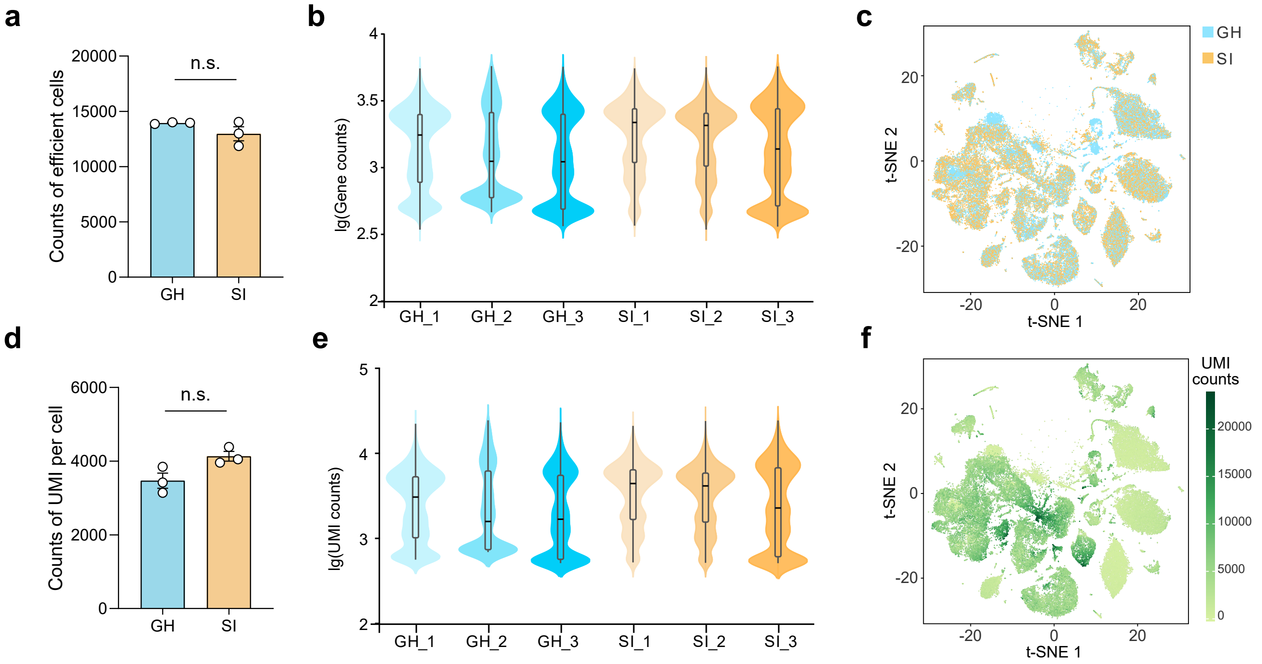

### Extended Data Fig. 17

**a**

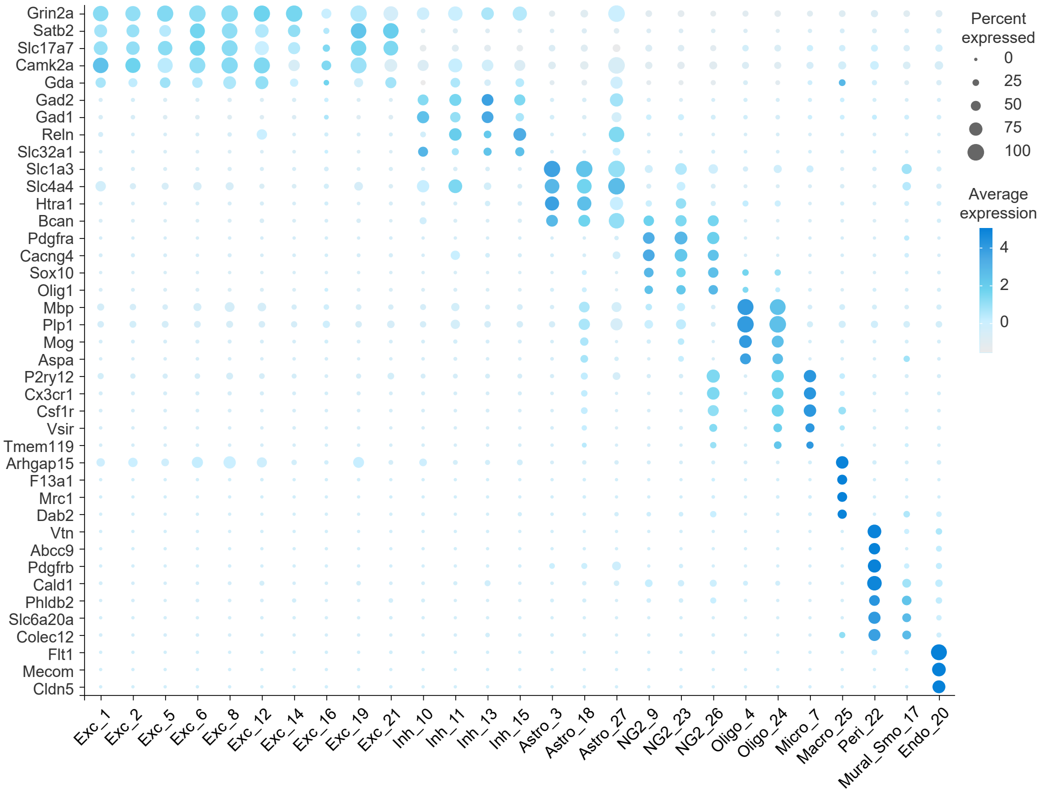


**b**

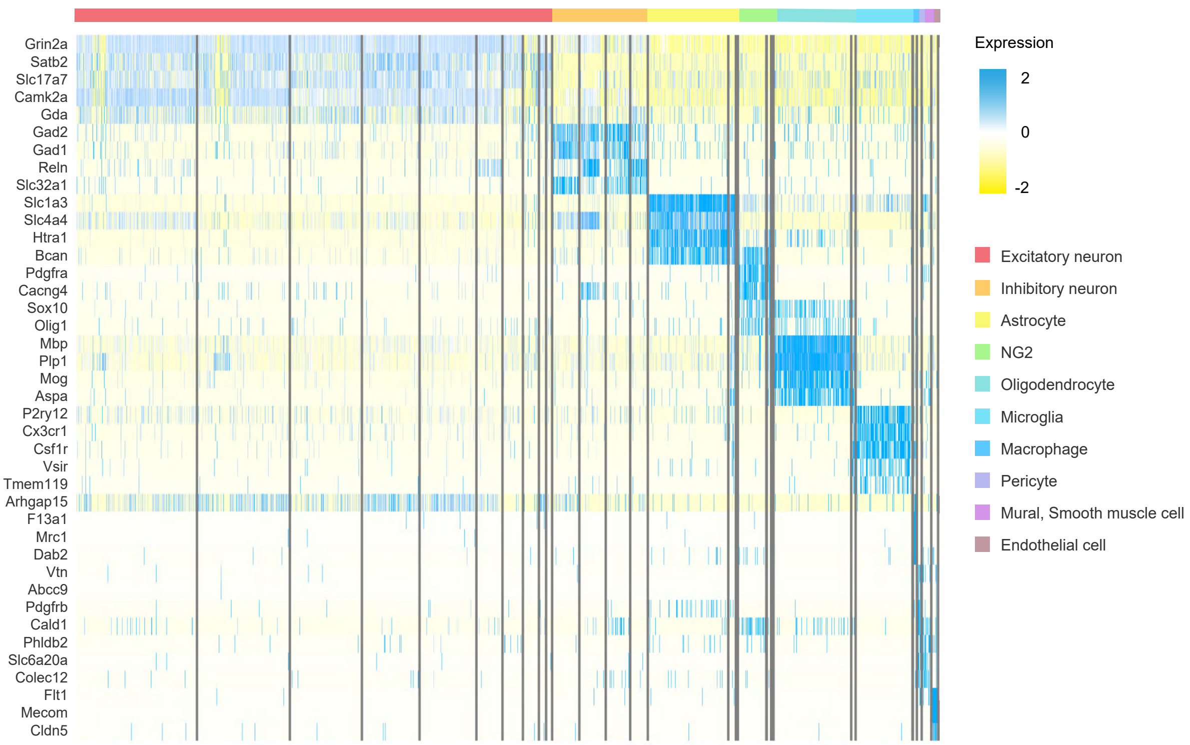

### Extended Data Fig. 18

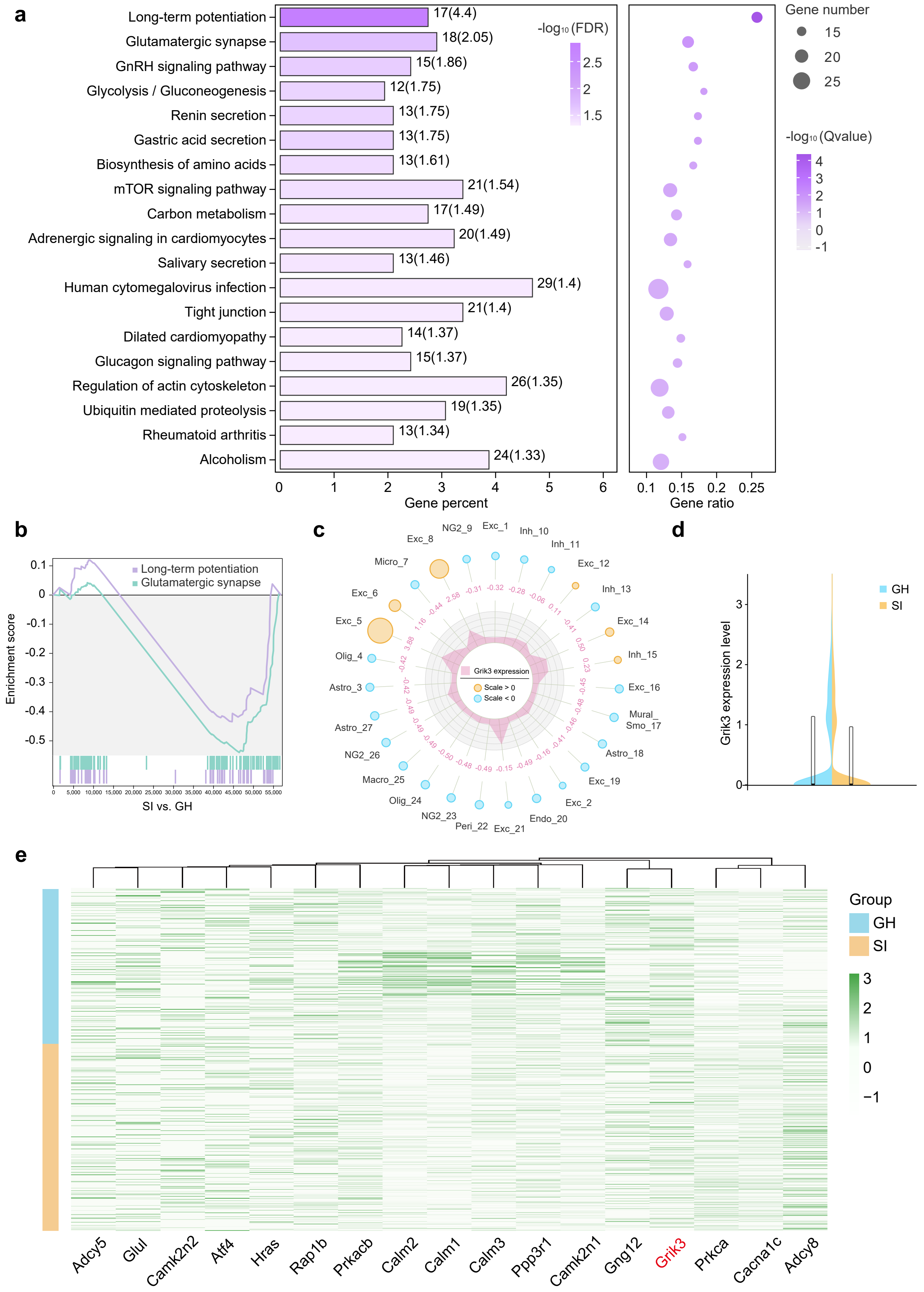

### Extended Data Fig. 19

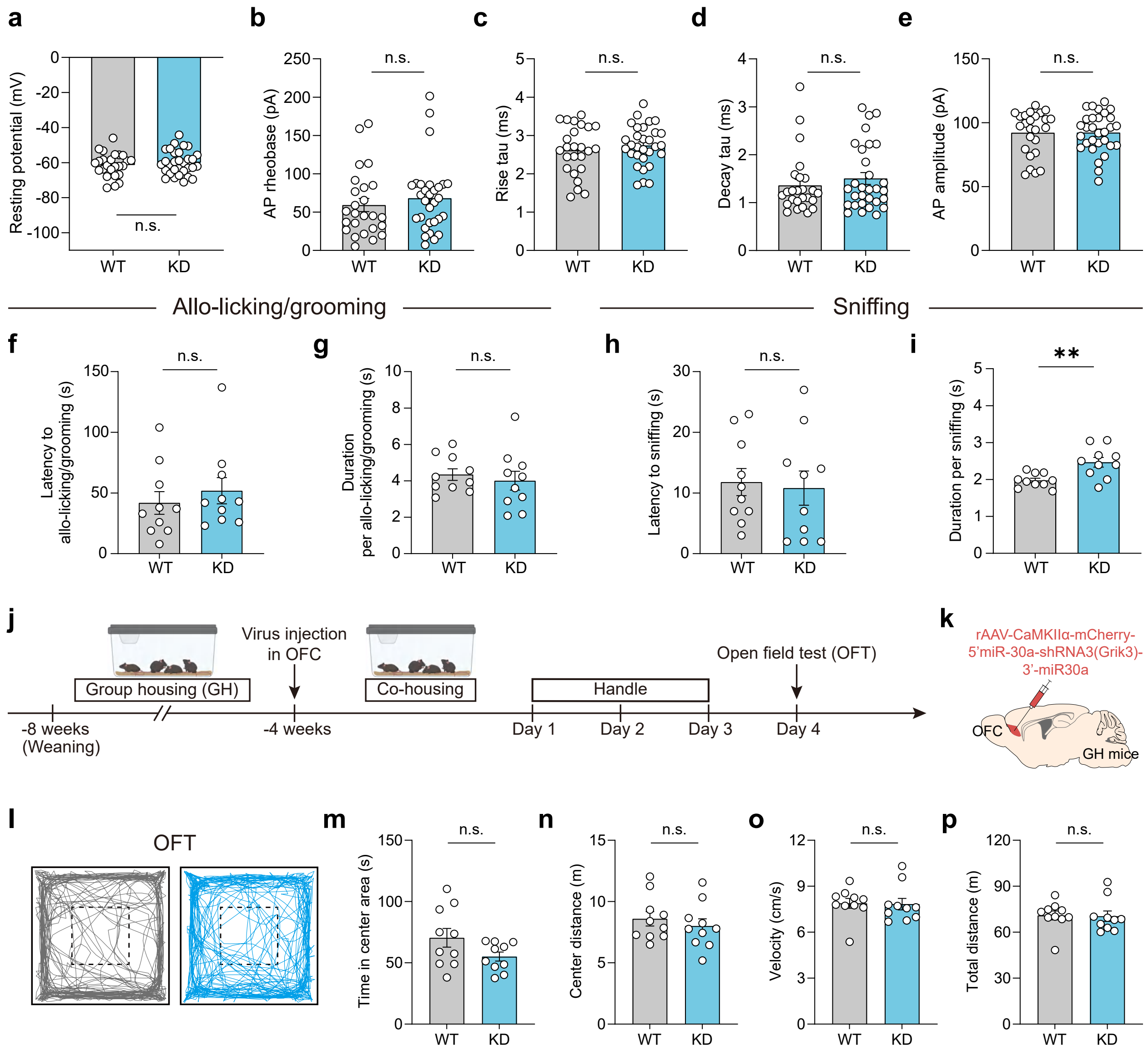
